# Phylogenomic Analysis of Target Enrichment and Transcriptome Data Uncovers Rapid Radiation and Extensive Hybridization in Slipper Orchid Genus *Cypripedium* L

**DOI:** 10.1101/2024.01.24.577114

**Authors:** Loudmila Jelinscaia Lagou, Gudrun Kadereit, Diego F. Morales-Briones

## Abstract

**Background and Aims:** *Cypripedium* is the most widespread and morphologically diverse genus of slipper orchids. Despite several published phylogenies, the topology and monophyly of its infrageneric taxa remained uncertain. Here, we aimed to reconstruct a robust section-level phylogeny of *Cypripedium* and explore its evolutionary history using target capture data for the first time.

**Methods:** We used the orchid-specific bait set Orchidaceae963 in combination with transcriptomic data to reconstruct the phylogeny of *Cypripedium* based on 913 nuclear loci, covering all 13 sections. Subsequently, we investigated discordance among nuclear and chloroplast trees, estimated divergence times and ancestral ranges, searched for anomaly zones, polytomies, and diversification rate shifts, and identified potential gene (genome) duplication and hybridization events.

**Key Results:** All sections were recovered as monophyletic, contrary to the two subsections within sect. *Cypripedium*. The two subclades within this section did not correspond to its subsections but matched the geographic distribution of their species. Additionally, we discovered high levels of discordance in the short backbone branches of the genus and within sect. *Cypripedium*, which can be attributed to hybridization events detected based on phylogenetic network analyses, and incomplete lineage sorting caused by rapid radiation. Our biogeographic analysis suggested a Neotropical origin of the genus during the Oligocene (∼30 Ma), with a lineage of potentially hybrid origin spreading to the Old World in the Early Miocene (∼22 Ma). The rapid radiation at the backbone likely occurred in Southeast Asia around the Middle Miocene Climatic Transition (∼15–13 Ma), followed by several independent dispersals back to the New World. Moreover, the Pliocene-Quaternary glacial cycles may have contributed to further speciation and reticulate evolution within *Cypripedium*.

**Conclusions:** Our study provided novel insights into the evolutionary history of *Cypripedium* based on high-throughput molecular data, shedding light on the dynamics of its distribution and diversity patterns from its origin to the present.

## INTRODUCTION

The family Orchidaceae comprises the most species-rich family of vascular plants, with c. 28,000 species in five subfamilies and ∼750 genera (Chase *et al.,* 2015; Christenhusz *et al.,* 2017). Their great diversity has fascinated and puzzled scientists for centuries, including the father of evolutionary theory, Charles Darwin, who once wrote, “I never was more interested in any subject in my life, than in this of Orchids” (to J. D. Hooker on 13 October 1861; Burkhardt *et al*., 1995). Unfortunately, today, their diversity is highly threatened mainly due to habitat destruction and unsustainable harvesting (DL Roberts and Dixon, 2008), prompting their protection by local and national laws, as well as the Convention on International Trade in Endangered Species (aka CITES; *Appendices I, II, and III*, 2023). In efforts to describe their diversity and facilitate informed conservation measures, a variety of molecular data, including high-throughput genomic and transcriptomic data, has been used to reveal the relationships between orchid subfamilies in recent decades (Cameron *et al.,* 1999; Freudenstein *et al.,* 2004; Givnish *et al.,* 2015; Kim *et al.,* 2020; Pérez-Escobar *et al.,* 2021; Serna-Sánchez *et al.,* 2021). However, phylogenetic support at lower taxonomic ranks in Orchidaceae has been low primarily due to the limited genetic variation in the commonly used markers (i.e., rbcL, matK, ITS, chloroplast intergenic spacers). The genus *Cypripedium* is one such orchid taxon whose internal phylogenetic relationships have yet to be resolved.

*Cypripedium* is a genus of temperate perennial herbs in the subfamily of lady’s slipper orchids, Cypripedioideae, and it currently consists of approximately 50 accepted species (Frosch and Cribb, 2012; SC Chen *et al*., 2013; Christenhusz *et al*., 2017; POWO, 2023). Although *Cypripedium* has only about half the species number of the largest slipper orchid genus, *Paphiopedilum*, it is the most morphologically diverse (Fig. 1) and widespread (Supplementary Data Fig. S1) of all five cypripedioid genera. Its distribution is mainly circumboreal, but its range extends from the Arctic Circle to Central America (∼14°–70° North; J Li *et al*., 2011; Frosch and Cribb, 2012). Eastern Asia, especially temperate China, constitutes the genus’ main center of diversity, harboring approximately 70% of all *Cypripedium* species (J Li *et al*., 2011). They occur in various habitats and altitudes, from forests to wetlands and grasslands and from sea level to 4,900 m in the Himalayas (Frosch and Cribb, 2012).

**Figure 1.**
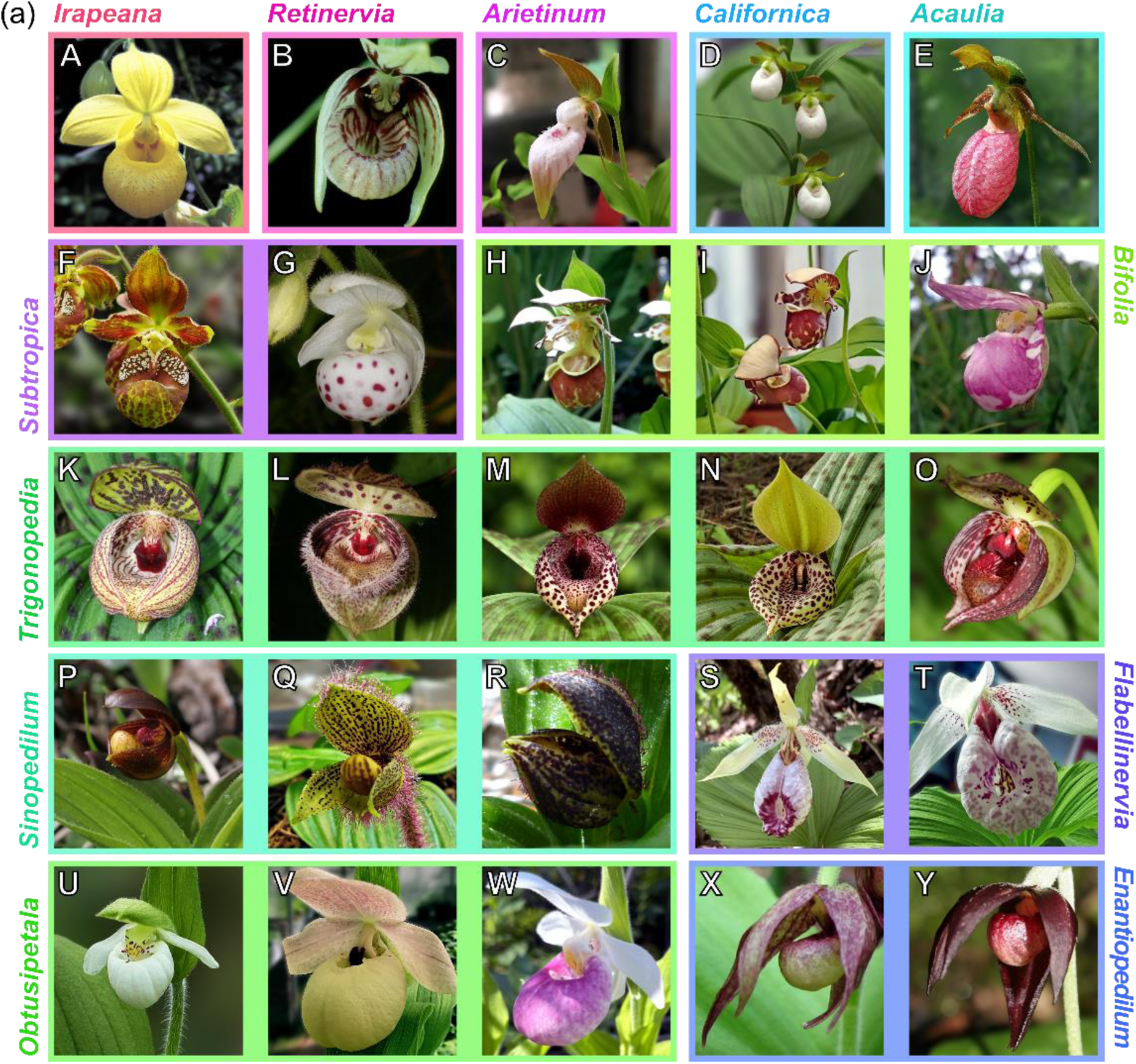
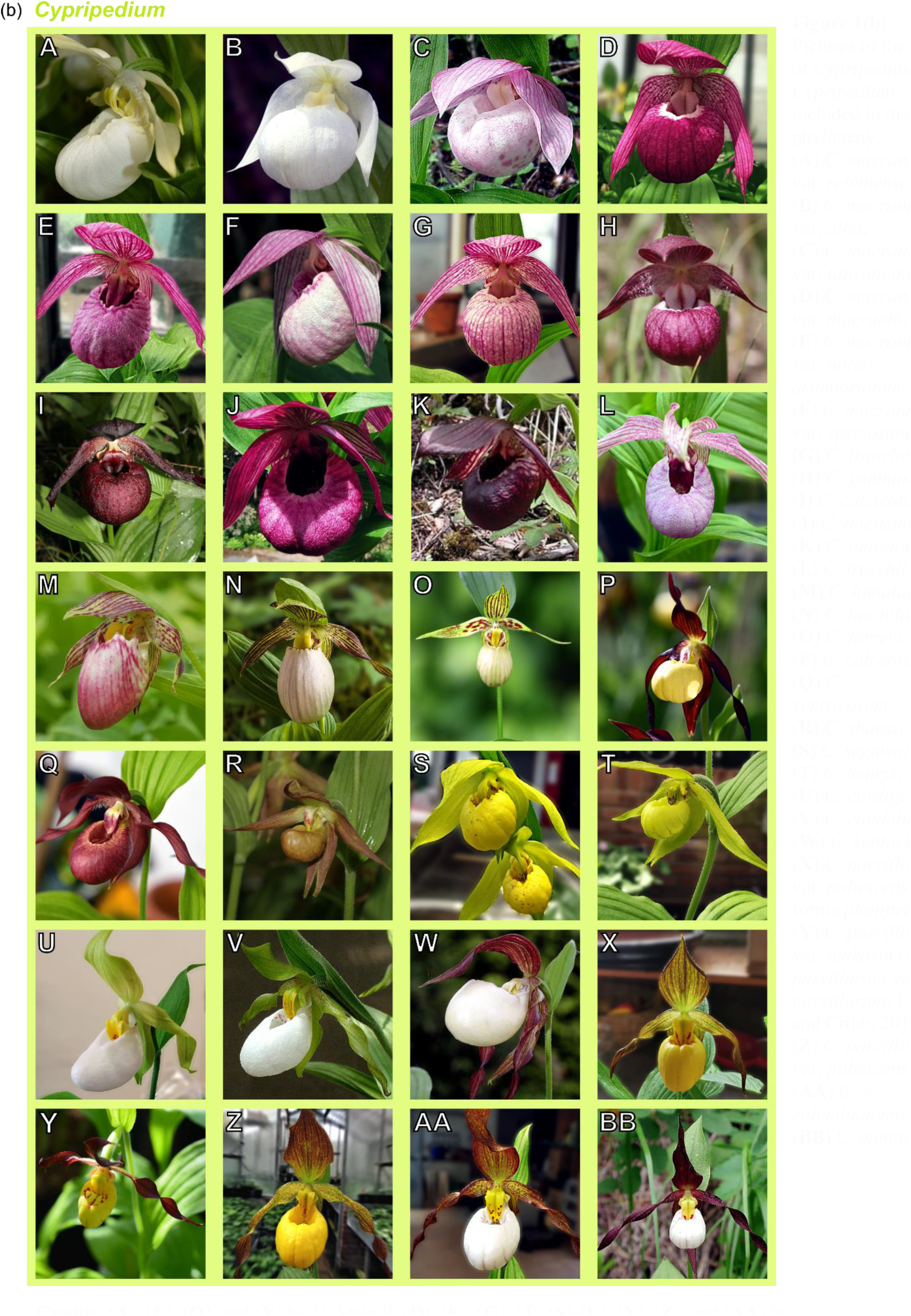
**(a):** Pictures of the *Cypripedium* taxa per section included in the final phylogeny. **(A)** *C. irapeanum*, **(B)** *C. debile*, **(C)** *C. plectrochilum*, **(D)** *C. californicum*, **(E)** *C. acaule*, **(F)** *C. subtropicum* (= *C. singchii*; Frosch and Cribb, 2012), **(G)** *C. wardii*, **(H)** *C. yatabeanum*, **(I)** *C.* × *alaskanum*, **(J)** *C. guttatum*, **(K)** *C. margaritaceum*, **(L)** *C. fargesii*, **(M)** *C. lichiangense*, **(N)** *C. lentiginosum*, **(O)** *C. sichuanense*, **(P)** *C. bardolphianum*, **(Q)** *C. micranthum*, **(R)** *C. forrestii*, **(S)** *C. japonicum*, **(T)** *C. formosanum*, **(U)** *C. passerinum*, **(V)** *C. flavum*, **(W)** *C. reginae*, **(X)** *C. fasciculatum*, **(Y)** *C. palangshanense*. **Credits**: (**A**) by M. Béhar; (**B**), (**D**), (**K**)-(**M**), (**O**), (**P**), (**R**) and (**Y**) by S. Urban; (**C**), (**V**), (**H**), (**I**), (**T**), and (**Q**) by J.-B. Chazalon; (**F**), (**G**), (**N**), (**U**), and (**E**) by W. Frosch; (**J**) and (**S**) by L. Chen; (**W**) by B. Isaac; (**X**) by the Forest **(b):** Pictures of the taxa of *Cypripedium* sect. *Cypripedium* included in the final phylogeny. **(A)** *C. macranthos* var. *rebunense*, **(B)** *C. macranthos* var. *alba*, **(C)** *C. macranthos* var. *taiwanianum,* **(D)** *C. macranthos* var. *macranthos*, **(E)** *C. macranthos* var. *hotei-atsumorianum*, **(F)** *C. macranthos* var. *speciosum*, **(G)** *C. franchetii*, **(H)** *C. yunnanense*, **(I)** *C. calcicola*, **(J)** *C. tibeticum*, **(K)** *C. amesianum*, **(L)** *C. froschii*, **(M)** *C. himalaicum*, **(N)** *C. fasciolatum*, **(O)** *C. farreri*, **(P)** *C. calceolus*, **(Q)** *C.* × *ventricosum*, **(R)** *C. shanxiense*, **(S)** *C. segawai,* **(T)** *C. henryi*, **(U)** *C. cordigerum*, **(V)** *C. candidum*, **(W)** *C. kentuckiense*, **(X)** *C. parviflorum* var. *pubescens* forma *planipetalum*, **(Y)** *C. parviflorum* var. *makasin* (= *C. parviflorum* var. *parviflorum*; Frosch and Cribb, 2012), **(Z)** *C. parviflorum* var. *pubescens*, (**AA**) *C.* × *columbianum*, (**BB**) *C. montanum*. **Credits**: (**A**), (**L**), (**Q**), and (**Y**) by V. Steindl; (**D**), (**E**), (**G**), (**J**), (**S**)-(**U**), (**X**), (**Z**), and (**AA**) by J.-B. Chazalon; (**P**) by L. Chen; (**W**) by Orchi; (**B**), (**F**), (**H**), (**I**), (**K**),(**M**), (**O**), and (**R**) by S. Urban; (**C**), (**N**), (**V**), and (**BB**) by W. Frosch (see Acknowledgements for more details).

Like other slipper orchids, flowers of *Cypripedium* species have a profoundly inflated, slipper-shaped lip (labellum) that gives them their distinctive morphology. The lip traps pollinators that enter through the upward-facing opening thanks to its incurved, glabrous, slippery margins, with the only escape routes passing through its basal orifices under the two anthers at each side of the column (Cribb, 1997; Frosch and Cribb, 2012). Unlike other slipper orchids, *Cypripedium* species are traditionally recognized by their (usually) plicate leaves and unilocular ovaries with parietal placentation (Cox *et al.,* 1997; Cribb, 1997). Although the reliability of these distinctive characters has been questioned (Atwood, 1984), phylogenetic studies consistently support the monophyly of the genus (Fatihah *et al*., 2011; J Li *et al*., 2011; Guo *et al*., 2012; H Liu *et al*., 2021*a*; Szlachetko *et al*., 2021; J-Y Zhang *et al*., 2022). On the other hand, its infrageneric classification has constantly changed during the last centuries.

Following *Cypripedium*’s description, the great interest in Cypripedioideae led to numerous taxonomic revisions in the subfamily with often incongruent results (Linnaeus, 1753; Rafinesque, 1836; Lindley, 1840; Reichenbach, 1854; Pfitzer, 1888, 1894; Rolfe, 1896; Atwood, 1984; Cox *et al*., 1997; Eccarius, 2009; Perner, 2008; Supplementary Data Table S1). To name a few, Linnaeus (1753) initially recognized only one species of *Cypripedium* (i.e., *C. calceolus* L.) and a few varieties currently holding a species status. Lindley (1840) described 22 species within the genus, classifying them into a number of subgeneric groups. The classifications of Pfitzer (1903) taxonomically expanded *Cypripedium* with 28 species and numerous subgeneric categories, including four sections. In Cribb’s (1997) taxonomic treatment, the number of species increased to 45 and the sections to 11, while Eccarius (2009) divided *Cypripedium* into two subgenera, 13 sections, and 37 species, lowering the rank of multiple species to subspecies or varieties.

Recent molecular phylogenies based on nrDNA ITS and five cpDNA markers by J Li *et al*. (2011) indicated that, among the non-monotypic groups, eight sections are monophyletic (*Arietinum, Bifolia, Cypripedium, Flabellinervia, Obtusipetala, Sinopedilum*, *Subtropica*, and *Trigonopedia*) while two sections (*Irapeana* and *Retinervia*) and the two subsections of sect. *Cypripedium* (*Cypripedium* and *Macrantha*) are non-monophyletic, following the classification by Cribb (1997) and Perner (2008). These results prompted further infrageneric treatments by Frosch and Cribb (2012) and SC Chen *et al*. (2013), producing the two currently used classification systems of *Cypripedium*. Although based on the same phylogenies by J Li et al. (2011), Frosch and Cribb (2012) proposed 13 sections with 48 species, whereas SC Chen *et al*. (2013) increased these numbers to 15 and 51, respectively, adding two new monotypic sections: *Palangshanensia* and *Wardiana* (Supplementary Data Table S1).

After the publication of J Li *et al*. (2011), several studies included molecular phylogenies with *Cypripedium* species, five of which investigated the relationships of the infrageneric taxa of *Cypripedium* in detail (Fatihah *et al*., 2011; Guo *et al*., 2012; Liao *et al*., 2024; H Liu *et al*., 2021*a*; Pérez-Escobar *et al*., 2023; Szlachetko *et al*., 2021; J-Y Zhang *et al*., 2022). These studies employed different phylogenetic reconstruction methodologies (i.e., Parsimony, Maximum Likelihood, and Bayesian Inference) with most using up to eight Sanger-sequenced nuclear and chloroplast DNA markers in different combinations, while the most recent study used plastome data and 41 nuclear loci derived from transcriptomes (Liao *et al*., 2024). The topologies and the monophyly of some subgeneric taxa were congruent among the produced phylogenies (e.g., sect. *Irapeana* being sister to the rest; monophyly of sect. *Arietinum, Bifolia, Cypripedium, Flabellinervia, Obtusipetala, Sinopedilum,* and *Trigonopedia*). However, the topology and monophyletic status of other taxa (e.g., the monophyly of the two subsections within sect. *Cypripedium*) and the topology at the backbone of the phylogeny remain uncertain.

The unresolved phylogeny of the genus *Cypripedium* not only prevents the accurate evaluation of the relationships between the currently established subgeneric groups but also our understanding of their evolutionary history. A well-resolved and robust phylogeny is fundamental for addressing further questions regarding their divergence time, diversification rate shifts, ancestral spatial distribution patterns, and hybridization events. Furthermore, it will provide a solid foundation for the efficient management of their conservation, especially as their continuous human-driven population decline is predicted to exacerbate due to climate change (Nicolè *et al*., 2005; Izawa *et al*., 2007; Minasiewicz *et al*., 2018; Kolanowska and Jakubska-Busse, 2020; H Liu *et al*., 2021*b*; Chandra *et al*., 2023; Yamashita *et al*., 2023).

It is widely recognized that the use of multiple genes can improve the accuracy of phylogenetic reconstruction, and single- or low-copy genes are increasingly used as they reduce the sequencing of paralogous genes (Guo *et al*., 2012; N Zhang *et al*., 2012; Z Li *et al*., 2017). Additionally, nuclear genes are often preferred for phylogenetic studies in plants due to generally higher evolutionary rates and biparental inheritance compared to plastid genes (Wolfe *et al*., 1987; Choi *et al*,. 2019). A target enrichment approach would allow the sequencing of hundreds of low-copy markers via high-throughput sequencing methods and, therefore, more robust estimates of relationships with greater support. Moreover, the use of the recently designed orchid-specific baits Orchidaceae963 by Eserman *et al*. (2021) could provide sufficient information to resolve recent and rapid radiations in deep and shallow phylogenetic scales, allowing for the characterization of species-level relationships and the resolution of long-debated polytomies within Orchidaceae.

In this study, we used a target enrichment approach with the Orchidaceae963 baits to reconstruct a well-supported phylogeny of the genus *Cypripedium* at the section level. Based on our results, we evaluated the two most recently published classification systems of the genus by Frosch and Cribb (2012) and SC Chen *et al*. (2013) and the congruence of the recovered relationships with published phylogenies based on Sanger data. Additionally, we aimed to gain new insights into the evolution of *Cypripedium* by answering the following questions: (1) Does the current classification stand up to phylogenetic reconstructions based on genomic data? (2) Which biological processes explain the increased levels of gene tree discordance in some parts of the phylogeny? (3) When and where did *Cypripedium* originate and diversify, and how did this diversification relate to the geographic expansion of the lineages and the paleoclimate? To answer these questions, we explored the discordance among gene trees, as well as between nuclear and chloroplast trees, estimated divergence times and ancestral ranges, searched for anomaly zones and diversification rate shifts, and identified gene duplication and hybridization events.

## MATERIALS AND METHODS

### Taxon Sampling for Target Enrichment

We sampled leaf tissue from 58 specimens representing 37 species, seven varieties, and three natural hybrids of the genus *Cypripedium* (following the taxonomy of Frosch and Cribb, 2012; Supplementary Data Table S2). Fifty-one of the sampled individuals came from the Botanical Collection at Oberhof, Eurasburg, associated with the Botanical Garden Munich-Nymphenburg (BGM), and seven from the Botanische Staatssammlung München herbarium (BSM-SNSB, herbarium acronym M). The material from the living collection was stored in silica-gel and dried immediately after collection. In addition, we included four DNA samples kindly provided by the DNA and Tissue Bank of the Kew Royal Botanical Gardens (Supplementary Data Table S3). The samples represented four different species, one of which (*C. fasciculatum* Kellogg ex S. Watson) was new to our sampling.

The sequence data from the above tissue and DNA samples was combined with publicly available orchid genomes (four), transcriptomes (20), and genome skimming libraries (two; Supplementary Data Table S4). These represented species from all slipper orchid genera (incl. 12 *Cypripedium* species, five of which were new to our sampling) as well as orchids from three outgroup subfamilies (i.e., Apostasioideae, Epidendroideae, and Vanilloideae). As a result, our final dataset included species from all 13 sections and subsections of the genus *Cypripedium* (following Frosch and Cribb, 2012).

### Library Preparation, Target Enrichment, and Sequencing

We isolated total genomic DNA from silica-dried or herbarium leaf tissue using the NucleoSpin Plant II kit: Genomic DNA from plants (Macherey-Nagel, Düren, Germany). DNA was sheared to an average fragment size of 350 bp with a Covaris M220 Focused-ultrasonicator (Covaris, Woburn, Massachusetts, USA). We prepared dual-indexed libraries according to the instruction manual using the NEBNext Ultra II DNA Library Prep Kit for Illumina and the NEBNext Multiplex Oligos for Illumina (Dual Index Primers Set 1, New England Biolabs, Ipswich, Massachusetts, USA). We used the custom orchid-specific bait set Orchidaceae963 (Daicel Arbor Biosciences myBaits Target Capture Kit, Ann Arbor, MI, USA) for the hybridization enrichment reaction (myBaits Hybridization Capture for Targeted NGS, User Manual v. 5.02). The enriched pooled libraries were sequenced on an Illumina NovaSeq 6000 (SP flow cell) or NextSeq 1000 (P1.300 flow cell) sequencing systems at the Core Facility Genomics (CF-GEN) of the Helmholtz Zentrum München, Germany (Deutsches Forschungszentrum für Gesundheit und Umwelt, GmbH). For further details regarding DNA extraction and sonication, library preparation, and target enrichment, refer to Supplementary Data Methods S1.

### Read Processing and Assembly

We created a set of references from orchid genomes and transcriptomes available on the Sequence Read Archive (SRA) of NCBI to improve gene extractions (Supplementary Data Table S4; Sayers *et al*., 2022). Next, we used CAPTUS v.1.0.0 (Ortiz *et al*., 2023) to trim the sequencing adaptors and low-quality bases, assemble the reads, and extract the nuclear loci based on the reference dataset. Similarly, we assembled and extracted the nuclear loci from the genomes and transcriptomes used as references to combine them with our data for further analysis. We then extracted the coding sequences from the combined dataset using CAPTUS and obtained nuclear loci FASTA files (for further information, refer to Supplementary Data Methods S2).

### Orthology Inference of nuclear loci

Orthology inference was performed following a modified version of the methods described in Morales-Briones *et al*. (2022; https://bitbucket.org/dfmoralesb/target_enrichment_orthology). Specifically, we used the tree-based “monophyletic outgroup” (MO) approach described in Yang and Smith (2014). The MO method searches for clusters with monophyletic ingroups rooted at the outgroups in the homolog trees, discarding those with duplicated taxa in the outgroups. Subsequently, it infers the orthologs from root to tip, keeping the ortholog subtree with the most taxa. To infer the orthologs, we set all Cypripedioideae members as ingroup, and the remaining taxa [i.e., *Apostasia shenzhenica* Z. J. Liu & L. J. Chen, *Dendrobium catenatum* Lindl., *Phalaenopsis equestris* (Schauer) Rchb.f., *Vanilla planifolia* Andrews, *Vanilla shenzhenica* Z. J. Liu & S. C. Chen] as outgroups, keeping 913 orthologs with at least 20 ingroup taxa. For further details regarding orthology inference, see Supplementary Data Methods S3.

### Phylogenetic Reconstruction

We used concatenation and coalescent-based methods to reconstruct the phylogeny of *Cypripedium*. First, a concatenated alignment of all 913 nuclear loci was produced using the clean ortholog alignments. We estimated an ML tree of the concatenated matrix with IQ-TREE v.2.2.6 (Minh *et al*., 2020). We searched for the best partition scheme using ModelFinder implemented within IQ-TREE (Kalyaanamoorthy *et al.,* 2017) and 1,000 ultrafast BS replicates to assess clade support. Regarding the coalescent-based approach, we first inferred ML trees from the same 913 individual orthologs used for the concatenation-based phylogeny. Individual ortholog ML trees were inferred as previously described for the final homolog trees. Then we used the quartet-based species-tree inference method ASTRAL v1.15.2.4 (wASTRAL-unweighted), which is statistically consistent under the multispecies coalescent (MSC) model and thus useful for handling incomplete lineage sorting (C Zhang *et al.,* 2018; C Zhang and Mirarab, 2022). We inferred the species tree using the 913 individual ML ortholog trees with default ASTRAL parameters and assessed branch support with local posterior probabilities (LPP; Sayyari and Mirarab, 2016). Due to the similarity in the topologies recovered between the concatenation and coalescent-based approaches, all subsequent analyses were carried out using the ASTRAL species tree unless stated otherwise.

### Gene Tree Discordance Estimation

We quantified the conflict among nuclear gene trees on each node of the inferred species tree by estimating the number of conflicting and concordant bipartitions with Phyparts (Smith *et al.,* 2015). To do this, we used the individual ML ortholog trees and set a threshold of at least 70% BS support for a node to be considered informative. We plotted the Phyparts result using the “missing and uninformative” script (i.e., “phypartspiecharts_missing_uninformative.py; https://bitbucket.org/dfmoralesb/target_enrichment_orthology) with Python v3.10.10 to add pie charts at the nodes while taking into consideration missing data (i.e., when input trees do not have the same number of tips).

We also used Quartet Sampling (QS; Pease *et al*., 2018) to differentiate between lack of support and conflicting nodes on the species tree. QS estimates branch support and conflict by sampling quartets from the species tree and the corresponding concatenated alignment and calculating the proportion of the three possible topologies at each node. As a result, it simultaneously evaluates the consistency of information (Quartet Concordance, QC), the presence of secondary evolutionary histories (Quartet Differential, QD), and the amount of information (Quartet Informativeness, QI) of internal nodes. We ran 1,000 QS replicates with RAxML-NG (Kozlov *et al.,* 2019) as the ML inference tool. The results were plotted using R (R Core Team, 2023) by color-coding the values of QC on each node and annotating them with the rest of the estimated values (https://bitbucket.org/yanglab/conflict-analysis/src/master/).

### Plastid CDS assembly and tree inference

We assembled cpDNA coding sequences (CDS) from off-target reads of the target enrichment (56 samples; Supplementary Data Tables S2 and S3) and transcriptome data (15 samples; Supplementary Data Table S4) with CAPTUS and its provided set of chloroplast proteins as references. Additionally, we used CAPTUS to extract cpDNA CDS from plastomes available in NCBI GenBank for 25 samples, including 15 additional representatives of species already present in our target enrichment or transcriptome data (Supplementary Data Table S4). For most non-*Cypripedium* taxa (except three) only data from NCBI GenBank plastomes was used. We wrote FASTA files for 80 cpDNA CDS with CAPTUS, keeping only the best contig. We aligned individual cpDNA CDS using MACSE v.2.0.7 (Ranwez *et al*., 2018) with the default parameters, replacing the “!” with gaps at frameshifts and removing aligned columns with >90% missing data with Phyx (‘pxclsq’ JW Brown *et al.,* 2017). Then, we concatenated all 80 CDS and estimated an ML tree with IQ-TREE as previously described for the concatenated nuclear loci. Additionally, we used QS with 1000 replicates to also evaluate potential conflict.

### Anomaly Zone Test

The anomaly zone, characterized by the presence of a set of short internal branches in the nuclear species tree, occurs when gene tree topologies that are discordant with the species tree topology are observed more frequently than those that are concordant (Linkem *et al.,* 2016). It arises from consecutive rapid diversification events leading to incomplete lineage sorting (ILS).

We estimated the boundaries of the anomaly zone for the internal nodes of our species tree following the calculations in Linkem *et al*. (2016) (https://github.com/cwlinkem/anomaly_zone) to investigate whether the high amount of gene tree discordance observed in numerous short branches of the tree could be explained by ILS. The calculations were based on equation 4 of Degnan and Rosenberg (2006), which defines the boundaries of the anomaly zone, α(*x*). In this equation, *x* is the length of an internal branch in the species tree, and its descendant internal branch has a length *y* (in coalescent units). If *y* is < α(*x*), then the internode pair is considered to be in the anomaly zone.

### Polytomy Test

Additionally, due to the presence of short branches with low support, we tested whether we could reject the null hypothesis that any branch in the nuclear species tree has a length equal to 0, or in other words, is a polytomy. We used the polytomy test (-t 10) option in ASTRAL version 5.7.8 with default parameters (Sayyari and Mirarab, 2018). The ASTRAL polytomy test relies on the distribution of the quartet frequencies of gene trees around each branch of the species tree to test this hypothesis, annotating the branches of the output tree with the resulting p-values. Under the null hypothesis, the three unrooted quartet topologies defined around the branch are expected to have equal frequencies. Although failure to reject the null hypothesis may indicate a real (i.e., hard) polytomy, it might also be caused by a lack of power or signal (i.e., soft polytomy).

### Mapping Gene Duplications

We mapped gene duplication events on our nuclear species tree based on the subclade orthogroup tree topology method described in Yang *et al*. (2018; https://bitbucket.org/blackrim/clustering/src/master/). This method extracts the rooted orthogroups from each homolog tree. Then, it detects gene duplication events when the orthogroup subclades share two or more ingroup taxa and maps the proportion of duplicated genes to the corresponding branch of the species tree. Alternatively, the duplications are mapped on the most recent common ancestor branch if the gene tree has missing taxa or if its topology is incongruent with the species tree. To avoid the overestimation of the duplication proportions due to nested duplications, each branch of the species tree is restricted to one duplication event for each extracted clade (Yang *et al.,* 2015). Similarly to Yang *et al*. (2018), we tested two filters to map the gene duplications: a bootstrap and a local topology filter. The bootstrap filter requires orthogroups to have an average bootstrap percentage of ≥70% (Z Li *et al.,* 2015), while the local topology filter requires the sister clade of the gene duplication branch in the orthogroup to include a subset of the taxa in the corresponding sister clade in the species tree (Cannon *et al.,* 2015). We plotted the proportions of gene duplications per number of branches in R.

To explore further potential whole genome duplication (WGD) events and compare them to gene duplication events identified from target enrichment data, we analyzed the distribution of synonymous distances (Ks) from RNA-seq data of 23 species of *Cypripedium*, six species of other slipper orchids, and three orchid outgroups. This included the sequencing of ten new transcriptomes of *Cypripedium* to cover all sections except Sect. *Retinervia* (Supplementary Data Table S6). Total RNA was extracted from leaves or shoot buds using the innuPREP Plant RNA Kit (Innovative Sensor Technology; Ebnat-Kappel, Switzerland). Library preparation and sequencing on an Illumina NovaSeq X Plus platform were performed at Novogene Co. (Cambridge, United Kingdom). Read processing and transcriptome assembly followed Morales-Briones *et al*. (2021). For each species, a Ks plot of within-species paralog (whole paranome) was done using *wgd* v2.0.31 (*dmd* and *ksd* programs with default parameters; H Chen *et al*., 2024) Additionally, we used Ks plots of between-species (reciprocal best hits) to establish the relative timing of the split between species and compare it to WGD events inferred with within-species Ks plots. Between-species Ks plots were carried out with *wgd* between *Cypripedium* species, and between *Cypripedium* and other slipper orchids and orchid outgroups.

### Testing for Hybridization Events

Due to high discordance at the backbone of the phylogeny, we tested whether our data indicates any hybridization events between taxa of different sections of *Cypripedium* using explicit phylogenetic networks in PhyloNet (Wen *et al*., 2018). PhyloNet allows for horizontal edges that visualize the genetic inheritance through gene flow, mapping the inheritance probabilities (γ) for each parent hybrid edge to estimate the proportion of loci a hybrid inherited from each parent.

To reconstruct the phylogenetic networks, we first rooted the final ortholog trees and extracted subclades containing at least one representative taxon from each section and subsection, favoring the taxa present in most orthologs to maximize the final number of loci used for PhyloNet. Similarly, *Dendrobium catanetum* was chosen as an outgroup taxon because it had the highest amount of retained loci. Gene trees missing any of these selected taxa were excluded from the analysis. Since calculating the likelihood of a phylogenetic network is computationally intensive in PhyloNet, we inferred the phylogenetic networks based on a maximum pseudo-likelihood (MPL) measure via the InferNetwork_MPL command (Yu and Nakhleh, 2015). We set the number of maximum reticulation events from one to ten and the number of optimal output networks to ten.

As a secondary objective, since our sampling included three taxa described as hybrids by both Frosch and Cribb (2012) and SC Chen *et al*. (2013), namely, *C.* × *alaskanum* P. M. Br. (*C. guttatum* Sw. × *C. yatabeanum* Makino), *C.* × *columbianum* Sheviak [*C. montanum* Douglas ex Lindl. × *C. parviflorum* Salisb. var*. pubescens* (Willd.) O. W. Knight] and *C.* × *ventricosum* Sw. (*C. calceolus* × *C. macranthos* Sw.), we tested whether our data supports their status as hybrids of their putative parent taxa (see Supplementary Data Methods S4 for more information). Moreover, we ran further tests setting the number of maximum reticulation events from two to ten for the networks containing *C.* × *columbianum* and *C.* × *ventricosum* to investigate whether there is support for more than one hybridization event in each case.

The phylogenetic networks with the highest total log probability between networks testing for up to one hybridization event and between all networks overall were visualized in Dendroscope v.3.8.8 (Huson and Scornavacca, 2012). The inheritance probabilities were mapped with PhyloNetworks’ v.0.16.2 (Solís-Lemus *et al.,* 2017) companion package PhyloPlots v.1.0.0 (https://github.com/cecileane/PhyloPlots.jl) in Julia v.1.9.2 (Bezanson *et al.,* 2012).

### Divergence Time Estimation

We used a Bayesian Inference approach for divergence time estimation. To decrease computational resources, we reduced the volume of the datasets to a subset of genes providing the most useful information relevant to time calibration via a “gene shopping” method. Specifically, we used the SortaDate package developed by Smith *et al*. (2018) to filter the 20 best ortholog genes based on the (a) least topological conflict with a focal species tree (i.e., bipartition calculation), (b) clock-likeness (i.e., root-to-tip variance statistic calculation), and (c) discernible information content (i.e., total tree length), sorting the genes in the respective order of these properties (i.e., a, b, c).

We concatenated the resulting subset of genes and defined the positions of the 20 loci as data blocks to find the best partitioning schemes and models of nucleotide evolution with PartitionFinder 2 (Lanfear *et al.,* 2017) to inform our site model selection for the molecular calibration. The branch lengths were estimated independently for each subset (i.e., unlinked), AICc was used to select the best-fit nucleotide substitution models among those available in BEAST v2.7.4 (Bouckaert *et al.,* 2019), while the “greedy” algorithm was used to search for a good partitioning scheme. The results suggested 16 partitioning sets, with GTR+I+G4+X as the best-fit model for 15 sets and GTR+G4+X as the best-fit model for the remaining one. For this reason and to reduce computational time, we decided to carry out molecular dating with the nucleotide substitution model GTR+I+G4+X without partitioning the selected loci.

Additionally, we used a relaxed uncorrelated lognormal clock model (Optimized Relaxed Clock, ORC; Mean clock rate: 1.0) and the topology of the ASTRAL nuclear phylogeny as a starting tree for 100 million generations, sampling every 10,000 generations. Since no available fossil of slipper orchids can be used for calibration and our sampling of orchids apart from Cypripedioideae is scarce, we used a secondary calibration point based on the age estimates by Givnish *et al*. (2015). In this study, the authors reconstructed a broad-scale phylogeny with species representing all orchid subfamilies with 75 plastid genes and calibrated against 17 angiosperm fossils using BEAST v. 1.8.0 (Drummond *et al.,* 2012). Based on their estimates, we set a normal distribution for the crown age of Orchidaceae (mean = 89.46, sigma = 1) and the crown age of Cypripedioideae (mean = 31.3, sigma = 1), and used a Birth-Death tree model (Gernhard, 2008). The rest of the priors were not modified from their default values. Four identical runs with distinct seed numbers were performed simultaneously to determine whether they converged on the same stationary distribution. A fifth run was performed sampling from the prior to examine whether the results were significantly skewed by the prior assumptions or informed by our data.

Convergence of the Markov Chain Monte Carlo (MCMC) chains was examined with Tracer v1.7.2 (Rambaut *et al.,* 2018) by checking that the Effective Sample Size (ESS) of the combined runs was >200 for all trace statistics and that the trace plots of the individual runs converged on the same posterior distribution. The tree files from the four independent runs were combined after removing 10% as burn-in using LogCombiner v1.8.2, and the maximum clade credibility chronogram was reconstructed using TreeAnnotator v1.8.2 with maximum clade median node height and 95% highest posterior density (HPD) intervals.

### Detection of Diversification Rate Shifts

We investigated if diversification rates changed throughout the evolutionary history of *Cypripedium* and whether there were significant rate shifts. To achieve this, we used BAMM v.2.5 (Rabosky *et al.,* 2013), a program developed to model the dynamics of speciation and extinction on phylogenetic trees. It considers time-dependent (e.g., a lineage’s age) and diversity-dependent (e.g., the number of lineages in a clade) effects to quantify diversification rates using a reversible-jump MCMC approach.

For the input tree, we modified the time-calibrated maximum clade credibility tree obtained from the divergence time estimation analysis by removing all non-*Cypripedium* species, as well as taxon duplicates, hybrids, and varieties, to avoid inflating the diversification rates. To account for non-random taxon sampling between the included *Cypripedium* sections, section-specific sampling fractions were calculated based on the classification by Frosch and Cribb (2012). The expected number of shifts was set to one, following the recommendation for small trees with less than 500 tips. The priors on the initial lambda, the lambda shift parameter, and the time mode for the speciation rate were calculated with the R package BAMMtools v2.1.10 (lambdaInitPrior = muInitPrior = 1.44786543849854, lambdaShiftPrior = 0.0522358069615021; Rabosky *et al*., 2014), the segment length (segLength) was set to 0.1, and the rest of the parameters were left as default. We ran four MCMC chains for 50 million generations and sampled every 1,000 generations. Subsequently, we used BAMMtools to check whether the MCMC runs converged (ESS >200) and discarded the first 25% of samples as burn-in. Then, we plotted the maximum a posteriori probability shift configuration (aka best shift configuration) as well as the speciation, extinction, and net diversification rates through time.

### Ancestral Range Estimation

We used the R package BioGeoBEARS (Matzke, 2013) to infer the biogeographic history of *Cypripedium*. BioGeoBEARS reconstructs the ancestral geographic distributions on phylogenies while testing for the best-fit model of range evolution. It replicates the basic assumptions of three widely used models in historical biogeography: DEC (Dispersal-Extinction-Cladogenesis; Ree and Smith, 2008), DIVA (Dispersal-Vicariance Analysis; Ronquist, 1997) and BayArea (Bayesian Inference of Historical Biogeography for Discrete Areas; Landis *et al*., 2013), implementing them in a Maximum Likelihood framework to allow for direct comparison. Together, these models allow for a broad range of processes, such as vicariance, sympatric speciation, range expansion, and contraction. They can also be combined with a founder-event (“jump”) speciation model specified with the parameter “j” (Matzke, 2014), the use of which has received some criticism due to conceptual and statistical issues (Ree and Sanmartín, 2018).

To infer the ancestral range estimation in *Cypripedium*, we conducted two analyses using different sets of areas for comparison. For our first analysis, we decided on nine areas based on the current distribution of the taxa, their proximity, and their distinct floristic and topoclimatic characteristics (see Supplementary Data Methods S5 for details), allowing up to five areas to be combined in an ancestral range. A distance matrix was also used to adjust the dispersal probabilities. The matrix included the distances between the closest points at the perimeters of every area pair combination in kilometers, measured in Google Maps. When two areas were adjacent, we set the distance between them to 1 km as recommended by the guidelines. The second analysis included only two specified areas, the Old and the New World, setting the maximum number of areas to two, without including a distance matrix.

For both analyses, we used a modified version of the time-calibrated maximum clade credibility tree obtained from the divergence time estimation analysis as our input tree (see Supplementary Data Methods S6 for details). We performed the analyses with all six biogeographical models (i.e., DEC, DEC+J, DIVALIKE, DIVALIKE+J, BAYAREALIKE, and BAYAREALIKE+J) but since the use of the “j” parameter has been disputed (Ree and Sanmartín, 2018) we plotted the best-fit models that did not implement the “j” parameter on the maximum clade credibility tree.

## RESULTS

### Assembly and Orthology Inference

The number of total extracted loci sequenced with target enrichment per *Cypripedium* species (with ≥75% identity and ≥50% coverage) ranged from 69 (*C. debile* Rchb.f., sample No. 62) to 846 (*C. fasciolatum* Franch., sample No. 84) for herbarium or old silica-dried material, and from 557 (*C. lichiangense* S. C. Chen & P. J. Cribb., sample No. 14) to 907 (*C. irapeanum* La Llave & Lex., sample No. 14) for recently collected silica-dried samples out of the 950 loci from the extended reference dataset. Overall, ∼760 loci were recovered with target enrichment on average, which is considerably higher than the corresponding proportion of loci reported for slipper orchids in the original publication by Eserman *et al*. [2021; i.e., 430 loci for *Paphiopedilum exul* (Ridl.) Rolfe and 533 loci for *Phragmipedium longifolium* (Warsz. & Rchb. f.) Rolfe]. Paralogs were found in all samples, from 5 (*C. debile*, sample No. 62) to 1,479 (*C. bardolphianum* W. W. Sm. & Farrer, sample No. 83) for herbarium or old silica-dried material, and from 55 (*C. irapeanum*, sample No. 14) to 1,823 (*C. micranthum* Franch., sample No. 22) for recently collected silica-dried samples, with ∼738 paralogs per sample on average for all loci combined. The orthology inference resulted in 913 MO orthologs (18–900 ortholog trees per taxon, ∼649 on average; Supplementary Data Table S7), producing a concatenated alignment with a length of 978,760 bp and a character occupancy of 66% for ≥ 20 ingroup taxa. In total, 77 *Cypripedium* specimens representing 58 taxa, among which 42 out of 48 accepted species from all 13 sections (following Frosch and Cribb, 2012) were included in the final dataset of nuclear orthologs (Fig. 1). Regarding the chloroplast orthologs, recovered CDS per sample ranged from 36 to 80 with an average of 72 (Supplementary Data Table S8). The final cpDNA alignment included 96 taxa and 68,823 bp with a character occupancy of 87%.

### Inferred Nuclear Species Phylogeny and Discordance

Both concatenation and coalescent-based phylogenetic analyses based on nuclear target enrichment data recovered the genus *Cypripedium* as monophyletic with maximum support (i.e., BS = 100, LPP =1; Fig. 2; Supplementary Data Fig. S2). Additionally, both employed gene tree concordance analysis approaches for the ASTRAL species tree showed high concordance for the most recent common ancestor (MRCA) of *Cypripedium*, with Phyparts identifying 638 informative concordant genes out of 677 and QS giving full support (i.e., 1.0/–/1.0), indicating that all sampled quartet replicates support the node (Supplementary Data Figs S3 and S4). The phylogenetic relationships between the slipper orchid genera were congruent between the ASTRAL and IQ-TREE trees, with *Cypripedium* being the sister to the rest. Within its sister clade, the plicate-leaved genus *Selenipedium* was recovered as sister to the clade of the conduplicate-leaved genera *Mexipedium*, *Phragmipedium*, and *Paphiopedilum*, with the two New World genera *Mexipedium* and *Phragmipedium* more closely related to each other than to the Old World *Paphiopedilum*.

**Figure 2:**
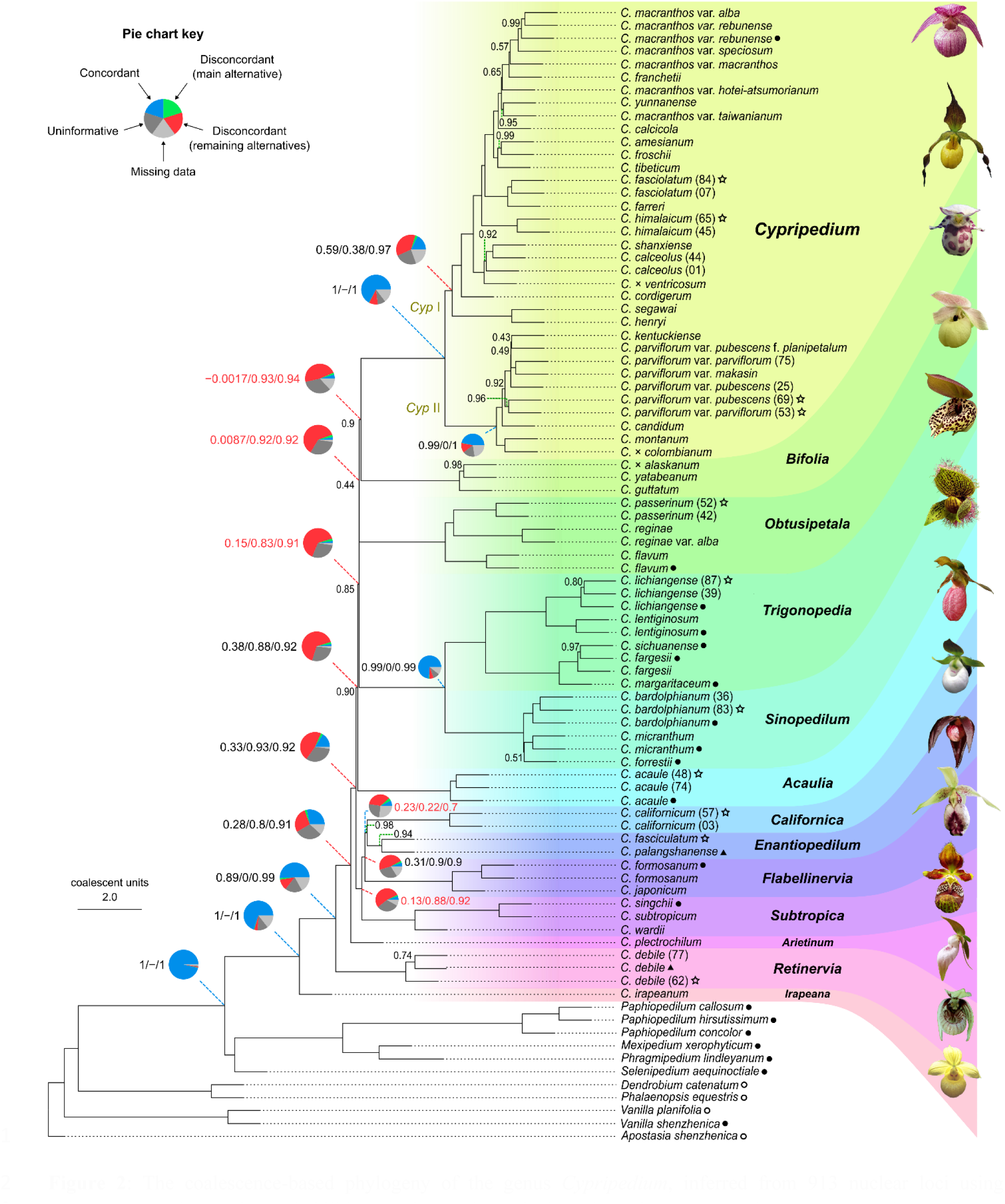
The coalescence-based phylogeny of the genus *Cypripedium*, inferred from 913 nuclear loci using ASTRAL. Local posterior probabilities are shown above the branches when <1. Branches are annotated according to their section-level classification following Frosch and Cribb (2012). Quartet Sampling (QS) support results (i.e., Quartet Concordance/Quartet Differential/Quartet Informativeness, in the same order) are shown to the left of the Phyparts piecharts for the backbone nodes at the MRCA between the sections, as well as the nodes leading to the two large clades within sect. *Cypripedium*, “*Cyp* I” and “*Cyp* II”. QS support values with Quartet Concordance < 0.25 are shown in red. Colors in the pie charts: blue denotes the proportion of concordant gene tree topologies, green denotes the proportion of gene trees with the main alternative topology, red denotes the proportion of gene trees with the remaining discordant topologies, light grey denotes the proportion of gene trees with missing taxa, dark grey denotes the proportion of uninformative gene trees. Tip symbols: filled circles “⬤” denote transcriptomes, unfilled circles “〇” denote genomes, filled triangles “▲” denote genome skimming sequences, and unfilled stars “☆” denote herbarium or old silica-dried specimens. Tips without symbols come from living specimens of the Botanical Collection at Oberhof. Flowers of representative species from each section are displayed to the right. Species names corresponding to the flower pictures, from top to bottom: *C. franchetii, C. parviflorum* var. *pubescens, C. guttatum, C. flavum, C. lentiginosum, C. micranthum, C. acaule, C. californicum, C. palangshanense*, *C. japonicum, C. subtropicum, C. plectrochilum*, *C. debile*, and *C. irapeanum*. See the legends of Figure 1 and the Acknowledgements section for the credits of the flower pictures.

All 13 sections were monophyletic within *Cypripedium*, but the subsections *Cypripedium* and *Macrantha* within sect. *Cypripedium* were non-monophyletic based on the classification of Frosch and Cribb (2012). However, when considering the classification of SC Chen *et al*. (2013), subsect. *Macrantha* was monophyletic whereas subsect. *Cypripedium* was paraphyletic, and these results were consistent between the two phylogenies. Although the species grouped in the two largest sister clades within sect. *Cypripedium* (clades I and II) did not correspond to the species compositions of its two subsections, they matched the distribution of the species within them, with clade I only found in the New World and clade II in the Old World (Supplementary Data Fig. S1).

Maximum support was recovered for most branches in both inferred phylogenies, except for some branches along the backbone, the MRCA of sect. *Enantiopedilum*, the MRCA of the (*Californica*, *Enantiopedilum*) clade, and within the sections *Retinervia, Trigonopedia*, *Sinopedilum*, *Bifolia*, and *Cypripedium*. Among the topologies with maximum support, high concordance, and congruence between the two phylogenies was the placement of the Mesoamerican sect. *Irapeana* being sister to the rest (LPP = 1; BS = 100; Phyparts: 638/677; QS: 1/–/1), followed by sect. *Retinervia* (LPP = 1, BS =100, Phyparts: 464/589, QS: 0.89/0/0.99). Additionally, sect. *Subtropica*, following the classification by Frosch and Cribb (2012), was recovered as monophyletic (LPP = 1; BS = 100; Phyparts: 509/580; QS: 0.84/0.97/0.97), including both sect. *Wardiana* and sect. *Subtropica* as described by SC Chen *et al*. (2013). The two fly-pollinated sections *Sinopedilum* and *Trigonopedia* were also supported as most closely related to each other, which was congruent between the gene trees and between the ASTRAL and IQ-TREE trees (LPP = 1; BS = 100; Phyparts: 674/735; QS: 0.99/–/0.99).

Nevertheless, the remaining inter-sectional relationships showed decreased gene tree concordance based on Phyparts and QS. For instance, although both inferred phylogenies supported the sister relationship between sect. *Subtropica* and the clade containing the sections *Flabellinervia*, *Californica*, and *Enantiopedilum* (LPP = 1; BS = 100), both gene tree concordance analyses indicated elevated levels of discordance for these nodes (Phyparts: 48/555; QS: 0.13/0.88/0.92). A similar pattern was observed for the inter-sectional MRCAs within the latter clade. The gene tree disagreement regarding the rest of the section-level relationships is largely attributed to the five short-length backbone branches (Fig. 3 B, branches 4 to 8; Supplementary Data Fig. S7), with the proportion of informative concordant genes estimated by Phyparts falling between 3.2–48%, and QS indicating either weak or counter support for these branches.

**Figure 3:**
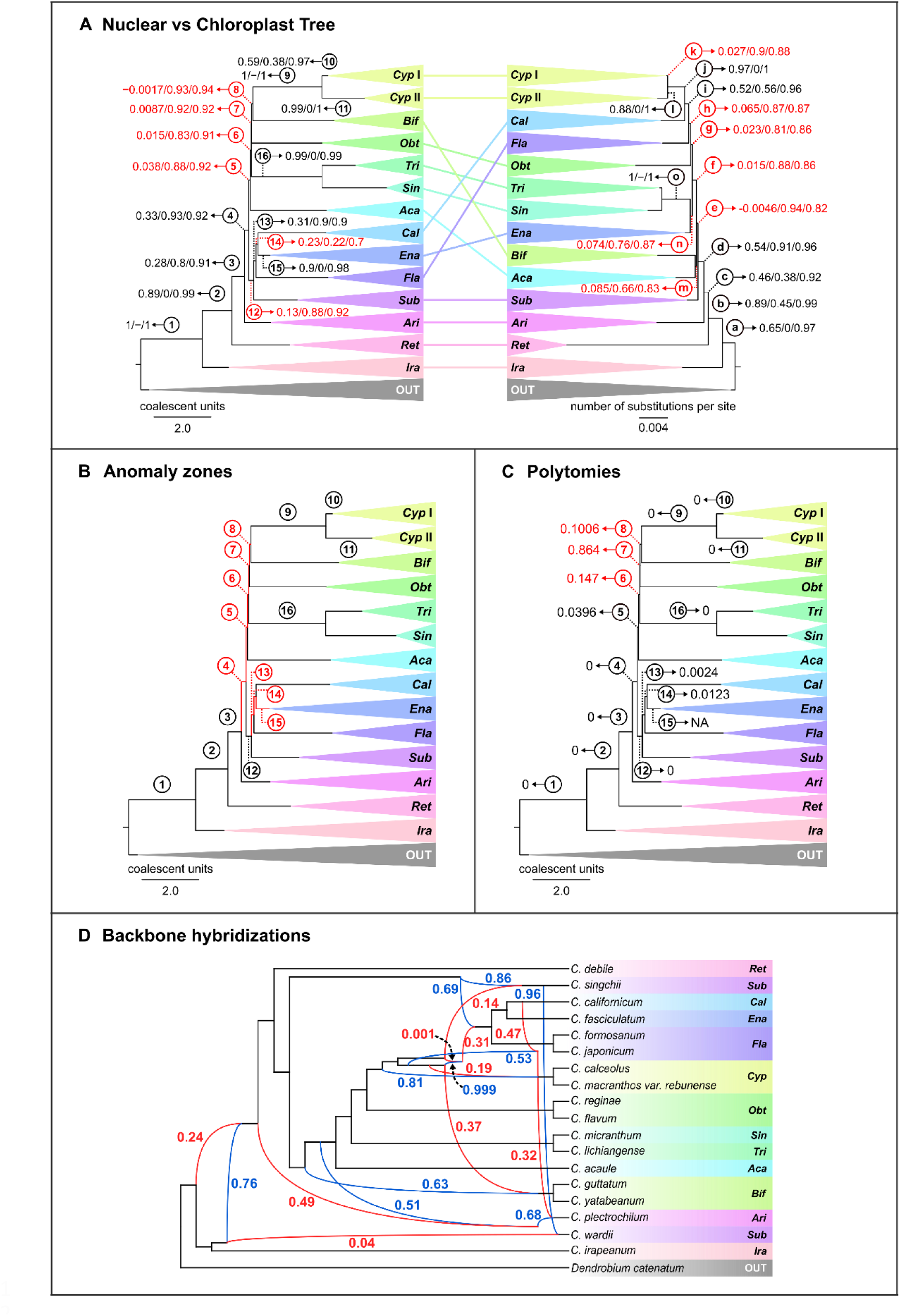
(**A**) Comparison of the section-level topologies between the (right) ASTRAL phylogeny based on nuclear loci and the (left) IQ-TREE phylogeny based on chloroplast loci, including Quartet Sampling support annotations (i.e., Quartet Concordance/Quartet Differential/Quartet Informativeness in order; for interpretation, see Pease et al., 2018]. QS support values with Quartet Concordance < 0.25 are shown in red. (**B**) Anomaly zone test: Branches of the collapsed ASTRAL target enrichment species tree shown in red are in the anomaly zone. (**C**) Polytomy test: Resulting p-values are annotated on the backbone branches of the collapsed ASTRAL target enrichment species tree. Polytomies based on α = 0.05 are shown in red. (**D**) Phylogenetic network with the highest total log probabilities resulting from the PhyloNet analysis testing for ten hybridization events using extracted subclades of all *Cypripedium* sections to test for reticulation in the backbone of the phylogeny. The inheritance probabilities are shown for each parent hybrid edge (blue = major hybrid edge; red = minor hybrid edge). Branch numbers are shown in circles. Key to collapsed clades: *Cyp =* sect. *Cypripedium, Cyp* I = clade I of sect. *Cypripedium* (see Fig. 2); *Cyp* II = clade II of sect. *Cypripedium* (see Fig. 2); *Bif = Bifolia*; *Obt* = *Obtusipetala*; *Tri* = *Trigonopedia*; *Sin* = *Sinopedilum*; *Aca* = *Acaulia*; *Cal* = *Californica*; *Ena = Enantiopedilum; Fla* = *Flabellinervia*; *Sub* = *Subtropica*; *Ari = Arietinum; Ret = Retinervia; Ira = Irapeana*. OUT = Outgroups.

Gene tree heterogeneity was mostly higher for nodes within sections than between sections, except for the sections *Cypripedium*, *Sinopedilum,* and some nodes within *Trigonopedia*, while sections *Retinervia* and *Enantiopedilum* showed a high degree of missing data. The latter was caused due to the single specimen of *C. palangshanense* T. Tang & F. T. Wang and two out of three specimens of *C. debile* being present in a small fraction of the total gene trees (Supplementary Data Table S7). However, sections *Retinervia* and *Enantiopedilum* were highly supported as monophyletic (LPP = 1, BS = 100, and LPP = 0.94, BS = 95, respectively), and the available informative gene trees for the respective nodes showed relatively low discordance (Phyparts: 49/58, QS: 1/–/1, and Phyparts: 5/12, QS: 0.9/0/0.98, respectively). On the other hand, sect. *Cypripedium* was highly supported as monophyletic with high concordance, but the number of informative concordant genes was lower than the number of discordant genes for almost all nodes within it. Only two nodes within this clade showed substantial concordance; namely, the (*C. henryi* Rolfe, *C. segawai* Masam.) clade (LPP = 1; BS =100; Phyparts: 355/387; QS: 1/–/1) and the node corresponding to the MRCA of clade II within the section (LPP = 1; BS =100; Phyparts: 427/554; QS: 0.99/0/1).

Regarding the phylogenetic relationships at the species level, most species were recovered as monophyletic, with some notable exceptions, such as *C. macranthos* and *C. parviflorum* Salisb, which were paraphyletic. For further details regarding the species-level phylogenetic results, refer to Supplementary Data Results S1.

### Comparison of Chloroplast and Nuclear Trees

The concatenation-based chloroplast phylogeny of *Cypripedium* inferred the same relationships between slipper orchid genera as the phylogenies based on nuclear data (Supplementary Data Fig. S5). The bootstrap support was generally high (i.e., BS > 80) for most branches, except for some topologies within sections *Bifolia, Sinopedilum,* and *Cypripedium*. However, the section-level topologies of the chloroplast tree were incongruent with the nuclear tree (Fig. 3 A). For instance, although sect. *Subtropica* was recovered as sister to the (*Flabellinervia* (*Californica*, *Enantiopedilum*)) clade in the nuclear tree, it was placed as sister to the rest following sections *Irapeana*, *Retinervia*, and *Arietinum* in the chloroplast tree. Furthermore, in the chloroplast tree, sect. *Enantiopedilum* was most closely related to the fly-pollinated clade of (*Sinopedilum*, *Trigonopedia*). Sections *Flabellinervia* and *Californica*, on the other hand, diverged consecutively with sect. *Californica* instead of *Bifolia* sharing an MRCA with sect. *Cypripedium*, while sect. *Bifolia* was recovered as sister to sect. *Acaulia* following the split of sect. *Subtropica.* Similarly to the nuclear tree, the QS analysis of the chloroplast tree also indicated either weak or counter-support for most branches at the backbone of the *Cypripedium* phylogeny (Fig. 3A; Supplementary Data Fig. S6).

### Anomaly Zone and Polytomy Test

The anomaly zone boundary estimations detected four internode pairs at the backbone of the nuclear *Cypripedium* species tree that are in the anomaly zone [i.e., y < α(x); Fig. 3 B, pairs 4-5, 5-, 6-7, 7-8, with α(x) equal to ∼0.1409, ∼0.6657, ∼0.7637, and ∼1.9122, respectively], as well as the internode pair between the MRCAs of the (*Flabellinervia* (*Californica*, *Enantiopedilum*)) and (*Californica*, *Enantiopedilum*) clades [Fig. 3 B, pair: 13-14 with α(x) = ∼0.3597] and the internode pair between the MRCAs of the (*Californica*, *Enantiopedilum*) clade and the section *Enantiopedilum* [Fig. 3 B, pair: 14-15, with α(x) = ∼ 0.4566]. Additionally, one internode pair between the MRCAs of *C. micranthum* and *C. forrestii* P. J. Cribb within sect. *Sinopedilum*, as well as several internode pairs within the two subclades of sect. *Cypripedium* also fell into the anomaly zone (see Supplementary Data Table S9 and Fig. S7 for further details).

The ASTRAL polytomy test failed to reject the null hypothesis that the branch length is 0 (i.e., p-value > α; α = 0.05) for branches 6, 7, and 8 (Fig. 3 C). These branches were also found to be in the anomaly zone, while branch 7 also received a low LPP support (LPP = 0.44). Moreover, branches within the subclades I and II of sect. *Cypripedium* found in the anomaly zone were also identified as polytomies (Supplementary Data Fig. S8).

### Gene Duplication Events

We mapped nuclear target enrichment orthogroups to detect gene duplications on the ASTRAL species tree using two filtering methods (i.e., bootstrap and local topology filters; Supplementary Data Figs S9 and S10). Most proportions of duplicated genes ranged from 0.01 to 0.18 using both filtering approaches, except for an outlier proportion of 0.29, which was identified by the analysis implementing the bootstrap filter on the MRCA of sect. *Cypripedium* (corresponding to 0.18 in the analysis using the local topology filter), possibly suggesting a WGD event. Nonetheless, the results of the Ks plots analysis failed to identify a WGD event within *Cypripedium* (Supplementary Data Fig. S11). Aside from the peak corresponding to the ancestral whole genome triplication shared by core eudicots (Ks = ∼2; Jiao *et al.,* 2012), the only other peak that the Ks plots detected was at around Ks = 0.4. This peak was found in all Cypripedioideae and outgroups orchids, thus corresponding to a WGD shared by a common ancestor of all Orchidaceae (G-Q Zhang *et al*., 2017; Supplementary Data Fig. S12).

### Hybridization Networks

In our PhyloNet analysis, we tested for one to ten reticulation events within a subclade containing representative taxa from each section and subsection of the genus to check whether it could provide support for backbone hybridization events and explain the observed discordance at these nodes of the nuclear species tree as well as between the nuclear and chloroplast trees. The network with the overall highest total log probability (∼ −138,385.32; Supplementary Data Fig. S13 C) indicated 10 hybridization events predominantly between unsampled or extinct taxa at the backbone of the phylogeny, as well as within sect. *Subtropica*, with varying inheritance probabilities (i.e., minor edge γ values ranging from 0.001 to 0.49; Fig. 3 D). In detail, the network indicated that an unsampled taxon constituting the sister of *C. irapeanum* (γ = 0.76) hybridized with an unsampled taxon closely related to their clade (γ = 0.24), giving rise to the lineages that generated the remaining sections of *Cypripedium*. Other inter-sectional reticulation events represented more recent hybridizations, producing the lineages leading to sections *Arietinum*, *Bifolia*, *Cypripedium*, and the clade (*Flabellinervia* (*Californica*, *Enantiopedilum*)). The hybrid origins of sections *Bifolia* and *Cypripedium* were also further supported by the most likely network testing for a maximum of one reticulation event (Supplementary Data Fig. S14 C). Section *Subtropica* seemed to have a more complex history, with *C. singchii* Z. J. Liu & L. J. Chen arising between the hybridization of the first diverging lineage in the sister clade of sect. *Retinervia* (γ = 0.86) and another lineage that eventually generated the (*Flabellinervia* (*Californica*, *Enantiopedilum*)) clade (γ = 0.14).

Considering the results of the phylogenetic network analyses testing for reticulation within the subclades containing the three described hybrids, the most likely hybridization networks did not find conclusive evidence for the hybrid status of *C. × alaskanum* (Supplementary Data Fig. S15 and Table S10) and uncovered a more extensive hybridization within the subclades of *C.* × *columbianum* and *C.* × *ventricosum* in sect. *Cypripedium* (Supplementary Data Figs S13 and S16; for further details, see Supplementary Data Results S2).

### Divergence Times

The time-calibrated maximum clade credibility tree, which was produced with BEAST 2 using 20 nuclear genes amounting to 60,141 sites, supported that the subfamily Cypripedioideae diverged from the rest of the slipper orchids close to the K-Pg boundary (66.91 Ma; 95% HPD 88.24–50.26 Ma) while genus *Cypripedium* split from the rest of the slipper orchids in the Oligocene (31.28 Ma; 95% HPD 33.24–29.37 Ma; Supplementary Data Fig. S17). The Mesoamerican sect. *Irapeana* was the first to diverge within the genus, originating in the Early Miocene (22.04 Ma; 95% HPD 27.32–16.48 Ma), followed by the East Asian sect. *Retinervia* (18.5 Ma; 95% HPD 23.27–13.61 Ma) and sect. *Arietinum*, which contains a North American and an East Asian species (16.2 Ma; 95% HPD 20.6–12.15 Ma). After this split, rapid diversification occurred around the Middle Miocene (15.17–12.58 Ma), giving rise to most *Cypripedium* sections or lineages from which the sections diverged (15.17–8.62 Ma). Section *Cypripedium* bifurcated during the Late Miocene (8.32 Ma; 95% HPD 11.12–5.92 Ma), producing its two subclades.

### Diversification Rates

The BAMM analysis illustrated a pattern of an initial elevated net diversification rate during the early stages of *Cypripedium*’s evolution, which steadily declined through time (from 0.2 to 0.11) due to decreasing speciation rate (from 0.25 to 0.16; Fig. 4 A). No significant rate shifts have been detected in the maximum a posteriori probability shift configuration (f = 1; Fig. 4 B), suggesting that a single macroevolutionary rate better explains the diversification within *Cypripedium* over time. Nonetheless, the analysis supported that the internode pairs previously identified to be in the anomaly zone had higher diversification rates compared to the rate at later time points in the phylogeny, providing support to the hypothesis of ILS playing a role in their elevated levels of discordance.

**Figure 4:**
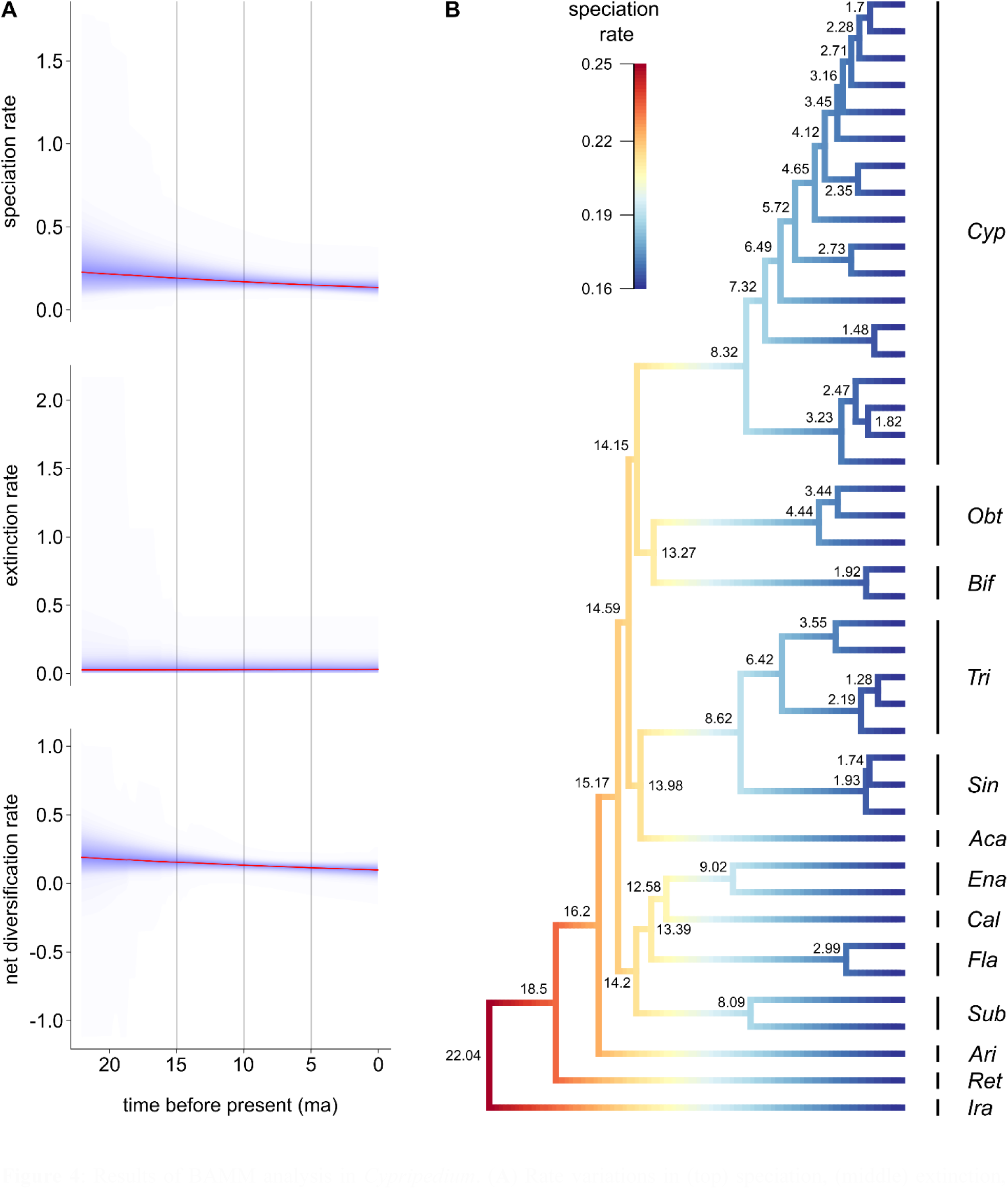
Results of BAMM analysis in *Cypripedium*. (**A**) Rate variations in (top) speciation, (middle) extinction, and (bottom) net diversification through time, based on all samples in the posterior distribution (density shading on confidence regions). (**B**) Maximum a posteriori probability shift configuration represented as a phylorate plot showing variations in speciation rates (cooler colors = slow, warmer colors = fast) along each branch of the dated *Cypripedium* phylogeny (posterior median node height estimates of the divergence times in Ma are shown on the nodes). The clades are annotated with the first three letters of the name of each section.

### Historical Biogeography

Our comparison of biogeographic models in BioGeoBEARS showed that the DEC model (d = 0.0237; e = 0.0078; x = −0.2739; j = 0.; LnL = −129.59) had the lowest AIC (265.185) and AICc (265.647) scores for the first test with nine defined areas, while the DIVALIKE model (d = 0.0229; e = 0.0013; j = 0; LnL = −36.61) had the lowest AIC (77.227) and AICc (77.454) scores for the second test with the New and Old World areas (see Supplementary Data Tables S11 and S12 for further details). The relative probabilities of the ancestral geographical ranges are illustrated with pie charts in Supplementary Data Figs S18 and S19 for the first and second test, respectively, while Fig. 5 combines the single most probable ancestral range from both analyses with the results of the latter shown only for nodes where they disagree.

**Figure 5:**
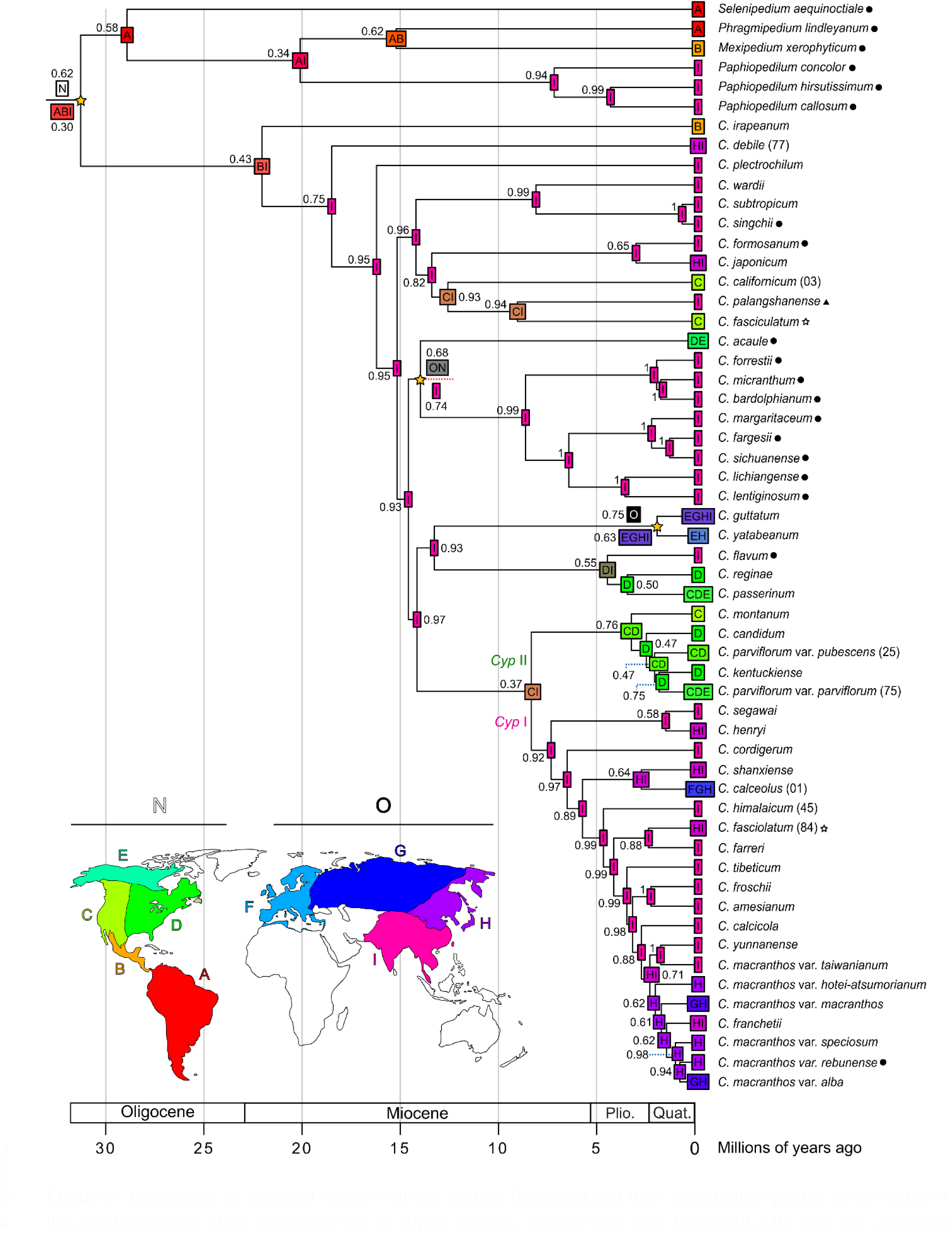
Estimations of ancestral ranges with the highest likelihood and their probabilities plotted at the nodes of the dated maximum clade credibility tree of slipper orchids. Results from both BioGeoBEARS runs (i.e., run with nine areas and with only New and Old World areas) are shown together when they disagree (below and above the branch, respectively; the corresponding nodes are marked with yellow stars); otherwise, only the results of the former test are shown. Distribution states on the nodes are right before cladogenesis. Key to area codes: A = South America; B = Central America and Mexico; C = Western North America; D = Eastern North America; E = Northern North America; F = Western and Central Europe, the Mediterranean, and Scandinavia; G = Eastern Europe and Eurasia; H = Eastern Russia and Northeast Asia; I = Southeast Asia; N = New World; O = Old World. Plio. = Pliocene; Quat. = Quaternary. *Cyp* I = clade I of sect. *Cypripedium*; *Cyp* II = clade II of sect. *Cypripedium* (see Fig. 2).

Regarding the results of the first test, the DEC model indicated that the ancestors of the Cypripedioideae and *Cypripedium* clades were more widespread, distributed across the Old and the New World (ranges ABI and BI, respectively; Fig. 5) and that potential long-distance dispersals and allopatric diversification took place when *Cypripedium* diverged from the rest of the slipper orchids in the Oligocene, as well as when the Mesoamerican sect. *Irapeana* split from the Southeastern ancestor of its sister clade in the Miocene. On the other hand, the DIVALIKE model implemented in the second test supported the New World being the ancestral range of Cypripedioideae, with *Cypripedium* acquiring a wider range across the Old and New World after its split from the rest, and the ancestor of the sister clade of sect. *Irapeana* speciating in the Old World. In both cases, the models suggested that the sister clade of sect. *Irapeana* rapidly diversified in the Old World during the Middle Miocene, specifically in Southeast Asia (i.e., area “I”), where most *Cypripedium* species occur today.

Many of the lineages produced during these rapid diversification events dispersed and speciated in other Old World regions, such as in Northeast Asia and the nearby islands of Japan and Taiwan (e.g., *C. japonicum* Thunb. and *C. formosanum* Hayata; see Supplementary Data Fig. S1 for current distributions). There were also multiple independent dispersals back to the New World between the Miocene and the Pliocene (e.g., sect. *Acaulia,* sect. *Cypripedium*, sect. *Obtusipetala*, and MRCA of sect. *Californica* and *Enantiopedilum*). The MRCA of sect. *Bifolia* spread both Eastwards and Westwards, acquiring a wide distribution in both the Old (i.e., Eastern Europe, Eurasia, Northeast Asia, Japan, East Russia, Southeast Asia) and the New World (i.e., Alaska), with the two species evolving sympatrically in the broader sense.

The MRCAs of clades I and II of sect. *Cypripedium* diverged with the isolation of each clade at the Old and New World during the Late Miocene, respectively. Following this, the two clades diversified allopatrically, with subclade II spreading throughout North America, evolving several closely related taxa within the *C. parviflorum* complex. Within subclade I, some lineages and species expanded their distribution from Southeast Asia to the adjacent area of continental Northeast Asia [e.g., *C. henryi*; MRCA of *C. shanxiense* S. C. Chen and *C. calceolus*; *C. fasciolatum*] as well as the neighboring island of Taiwan [e.g., *C. segawai*, *C. macranthos* var. *taiwanianum* (Masam.) Maekwa]. *C. shanxiense* also spread to the island of Japan, while *C. calceolus* speciated in Northeast Asia, establishing populations in large latitudinal ranges as the only known species in Western Europe, Scandinavia, and the Mediterranean. The ancestor of the *C. macranthos* complex, containing *C. yunnanense* Franch. and *C. franchetii* Wilson, also spread to Northeast Asia, where some of its descendant lineages were subsequently isolated, with certain lineages further dispersing to Japan, producing multiple endemic varieties [e.g., *C. macranthos* var. *rebunense* (Kudo) Ohwi*, C. macranthos* var. *speciosum* (Rolfe) Koidz., and *C. macranthos* var. *hotei-atsumorianum* Sadovsky] while *C. macranthos* var. *macranthos* also expanded to the Far East, Asiatic, and European Russia.

## DISCUSSION

### Monophyly and Topology of Established Infrageneric Taxa

In the present study, we reconstructed the first robust phylogeny of the genus *Cypripedium* based on high-throughput target enrichment and transcriptomic data of 913 nuclear loci using the Orchidaceae963 bait set (Eserman *et al.,* 2021) and sampling all 13 sections. The inferred phylogenetic tree showed that *Cypripedium* is sister to the clade of the other four slipper orchid genera, (*Selenipedium*, (*Paphiopedilum*, (*Phragmipedium*, *Mexipedium*))), which agrees with the topologies recovered by the supra-generic phylogenies of Guo *et al*. (2012), Wong and Peakall (2022), Pérez-Escobar *et al*. (2023), and Liao *et al*. (2024) based on Sanger sequences (chloroplast and nuclear markers), transcriptomic data, a combination of target enrichment (low-copy nuclear loci) and Sanger sequences (matK and ITS), and a combination of plastome and nuclear data, respectively. All sections were monophyletic based on the classification system proposed by Frosch and Cribb (2012) and SC Chen *et al*. (2013). However, subsect. *Macrantha*, which included the clade of *C. farreri* W. W. Sm. and *C. fasciolatum* (as proposed by SC Chen *et al*., 2013), was nested within subsect. *Cypripedium*.

Although the incongruence between the taxonomy and the monophyly of these subsections may be affected by the elevated gene tree discordance within sect. *Cypripedium*, our results match the findings of Szlachetko *et al*. (2021), who also evaluated the monophyly of these subsections based on the same two classification systems. Indeed, most published *Cypripedium* phylogenies supported that one or both subsections may be non-monophyletic and that the (*C. farreri*, *C. fasciolatum*) clade is more closely related to subsect. *Macrantha* rather than subsect. *Cypripedium* (Fatihah *et al*., 2011; J Li *et al*., 2011; Liao *et al*., 2024; H Liu *et al*., 2021*a*), as suggested by SC Chen *et al*. (2013). Moreover, the species composition of the two clades within sect. *Cypripedium* in our phylogeny matched their distributions, with clades I and II only found in the Old World or the New World, respectively. A similar trend was observed in the phylogenies by Fatihah *et al*. (2011), J Li *et al*. (2011), and Szlachetko *et al*. (2021), where the division between the two groups of species seemed to follow their distribution rather than the traditionally used morphological characteristics (i.e., the floral coloration and the shape of the labellum and the lateral petals).

Within these two subclades, we recovered the two morphologically diverse species, *C. parviflorum* and *C. macranthos*, as paraphyletic. Other authors also found that the latter is paraphyletic, with *C. kentuckiense* C. F. Reed embedded in the clade, similar to our phylogeny (Fatihah *et al*., 2011; J Li *et al*., 2011; H Liu *et al*., 2021*a*; Szlachetko *et al*., 2021). However, even though *C. yunnanense* and *C. franchetii* were shown to be closely related to *C. macranthos*, with the latter recovered monophyletic in previous studies (Fatihah *et al*., 2011; J Li *et al*., 2011; H Liu *et al*., 2021*a*; Szlachetko *et al*., 2021), our results showed that they are nested within the *C. macranthos* group, which could have resulted due to the inclusion of different *C. macranthos* varieties [i.e., the ambiguous variety *C. macranthos* var. *alba* (see Supplementary Data Table S2) and all five accepted natural varieties described by Frosch and Cribb, 2012]. *Cypripedium parviflorum* and *C. macranthos* are widespread and morphologically variable species, traditionally distinguished mainly based on flower size and coloration (SC Chen *et al*., 2013; Cribb, 1997). Due to high morphological variation within their infraspecific taxa, as well as the existence of intermediate forms, they have been historically difficult to classify. This difficulty could be attributed to the recent divergence of these varieties and forms within each species that could cause ILS, in addition to hybridization and gene duplication events, which were identified within sect. *Cypripedium* by our analyses.

Besides sect. *Cypripedium*, several supported topologies with high gene tree concordance recovered in our phylogeny, are consistent with those in other studies. For instance, *s*ect. *Irapeana,* the only Neotropical section of *Cypripedium,* was recovered as the sister to the rest in the majority of published molecular phylogenies (Cox *et al*., 1997; Fatihah *et al*., 2011; J Li *et al*., 2011; Guo *et al*., 2012; Liao *et al*., 2024; H Liu *et al*., 2021*a*; Szlachetko *et al*., 2021) as well as in the present study. The placement of sect. *Irapeana* is further supported by morphological and biogeographical data, as it is considered to share “ancestral” features with the plicate-leaved genus *Selenipedium* (e.g., habit, labellum shape, inflorescences with multiple flowers), also found in the Neotropics (Cox *et al.,* 1997; Szlachetko *et al.,* 2021). Similarly to the results of Li *et al*. (2011), Fatihah *et al*. (2011), Szlachetko *et al*. (2021), and Liao *et al*., (2024), sections *Arietinum*, *Retinervia*, and *Irapeana* formed a grade that was sister to the remaining *Cypripedium* included in our phylogeny. Furthermore, sect. *Trigonopedia* and sect. *Sinopedilum*, the two Southeast Asian fly-pollinated sections, previously grouped as a single section (Cribb, 1997), form a clade in our phylogeny. This sister relationship is consistently well-supported by other studies (Fatihah *et al*., 2011; J Li *et al*., 2011; Liao *et al*., 2024; H Liu *et al*., 2021*a*; Szlachetko *et al*., 2021), while the intra-sectional topology we recovered for *Trigonopedia* resembles that of H Liu *et al*. (2021*a*).

Our molecular data also provides support for the classification of sect. *Subtropica* following Frosch and Cribb (2012), as well as sect. *Subtropica* and *Wardiana* following SC Chen *et al*. (2013). Specifically, two species have been described to belong to sect. *Subtropica* in the former monograph, namely, *C. subtropicum* S. C. Chen & K. Y. Lang and *C. wardii* Rolfe, with *C. singchii* being a synonym of *C. subtropicum*. However, SC Chen *et al*. (2013) redefined *C. singchii* as a distinct species and transferred *C. wardii* to its own monotypic section, *Wardiana*. We showed that all three species share an MRCA based on our nuclear and chloroplast molecular data, with *C. subtropicum* and *C. singchii* being more closely related to each other than to *C. wardii*, matching the corresponding topology in J Li *et al*. (2011), Szlachetko *et al*. (2021), and Liao *et al*. (2024).

Concerning the remaining intersectional phylogenetic relationships, we uncovered high discordance at several backbone nodes, both when comparing nuclear gene trees and the nuclear and chloroplast phylogenies. Low backbone branch support has been observed in all molecular phylogenies focusing on the section-level relationships within *Cypripedium* (Cox *et al*., 1997; Fatihah *et al*., 2011; J Li *et al*., 2011; Szlachetko *et al*., 2021), except from H Liu *et al*. (2021*a*) and Liao *et al*. (2024), where four plastid regions, or plastome data and 41 nuclear loci were used for phylogenetic reconstruction, respectively. Our results suggest that gene tree heterogeneity could explain why the evolutionary relationships between the majority of the sections within *Cypripedium* remain unresolved, with different sets of loci and phylogenetic inference methods producing incongruent topologies.

### Rapid Radiation and Hybridization Promoted Diversification

The concordance analyses indicated that most gene tree topologies disagree at certain nodes of our phylogeny. High levels of discordance were particularly observed at nodes along the backbone, the MRCA between sect. *Subtropica* and the (*Flabellinervia* (*Californica*, *Enantiopedilum*)) clade, the inter-sectional relationships between the latter sections, and within the two subclades of sect. *Cypripedium*. Other authors (Fatihah *et al*., 2011; J Li *et al*., 2011; Szlachetko *et al*., 2021) have previously speculated that potential rapid radiation events could interpret the low support at the branches along the backbone of their *Cypripedium* phylogenies, as well as the high morphological differentiation between the sections. Here, we provided supporting evidence for this hypothesis based on our analyses of ∼900 nuclear loci and 80 chloroplast loci. Specifically, our anomaly zone test showed that the backbone incongruence among nuclear gene trees and between the nuclear and chloroplast phylogenies (also observed by Szlachetko et al., 2021), is largely owed to ILS caused by rapid radiation, with sections exhibiting diverging topologies following the split of sect. *Arietinum*. This is further corroborated by the increased diversification rates at the corresponding nodes and their chronologically close placement at the geological time scale.

In addition to the backbone nodes, the majority of the intra-sectional node pairs that were detected to be in the anomaly zone belonged to sect. *Cypripedium*, except a single internode pair within sect. *Sinopedilum*. This result indicates that rapid and recent diversification took place in sect. *Cypripedium*, leading to ILS and producing multiple closely related species and infra-specific taxa with high morphological variation and ambiguous phylogenetic relationships (e.g., within the *C. macranthos* and *C. parviflorum* complexes). However, we showed that other factors, such as reticulation, could have also contributed to the incongruence observed within this section and throughout the phylogeny.

Our phylogenetic network analyses identified multiple potential reticulation events both at the inter- and intra-sectional levels within the phylogeny of *Cypripedium*. Hybridization events likely took place between ancestral and unsampled taxa at inter-sectional nodes deep within the phylogeny, generating the large sister clade of sect. *Irapeana*. More recently diverged taxa may have also hybridized, giving rise to lineages that led to several extant sections, which also revealed evidence of reticulation (for a detailed discussion concerning the results of the intra-sectional PhyloNet analyses, see Supplementary Data Discussion S1). Therefore, hybridization, which is considered to be prevalent within *Cypripedium* in nature (Klier *et al.,* 1991; Hu *et al.,* 2011; Frosch and Cribb, 2012; Szlachetko *et al.,* 2017; Pupulin and Díaz-Morales, 2018), may constitute an important cause of the observed discordance at the backbone nodes between the sections of *Cypripedium*, contributing to the difficulty in the elucidation of their relationships. As for the results of our gene duplication analyses, we identified one relatively large-scale and several smaller-scale gene duplications across different clades of the phylogeny, explaining the elevated numbers of paralogous genes found in different species (∼738 on average per sample). Although Unruh *et al*. (2018) suggested that an increased taxon sampling in *Cypripedium*—the genus with the widest genome size range (4.1–43.1 pg/C) among all slipper orchids—may reveal a potential WGD, we did not identify such an event with our extensive taxon sampling based on the cross-evaluation of our results with a Ks plots analysis. Gene duplications identified through nuclear target enrichment data indicated an increased outlier proportion of duplicated genes at the MRCA of sect. *Cypripedium*, while the transcriptomic data failed to identify a WGD event exclusively shared between Cypripedioideae species. A possible reason for such conflicting results could be that the Orchidaceae963 bait set used for producing the target enrichment data in the present study may target genes that are highly duplicated in the species of sect. *Cypripedium*. These duplications may be confined to certain nuclear loci, but they could also be a result of a large-scale chromosomal duplication, as suggested by the increased number of chromosome counts in sect. *Cypripedium* (2n = 20–36) compared to the usual diploid count in the genus (2n =20), with *C. macranthos* having the widest range of recorded chromosome counts (i.e., 20, 21, 30, 36) in the genus (Eccarius, 2009; SC Chen et al., 2013). Another explanation could be assembly artifacts from target enrichment pipelines that produced a large number of putative gene copies that confounded our WGD analysis, something that needs to be carefully explored in future analyses of WGD based on target enrichment data sets.

### Climatic Fluctuations Influenced Current Distribution and Diversity Patterns

We dated the divergence of the slipper orchids at around the K-Pg boundary (66.91 Ma), similar to Liao *et al*. (2024; ∼64.6 Ma), a more recent time point compared to previous primary molecular calibrations (i.e., ∼75–82 Ma in the Late Cretaceous; Guo *et al*., 2012; Givnish *et al*., 2016). The age of the split of *Cypripedium* from its sister clade was nonetheless similar to previous primary and secondary calibrations (Givnish *et al.,* 2016; Liao *et al*., 2024; H Liu *et al*., 2021*a*), placing the event at ∼30 Ma in the Oligocene. During that time, according to our biogeographic analyses, the ancestors of the slipper orchids either had a wider distribution across the New and the Old World or were initially limited to the New World. Although such uncertainty was also observed by Guo *et al*. (2012) and Liao *et al*. (2024), another study including placeholders from all five orchid subfamilies, all genera of slipper orchids, and 96 outgroup angiosperms by Givnish *et al*. (2016) suggested that the ancestors of Cypripedioideae were in the Neotropics with >75% probability, supporting the result of our second biogeographic analysis that indicated the New World as the ancestral region of slipper orchids.

Our two biogeographic tests agreed on the most likely ancestral regions of the MRCA of *Cypripedium* (i.e., Central America + Southeast Asia or Old World + New World). Again, this partially reflected the results of Guo *et al*. (2012) and matched the results of Givnish *et al*. (2016), supporting the Neotropics + Eurasia as the most probable distribution of the ancestor of *Cypripedium*. However, even though a wider ancestral distribution across the Old and the New World received higher support, it is important to note that due to incomplete taxon sampling as well as the discordance in the backbone of the phylogeny, these results should be interpreted with caution.

A wider ancestral distribution could have been facilitated by the Bering Land Bridge acting as a dispersal corridor between the New and the Old World, as it was intermittently exposed throughout most of the Cenozoic (Hopkins, 1959; see distribution of sect. *Bifolia* at both sides of the Bering Strait in Supplementary Data Fig. S1 L). Combining the results of the biogeographic and the phylogenetic network analyses, it appears that the lineage of *Cypripedium* that dispersed to the Old World had a hybrid origin. The ability of hybrids to adapt to new environments faster than their parent taxa (Kulmuni *et al*., 2024) could be linked to this clade’s inferred adaptation to colder climates after diverging from the Neotropical sister clade (H Liu *et al*., 2021a). This, coupled with a significantly warmer climate (especially at higher latitudes) during the Early to Middle Miocene Climatic Optimum (MMCO; ca. 17–15 Ma; Herold *et al*., 2010; Steinthorsdottir *et al*., 2021), could have allowed *Cypripedium*’s dispersal northwards, via the Bering Land Bridge, or over the strait, and towards Southeast Asia, where the ancestors of extant *Cypripedium* lineages likely further diversified and hybridized.

The Middle Miocene Climate Transition (MMCT; 15–13 Ma), a period of climate transition between the MMCO and the Middle Miocene Glaciation (MMG; Frigola *et al*., 2018), coincided with the rapid radiation of *Cypripedium* in Southeast Asia according to our results. The southeastern margins of the Tibetan Plateau may have provided potential refugia to *Cypripedium* from the cooling climate of the MMCT due to their warmer and more humid conditions owing to the intensification of the Asian monsoons (Xing and Ree, 2017; S Li *et al*., 2020; Zuo *et al*., 2022). Specifically, the isolation of the *Cypripedium* population in deep valleys between mountains, such as the Hengduan mountains, a biodiversity and endemism hotspot harboring the most species of *Cypripedium* (H Liu *et al*., 2021*a*; J Liu *et al*., 2022), could have promoted allopatric diversification at a highly dissected landscape from which several sections or lineages leading to sections diverged, producing the observed pattern of rapid radiations along the backbone of our phylogeny (Favre *et al.,* 2015; Xing and Ree, 2017; Spicer *et al.,* 2020). Additionally, the continuous uplift post-Middle Miocene and the terrain of the mountains at the southeastern margins of the Tibetan Plateau may have provided a diverse variety of habitats throughout its wide altitudinal range, promoting adaptive radiation (López-Pujol *et al.,* 2011; Wang *et al.,* 2012; Xing and Ree, 2017), while the relatively stable conditions of the Hegduan mountains, which glaciated minimally, could have allowed time for speciation to take place (Spicer *et al*., 2020; H Liu *et al*., 2021*a*).

The cooling climate during the Middle to Late Miocene glacial events (Hansen *et al*., 2013; Frigola *et al*., 2018; Methner *et al*., 2020; RM Brown *et al*., 2022) may have prevented the dispersal of many lineages back to the New World through the Bering Land Bridge, besides the most cold tolerant lineages such as sect. *Bifolia* (H Liu *et al*., 2021*a*). Although we did not include models with “jump” speciation at cladogenesis in our final results, “jump” dispersals across the Pacific Ocean may have been a more likely route back to the New World for some lineages [e.g., MRCA of (*Californica, Enantiopedilum*), and New World clades of sections *Obtusipetala* and *Cypripedium*]. Dispersal models have indicated that the small, dust-like orchid seeds can travel a distance of up to 2,000 km by wind (Arditti and Ghani, 2000), but other routes may have also been possible, though uncertain (e.g., epizoochory, water currents, and endozoochory—see Suetsugu *et al.,* 2015; Suetsugu, 2020; Y Zhang *et al.,* 2021).

The glacial cycles of the Late Pliocene and the Quaternary may have promoted the intra-sectional diversification of *Cypripedium*. The isolation of populations in refugia, such as in the mountains at the margins of the Tibetan Plateau, the islands of Japan and Taiwan (Tang *et al.,* 2018), as well as numerous areas in North America (DR Roberts and Hamann, 2015) would have allowed survival and allopatric diversification throughout glacial periods. This could have led to the origin of several endemic taxa (e.g., *C. segawai* in Taiwan and the Taiwanese and Japanese varieties of *C. macranthos*). During the interglacial periods, population expansions resulting in overlapping distributions may have promoted hybridization and introgression, as indicated by the results of our PhyloNet analyses, further increasing genetic and morphological diversity (e.g., in the *C. macranthos* and *C. tibeticum* King ex Rolfe complexes; P Cribb, Royal Botanical Garden, Kew, UK, ‘pers. comm.’).

### Conclusion

The phylogenetic relationships within the genus *Cypripedium* have remained generally unresolved despite numerous phylogenies based on different molecular data and phylogenetic reconstruction methods, largely due to the low support and conflicting topologies at nodes between and within sections. Our study provided a new perspective on the possible causes of this phenomenon, using high-throughput sequence data from over 900 nuclear loci to gain more insight into the evolution of this genus. Although all established sections were supported as monophyletic, we uncovered potential rapid diversification events that led to incomplete lineage sorting at the backbone of the tree. A similar pattern was also observed within the two subclades of the largest section, *Cypripedium*, which did not correspond to its two subsections following Frosch and Cribb (2012) and SC Chen *et al*. (2013) but matched their respective geographic distributions. Additionally, hybridization events were observed at the backbone of *Cypripedium*’s phylogeny and within the sections *Bifolia, Cypripedium,* and *Subtropica*. All these sources of gene tree discordance likely produced conflicting phylogenetic signals observed among nuclear and chloroplast loci resembling the situation seen in other groups, including Amaranthaceae and Malpighiales (Morales-Briones *et al*., 2021; Cai *et al*., 2021), preventing the accurate resolution of *Cypripedium*’s phylogeny.

The events above were associated with climatic transitions that led to major ancestral distribution changes. After the slipper orchids originated in the Neotropics around the K-Pg boundary, the milder climate of the Early Miocene and faster adaptation to colder environments, potentially associated with an ancestral hybridization event, could have allowed them to disperse to the Old World via the Bering Land Bridge. The Middle Miocene Climatic Transition coincided with the rapid radiations following *Cypripedium*’s dispersal to Southeast Asia, possibly due to isolation in local refugia. Several independent dispersals back to the New World via the Bering Land Bridge or across the Pacific Ocean likely took place, while the glacial-interglacial cycles probably played a role in the further speciation and reticulate evolution observed in *Cypripedium*.

## SUPPLEMENTARY DATA

Table S1: Selected examples of taxonomic revisions for the infrageneric classification of *Cypripedium*.

Table S2: *Cypripedium* specimens sampled from the Botanical Collection at Oberhof, associated with the BGM, and the herbarium M.

Table S3: *Cypripedium* DNA samples provided by the DNA collection of the Kew Royal Botanic Gardens.

Table S4: List publicly available orchid sequence data used in this study.

Table S5: Modifications to the Macherey-Nagel NucleoSpin Plant II kit: Genomic DNA from plant (Macherey-Nagel – 07/2014, Rev.09) protocol.

Table S6: *Cypripedium* specimens sampled from the Botanical Collection at Oberhof, associated with the BGM, for the production of transcriptomic data to be subsequently used in the Ks plots analysis.

Table S7: Final occupancy statistics for 913 orthologous nuclear loci and characters (bp) per specimen following the concatenation step.

Table S8: Final occupancy statistics of the 80 orthologous chloroplast loci and characters (bp) per specimen following the concatenation step.

Table S9: The α(x) calculations of the anomaly zone test with the corresponding branch numbers and their branch lengths

Table S10: Calculation and comparison of 10 models produced by the PhyloNet analysis testing for up to one hybridization event in the network including *Cypripedium × alaskanum*.

Table S11: Results table of the BioGeoBEARS analysis using nine areas, showing the AIC and AICc scores and weights of each tested model for comparison.

Table S12: Results table of the BioGeoBEARS analysis using two areas (New and Old World), showing the AIC and AICc scores and weights of each tested model for comparison.

Figure S1: Distribution of *Cypripedium* species per section [classification following Frosch and Cribb (2012); distribution information based on Eccarius (2009), Frosch and Cribb (2012), Chen *et al*. (2013), and Walid et al. (2019)].

Figure S2: The concatenation-based phylogeny of *Cypripedium*, inferred with 913 nuclear loci using IQ-TREE.

Figure S3: The output nuclear phylogeny of *Cypripedium* from the Phyparts analysis.

Figure S4: The output nuclear phylogeny of *Cypripedium* from the Quartet Sampling analysis.

Figure S5: The concatenation-based phylogeny of *Cypripedium*, inferred with 80 chloroplast loci using IQ-TREE.

Figure S6: The output chloroplast *Cypripedium* phylogeny from the Quartet Sampling analysis.

Figure S7: The nuclear ASTRAL phylogeny of *Cypripedium* annotated with the corresponding branch numbers referred to in Supplementary Table S9 for the results of the anomaly zone test calculations and in Figure 3 B.

Figure S8: Results of the polytomy test on the nuclear ASTRAL phylogeny of *Cypripedium*.

Figure S9: Results of the gene duplication test based on the subclade orthogroup tree topology method implemented with the nuclear ASTRAL phylogeny of *Cypripedium*, using the bootstrap filtering approach.

Figure S10: Results of the gene duplication test based on the subclade orthogroup tree topology method implemented with the nuclear ASTRAL phylogeny of *Cypripedium*, using the local topology filtering approach.

Figure S11: Ks plots for the *Cypripedium* transcriptomes generated in this study and additional orchid transcriptomes obtained from the NCBI.

Figure S12: Ks plots comparison between species of the genus *Cypripedium* or between species of the genus *Cypripedium* and other orchids

Figure S13: The total log probabilities of the most likely network from each run for the tests including (A) *C. × columbianum*, (B) *C. × ventricosum,* and (C) taxa from all sections to test for hybridizations at the backbone.

Figure S14: Phylogenetic networks with the highest total log probabilities resulting from the PhyloNet analysis testing for one hybridization event for the extracted subclades with the hybrids (A) *C. × columbianum* and (B) *C. × ventricosum* or (C) subclades from all sections to test for reticulation in the backbone.

Figure S15: Phylogenetic networks including *C. × alaskanum* testing for one hybridization event (using the -po parameter) with the (A) first, (B) second, (C) third, and (D) fourth highest total log probability values.

Figure S16: Phylogenetic networks with the overall highest total log probabilities testing for nine and ten hybridization events for the extracted subclades with the hybrids (A) *C. × columbianum* and (B) *C. × ventricosum*, respectively.

Figure S17: The maximum clade credibility tree of *Cypripedium* resulting from the molecular dating analysis with BEAST 2.

Figure S18: The relative proportions of the ancestral ranges estimated by BioGeoBEARS (nine area test, DEC model) and plotted as pie charts on the corresponding nodes of the dated maximum clade credibility tree of slipper orchids.

Figure S19: The relative proportions of the ancestral ranges estimated by BioGeoBEARS (two-area test, DIVALIKE model) and plotted as pie charts on the corresponding nodes of the dated maximum clade credibility tree of slipper orchids.

Methods S1: Detailed library preparation and target enrichment protocol.

Methods S2: Detailed read processing and assembly protocol for the target enrichment sequence data.

Methods S3: Detailed orthology inference protocol.

Methods S4: Detailed methods regarding the phylogenetic network analyses for the test investigating intra-sectional hybridization within the subclades containing the three described hybrids that were included in this study, their putative parent taxa, and other taxa that share the same MRCA.

Methods S5: Division of nine areas for the biogeographical analyses in BioGeoBEARS.

Methods S6: Input tree and taxon distribution information for the biogeographical analyses in BioGeoBEARS.

Results S1: Species-level results of the *Cypripedium* phylogeny inferred based on target enrichment data of 913 nuclear loci.

Results S2: Detailed phylogenetic network analysis results for the test investigating intra-sectional hybridization within the subclades containing the three described hybrids that were included in this study, their putative parent taxa, and other taxa that share the same MRCA. Discussion S1: Detailed discussion of the phylogenetic network analysis results for the test investigating intra-sectional hybridization within the subclades containing the three described hybrids that were included in this study, their putative parent taxa, and other taxa that share the same MRCA.

## DATA AVAILABILITY

Target enrichment and transcriptomic data generated for this study can be found in the NCBI BioProject XXXXX (please refer to Supplementary Data Tables S2, S3, and S6 for SRA accession numbers). Analyses files are available from the Dryad repository https://doi.org/10.5061/dryad.XXXX

## ACKNOWLEDGEMENTS

We thank the curator of the BSM herbarium (acronym M), Dr. Hans-Joachim Esser, and Mr. Sebastian Urban from the Botanical Collection at Oberhof associated with BGM, for helping with the sampling of the herbarium and living *Cypripedium* specimens, respectively. We would also like to thank Dr. Elizabeth Joyce for providing technical support for the biogeographical analysis, Alina Höwener for her assistance in the preparation of target enrichment libraries, as well as Dr. Phillip Cribb and Tom Velardi, who took the time to share some of their knowledge about *Cypripedium* with us.

Lastly, we would like to acknowledge the people who provided images of the *Cypripedium* taxa included in Figures 1 and 2, namely, Jean-Baptiste Chazalon, Verena Steindl, Luo Chen, Sebastian Urban, Werner Frosch, and the authors of four pictures who made their work available in the Openverse (https://openverse.org/): Figure 1 (a), picture (A): “*Cypripedium irapeanum*” by Moises Béhar is licensed under CC BY-SA 3.0; Figure 1 (a), picture (W): “*Cypripedium reginae*” by Bonnie Isaac is marked with CC0 1.0; Figure 1 (a), picture (X): “Clustered Lady’s-slipper - *Cypripedium fasciculatum*” by Forest Service - Northern Region is marked with Public Domain Mark 1.0.; Figure 1 (b), picture (W): “*Cypripedium kentuckiense* Orchi 2012-05-21 009” by Orchi is licensed under CC BY-SA 3.0.

## LITERATURE CITED

Appendices I, II and III. 2023. https://cites.org/eng/app/appendices.php. 10 Feb. 2023.

Arditti J, Ghani AKA. 2000. Tansley Review No. 110. New Phytologist 145: 367–421.

Atwood JT. 1984. The relationships of the slipper orchids (subfamily Cypripedioideae, Orchidaceae). Selbyana 7: 129–247.

Bezanson J, Karpinski S, Shah V, Edelman A. 2012. Julia: A Fast Dynamic Language for Technical Computing.

Bouckaert R, Vaughan TG, Barido-Sottani J, et al. 2019. BEAST 2.5: An advanced software platform for Bayesian evolutionary analysis. PLOS Computational Biology 15: e1006650.

Brown JW, Walker JF, Smith SA. 2017. Phyx: phylogenetic tools for unix. Bioinformatics 33: 1886–1888.

Brown RM, Chalk TB, Crocker AJ, Wilson PA, Foster GL. 2022. Late Miocene cooling coupled to carbon dioxide with Pleistocene-like climate sensitivity. Nature Geoscience 15: 664–670.

Burkhardt F, Porter DM, Harvey J, Richmond M, Bowler PJ. 1995. The Correspondence of Charles Darwin. Vol. 9: 1861. History and Philosophy of the Life Sciences 17: 173.

Cai L, Xi Z, Lemmon EM, et al. 2021. The Perfect Storm: Gene Tree Estimation Error, Incomplete Lineage Sorting, and Ancient Gene Flow Explain the Most Recalcitrant Ancient Angiosperm Clade, Malpighiales. Systematic Biology 70: 491–507.

Cameron KM, Chase MW, Whitten WM, et al. 1999. A phylogenetic analysis of the Orchidaceae: evidence from rbcL nucleotide. American Journal of Botany 86: 208–224.

Cannon SB, McKain MR, Harkess A, et al. 2015. Multiple Polyploidy Events in the Early Radiation of Nodulating and Nonnodulating Legumes. Molecular Biology and Evolution 32: 193–210.

Chandra N, Singh G, Rai ID, et al. 2023. Predicting Distribution and Range Dynamics of Three Threatened Cypripedium Species under Climate Change Scenario in Western Himalaya. Forests 14: 633.

Chase MW, Cameron KM, Freudenstein JV, et al. 2015. An updated classification of Orchidaceae. Botanical Journal of the Linnean Society 177: 151–174.

Chen H, Zwaenepoel A, Van de Peer Y. 2024. wgd v2: a suite of tools to uncover and date ancient polyploidy and whole-genome duplication. Bioinformatics: btae 272.

Chen SC, Liu ZJ, Chen LJ, Li LQ. 2013. The Genus Cypripedium in China. Peking: Science Press.

Choi B, Crisp MD, Cook LG, et al. 2019. Identifying genetic markers for a range of phylogenetic utility–From species to family level. PLoS ONE 14: e0218995.

Christenhusz MJM, Fay MF, Chase MW. 2017. Plants of the World: An Illustrated Encyclopedia of Vascular Plants. University of Chicago Press.

Cox AV, Pridgeon AM, Albert VA, Chase MW. 1997. Phylogenetics of the slipper orchids (Cypripedioideae, Orchidaceae): Nuclear rDNA ITS sequences. Plant Systematics and Evolution 208: 197–223.

Cribb P. 1997. The Genus Cypripedium. Portland: Timber Press.

Degnan JH, Rosenberg NA. 2006. Discordance of Species Trees with Their Most Likely Gene Trees. PLOS Genetics 2: e68.

Drummond AJ, Suchard MA, Xie D, Rambaut A. 2012. Bayesian Phylogenetics with BEAUti and the BEAST 1.7. Molecular Biology and Evolution 29: 1969–1973.

Eccarius W. 2009. Orchideengattung Cypripedium. EchinoMedia.

Eserman LA, Thomas SK, Coffey EED, Leebens-Mack JH. 2021. Target sequence capture in orchids: Developing a kit to sequence hundreds of single-copy loci. Applications in Plant Sciences 9: e11416.

Fatihah HN, Fay M, Maxted N. 2011. Molecular Phylogenetics of Cypripedium L. (Cypripedioideae: Orchidaceae) Based on Plastid and Nuclear DNA Sequences. Journal of Agrobiotechnology 2: 111–118.

Favre A, Päckert M, Pauls SU, et al. 2015. The role of the uplift of the Qinghai-Tibetan Plateau for the evolution of Tibetan biotas. Biological Reviews 90: 236–253.

Freudenstein JV, van den Berg C, Goldman DH, Kores PJ, Molvray M, Chase MW. 2004. An expanded plastid DNA phylogeny of Orchidaceae and analysis of jackknife branch support strategy. American Journal of Botany 91: 149–157.

Frigola A, Prange M, Schulz M. 2018. Boundary conditions for the Middle Miocene Climate Transition (MMCT v1.0). Geoscientific Model Development 11: 1607–1626.

Frosch W, Cribb P. 2012. Hardy Cypripedium: Species, hybrids and cultivation. Kew Publishing Kew.

Gernhard T. 2008. The conditioned reconstructed process. Journal of Theoretical Biology 253: 769–778.

Givnish TJ, Spalink D, Ames M, et al. 2015. Orchid phylogenomics and multiple drivers of their extraordinary diversification. Proceedings of the Royal Society B: Biological Sciences 282: 20151553.

Givnish TJ, Spalink D, Ames M, et al. 2016. Orchid historical biogeography, diversification, Antarctica and the paradox of orchid dispersal. Journal of Biogeography 43: 1905–1916.

Guo Y-Y, Luo Y-B, Liu Z-J, Wang X-Q. 2012. Evolution and Biogeography of the Slipper Orchids: Eocene Vicariance of the Conduplicate Genera in the Old and New World Tropics. PLOS ONE 7: e38788.

Hansen J, Sato M, Russell G, Kharecha P. 2013. Climate sensitivity, sea level and atmospheric carbon dioxide. Philosophical Transactions of the Royal Society A: Mathematical, Physical and Engineering Sciences 371: 20120294.

Herold N, Müller RD, Seton M. 2010. Comparing early to middle Miocene terrestrial climate simulations with geological data. Geosphere 6: 952–961.

Hopkins DM. 1959. Cenozoic History of the Bering Land Bridge. Science 129: 1519–1528.

Hu S-J, Hu H, Yan N, Huang J-L, Li S-Y. 2011. Hybridization and asymmetric introgression between Cypripedium tibeticum and C. yunnanense in Shangrila County, Yunnan Province, China. Nordic Journal of Botany 29: 625–631.

Huson DH, Scornavacca C. 2012. Dendroscope 3: An Interactive Tool for Rooted Phylogenetic Trees and Networks. Systematic Biology 61: 1061–1067.

Izawa T, Kawahara T, Takahashi H. 2007. Genetic diversity of an endangered plant, Cypripedium macranthos var. rebunense (Orchidaceae): background genetic research for future conservation. Conservation Genetics 8: 1369–1376.

Jiao Y, Leebens-Mack J, Ayyampalayam S, et al. 2012. A genome triplication associated with early diversification of the core eudicots. Genome Biology 13: R3.

Kalyaanamoorthy S, Minh BQ, Wong TKF, von Haeseler A, Jermiin LS. 2017. ModelFinder: fast model selection for accurate phylogenetic estimates. Nature Methods 14: 587–589.

Kim Y-K, Jo S, Cheon S-H, et al. 2020. Plastome Evolution and Phylogeny of Orchidaceae, With 24 New Sequences. Frontiers in Plant Science 11.

Klier K, Leoschke MJ, Wendel JF. 1991. Hybridization and Introgression in White and Yellow Ladyslipper Orchids (Cypripedium candidum and C. pubescens). Journal of Heredity 82: 305–318.

Kolanowska M, Jakubska-Busse A. 2020. Is the lady’s-slipper orchid (Cypripedium calceolus) likely to shortly become extinct in Europe?—Insights based on ecological niche modelling. PLoS ONE 15: e0228420.

Kozlov AM, Darriba D, Flouri T, Morel B, Stamatakis A. 2019. RAxML-NG: a fast, scalable and user-friendly tool for maximum likelihood phylogenetic inference. Bioinformatics 35: 4453–4455.

Kulmuni J, Wiley B, Otto SP. 2024. On the fast track: hybrids adapt more rapidly than parental populations in a novel environment. Evolution Letters 8: 128–136.

Landis MJ, Matzke NJ, Moore BR, Huelsenbeck JP. 2013. Bayesian Analysis of Biogeography when the Number of Areas is Large. Systematic Biology 62: 789–804.

Lanfear R, Frandsen PB, Wright AM, Senfeld T, Calcott B. 2017. PartitionFinder 2: New Methods for Selecting Partitioned Models of Evolution for Molecular and Morphological Phylogenetic Analyses. Molecular Biology and Evolution 34: 772–773.

Li Z, Baniaga AE, Sessa EB, et al. 2015. Early genome duplications in conifers and other seed plants. Science Advances 1: e1501084.

Li Z, De La Torre AR, Sterck L, et al. 2017. Single-Copy Genes as Molecular Markers for Phylogenomic Studies in Seed Plants. Genome Biology and Evolution 9: 1130–1147.

Li S, Ji X, Harrison T, et al. 2020. Uplift of the Hengduan Mountains on the southeastern margin of the Tibetan Plateau in the late Miocene and its paleoenvironmental impact on hominoid diversity. Palaeogeography, Palaeoclimatology, Palaeoecology 553: 109794.

Li J, Liu Z, Salazar GA, et al. 2011. Molecular phylogeny of Cypripedium (Orchidaceae: Cypripedioideae) inferred from multiple nuclear and chloroplast regions. Molecular Phylogenetics and Evolution 61: 308–320.

Liao M, Zhang J-Y, Feng Y, Ren Z-X, Deng H-N, Xu B. 2024. Phylogenomic insights into the historical biogeography, character-state evolution, and species diversification rates of Cypripedioideae (Orchidaceae). Molecular Phylogenetics and Evolution: 108138.

Lindley J. 1840. The genera and species of orchidaceous plants. London: Ridgways.

Linkem CW, Minin VN, Leaché AD. 2016. Detecting the Anomaly Zone in Species Trees and Evidence for a Misleading Signal in Higher-Level Skink Phylogeny (Squamata: Scincidae). Systematic Biology 65: 465–477.

Linnaeus C. 1753. Species plantarum : exhibentes plantas rite cognitas ad genera relatas, cum diferentiis specificis, nominibus trivialibus, synonymis selectis, locis natalibus, secundum systema sexuale digestas. Stockholm: Holmiae, Impensis Laurentii Salvii.

Liu H, Jacquemyn H, Chen W, et al. 2021a. Niche evolution and historical biogeography of lady slipper orchids in North America and Eurasia. Journal of Biogeography 48: 2727–2741.

Liu H, Jacquemyn H, He X, et al. 2021b. The Impact of Human Pressure and Climate Change on the Habitat Availability and Protection of Cypripedium (Orchidaceae) in Northeast China. Plants 10: 84.

Liu J, Milne RI, Zhu G-F, et al. 2022. Name and scale matter: Clarifying the geography of Tibetan Plateau and adjacent mountain regions. Global and Planetary Change 215: 103893.

López-Pujol J, Zhang F-M, Sun H-Q, Ying T-S, Ge S. 2011. Centres of plant endemism in China: places for survival or for speciation? Journal of Biogeography 38: 1267–1280.

Matzke NJ. 2013. BioGeoBEARS: BioGeography with Bayesian (and likelihood) evolutionary analysis in R Scripts. R package, version 0.2 1: 2013.

Matzke NJ. 2014. Model Selection in Historical Biogeography Reveals that Founder-Event Speciation Is a Crucial Process in Island Clades. Systematic Biology 63: 951–970.

Methner K, Campani M, Fiebig J, Löffler N, Kempf O, Mulch A. 2020. Middle Miocene long-term continental temperature change in and out of pace with marine climate records. Scientific Reports 10: 7989.

Minasiewicz J, Znaniecka JM, Górniak M, Kawiński A. 2018. Spatial genetic structure of an endangered orchid Cypripedium calceolus (Orchidaceae) at a regional scale: limited gene flow in a fragmented landscape. Conservation Genetics 19: 1449–1460.

Minh BQ, Schmidt HA, Chernomor O, et al. 2020. IQ-TREE 2: New Models and Efficient Methods for Phylogenetic Inference in the Genomic Era. Molecular Biology and Evolution 37: 1530–1534.

Morales-Briones DF, Kadereit G, Tefarikis DT, et al. 2021. Disentangling Sources of Gene Tree Discordance in Phylogenomic Data Sets: Testing Ancient Hybridizations in Amaranthaceae s.l. Systematic Biology 70: 219–235.

Morales-Briones DF, Gehrke B, Huang C-H, et al. 2022. Analysis of paralogs in target enrichment data pinpoints multiple ancient polyploidy events in Alchemilla sl (Rosaceae). Systematic Biology 71: 190–207.

Nicolè F, Brzosko E, Till-Bottraud I. 2005. Population viability analysis of Cypripedium calceolus in a protected area: longevity, stability and persistence. Journal of Ecology 93: 716–726.

Ortiz EM, Höwener A, Shigita G, et al. 2023. A novel phylogenomics pipeline reveals complex pattern of reticulate evolution in Cucurbitales. bioRxiv: 2023.10.27.564367.

Pease JB, Brown JW, Walker JF, Hinchliff CE, Smith SA. 2018. Quartet Sampling distinguishes lack of support from conflicting support in the green plant tree of life. American journal of botany 105: 385–403.

Pérez-Escobar OA, Dodsworth S, Bogarín D, et al. 2021. Hundreds of nuclear and plastid loci yield novel insights into orchid relationships. American Journal of Botany 108: 1166–1180.

Pérez-Escobar OA, Bogarín D, Przelomska N, et al. 2023. The Origin And Speciation Of Orchids.

Perner H. 2008. Sinopedilum—a new section of the genus Cypripedium. Die Orchidee 59: 35–51.

Pfitzer EHH. 1888. Die Orchideen. In: Engler, A., Prantl, K., (Eds): Die Natiirlichen Pflanzen-familien. Leipzig: Engelmann, 52–96.

Pfitzer EHH. 1894. Beitraege zur Systematik der Orchideen. 19: 1–42.

Pfitzer EHH. 1903. Orchidaceae–Pleonandrae. Leipzig: Engelmann.

POWO. 2023. Plants of the World Online. Facilitated by the Royal Botanic Gardens, Kew. Published on the Internet. http://www.plantsoftheworldonline.org/. 12 Feb. 2023.

Pupulin F, Díaz-Morales M. 2018. On the meaning of Cypripedium × grande (Orchidaceae) and its taxonomic history, with a new name for the nothospecies occurring in Costa Rica and Panama. Phytotaxa 382: 167.

R Core Team. 2023. R: A language and environment for statistical computing.

Rabosky DL, Santini F, Eastman J, et al. 2013. Rates of speciation and morphological evolution are correlated across the largest vertebrate radiation. Nature Communications 4: 1958.

Rabosky DL, Grundler M, Anderson C, et al. 2014. BAMMtools: an R package for the analysis of evolutionary dynamics on phylogenetic trees. Methods in Ecology and Evolution 5: 701–707.

Rafinesque CS. 1836. Flora Telluriana. Philadelphia [Printed for the author by H. Probasco].

Rambaut A, Drummond AJ, Xie D, Baele G, Suchard MA. 2018. Posterior Summarization in Bayesian Phylogenetics Using Tracer 1.7. Systematic Biology 67: 901–904.

Ranwez V, Douzery EJ, Cambon C, Chantret N, Delsuc F. 2018. MACSE v2: toolkit for the alignment of coding sequences accounting for frameshifts and stop codons. Molecular biology and evolution 35: 2582–2584.

Ree RH, Smith SA. 2008. Maximum likelihood inference of geographic range evolution by dispersal, local extinction, and cladogenesis. Systematic Biology 57: 4–14.

Ree RH, Sanmartín I. 2018. Conceptual and statistical problems with the DEC+J model of founder-event speciation and its comparison with DEC via model selection. Journal of Biogeography 45: 741–749.

Reichenbach HG. 1854. Xenia orchidacea 1. Leipzig: Brockhaus.

Roberts DL, Dixon KW. 2008. Orchids: status survey and conservation action plan. Current Biology 18: R325–R329.

Roberts DR, Hamann A. 2015. Glacial refugia and modern genetic diversity of 22 western North American tree species. Proceedings of the Royal Society B: Biological Sciences 282: 20142903.

Rolfe RA. 1896. The Cypripedium group. Orchid Review 4: 327–334.

Ronquist F. 1997. Dispersal-Vicariance Analysis: A New Approach to the Quantification of Historical Biogeography. Systematic Biology 46: 195–203.

Sayers EW, Bolton EE, Brister JR, et al. 2022. Database resources of the national center for biotechnology information. Nucleic Acids Research 50: D20–D26.

Sayyari E, Mirarab S. 2016. Fast coalescent-based computation of local branch support from quartet frequencies. Molecular biology and evolution 33: 1654–1668.

Sayyari E, Mirarab S. 2018. Testing for Polytomies in Phylogenetic Species Trees Using Quartet Frequencies. Genes 9: 132.

Serna-Sánchez MA, Pérez-Escobar OA, Bogarín D, et al. 2021. Plastid phylogenomics resolves ambiguous relationships within the orchid family and provides a solid timeframe for biogeography and macroevolution. Scientific Reports 11: 6858.

Smith SA, Moore MJ, Brown JW, Yang Y. 2015. Analysis of phylogenomic datasets reveals conflict, concordance, and gene duplications with examples from animals and plants. BMC evolutionary biology 15: 1–15.

Smith SA, Brown JW, Walker JF. 2018. So many genes, so little time: A practical approach to divergence-time estimation in the genomic era. PLOS ONE 13: e0197433.

Solís-Lemus C, Bastide P, Ané C. 2017. PhyloNetworks: A Package for Phylogenetic Networks. Molecular Biology and Evolution 34: 3292–3298.

Spicer RA, Farnsworth A, Su T. 2020. Cenozoic topography, monsoons and biodiversity conservation within the Tibetan Region: An evolving story. Plant Diversity 42: 229–254.

Steinthorsdottir M, Coxall HK, de Boer AM, et al. 2021. The Miocene: The Future of the Past. Paleoceanography and Paleoclimatology 36: e2020PA004037.

Suetsugu K, Kawakita A, Kato M. 2015. Avian seed dispersal in a mycoheterotrophic orchid Cyrtosia septentrionalis. Nature Plants 1: 1–2.

Suetsugu K. 2020. A novel seed dispersal mode of Apostasia nipponica could provide some clues to the early evolution of the seed dispersal system in Orchidaceae. Evolution Letters 4: 457–464.

Szlachetko DL, Kolanowska M, Muller F, Vannini J, Rojek J, Górniak M. 2017. First Guatemalan record of natural hybridisation between Neotropical species of the lady’s slipper orchid (Orchidaceae, Cypripedioideae). PeerJ 5: e4162.

Szlachetko DL, Górniak M, Kowalkowska AK, Kolanowska M, Jurczak-Kurek A, Archila Morales F. 2021. The natural history of the genus Cypripedium (Orchidaceae). Plant Biosystems-An International Journal Dealing with all Aspects of Plant Biology 155: 772–796.

Tang CQ, Matsui T, Ohashi H, et al. 2018. Identifying long-term stable refugia for relict plant species in East Asia. Nature Communications 9: 4488.

Unruh SA, McKain MR, Lee Y-I, et al. 2018. Phylotranscriptomic analysis and genome evolution of the Cypripedioideae (Orchidaceae). American Journal of Botany 105: 631–640.

Wang Y, Zheng J, Zhang W, et al. 2012. Cenozoic uplift of the Tibetan Plateau: Evidence from the tectonic–sedimentary evolution of the western Qaidam Basin. Geoscience Frontiers 3: 175–187.

Wen D, Yu Y, Zhu J, Nakhleh L. 2018. Inferring Phylogenetic Networks Using PhyloNet. Systematic Biology 67: 735–740.

Wolfe KH, Li WH, Sharp PM. 1987. Rates of nucleotide substitution vary greatly among plant mitochondrial, chloroplast, and nuclear DNAs. Proceedings of the National Academy of Sciences 84: 9054–9058.

Wong DCJ, Peakall R. 2022. Orchid Phylotranscriptomics: The Prospects of Repurposing Multi-Tissue Transcriptomes for Phylogenetic Analysis and Beyond. Frontiers in Plant Science 13.

Xing Y, Ree RH. 2017. Uplift-driven diversification in the Hengduan Mountains, a temperate biodiversity hotspot. Proceedings of the National Academy of Sciences 114: E3444–E3451.

Yamashita Y, Satoh N, Kurosawa T, Kaneko S. 2023. Genetic diversity and structure of the endangered lady’s slipper orchid Cypripedium japonicum Thunb. (Orchidaceae) in Japan. Population Ecology 65: 54–63.

Yang Y, Smith SA. 2014. Orthology inference in nonmodel organisms using transcriptomes and low-coverage genomes: improving accuracy and matrix occupancy for phylogenomics. Molecular biology and evolution 31: 3081–3092.

Yang Y, Moore MJ, Brockington SF, et al. 2015. Dissecting Molecular Evolution in the Highly Diverse Plant Clade Caryophyllales Using Transcriptome Sequencing. Molecular Biology and Evolution 32: 2001–2014.

Yang Y, Moore MJ, Brockington SF, et al. 2018. Improved transcriptome sampling pinpoints 26 ancient and more recent polyploidy events in Caryophyllales, including two allopolyploidy events. New Phytologist 217: 855–870.

Yu Y, Nakhleh L. 2015. A maximum pseudo-likelihood approach for phylogenetic networks. BMC Genomics 16: S10.

Zhang C, Rabiee M, Sayyari E, Mirarab S. 2018. ASTRAL-III: polynomial time species tree reconstruction from partially resolved gene trees. BMC bioinformatics 19: 15–30.

Zhang C, Mirarab S. 2022. Weighting by Gene Tree Uncertainty Improves Accuracy of Quartet-based Species Trees. Molecular Biology and Evolution 39: msac215.

Zhang G-Q, Liu K-W, Li Z, et al. 2017. The Apostasia genome and the evolution of orchids. Nature 549: 379–383.

Zhang J-Y, Liao M, Cheng Y-H, et al. 2022. Comparative chloroplast genomics of seven endangered Cypripedium species and phylogenetic relationships of Orchidaceae. Frontiers in Plant Science: 2029.

Zhang N, Zeng L, Shan H, Ma H. 2012. Highly conserved low-copy nuclear genes as effective markers for phylogenetic analyses in angiosperms. New Phytologist 195: 923–937.

Zhang Y, Li Y-Y, Wang M, Liu J, Luo F, Lee Y-I. 2021. Seed dispersal in Neuwiedia singapureana: novel evidence for avian endozoochory in the earliest diverging clade in Orchidaceae. Botanical Studies 62: 3.

Zuo M, Sun Y, Zhao Y, Ramstein G, Ding L, Zhou T. 2022. South Asian summer monsoon enhanced by the uplift of Iranian Plateau in Middle Miocene. Climate of the Past Discussions: 1–31.

