## Supplementary Material for "Phylogenomic Analysis of Target Enrichment and Transcriptome Data Uncovers Rapid Radiation and Extensive Hybridization in Slipper Orchid Genus *Cypripedium* L"

Table S7: Final occupancy statistics for 913 orthologous nuclear loci and characters (bp) per specimen following the concatenation step .......... 15-17

Table S8: Final occupancy statistics of the 80 orthologous chloroplast loci and characters (bp) per specimen following the concatenation step .......... 17-19

Figure S11: Ks plots for the *Cypripedium* transcriptomes generated in this study and additional orchid transcriptomes obtained from the NCBI........................................................................ 34-38

Figure S12: Ks plots comparison between species of the genus *Cypripedium* or between species of the genus *Cypripedium* and other orchids ............................................................................ 39-40

Methods S4: Detailed methods regarding the phylogenetic network analyses for the test investigating intra-sectional hybridization within the subclades containing the three described hybrids that were included in this study, their putative parent taxa, and other taxa that share the same MRCA. ...... 51-52

Results S2: Detailed phylogenetic network analysis results for the test investigating intra-sectional hybridization within the subclades containing the three described hybrids that were included in this study, their putative parent taxa, and other taxa that share the same MRCA ................................ 55-56

*Rapid Radiation and Hybridization Promoted Diversification* ............................................................... 56

Discussion S1: Detailed discussion of the phylogenetic network analysis results for the test investigating intra-sectional hybridization within the subclades containing the three described hybrids that were included in this study, their putative parent taxa, and other taxa that share the same MRCA ........................................................................................................................................ 56-59

LITERATURE CITED........................................................................................................... 59

### Supplementary Tables

Table S1: Selected examples of taxonomic revisions for the infrageneric classification of *Cypripedium*.

| (Lindley 1840) | (Pfitzer 1903) | (Cribb 1997) | (Eccarius 2009) | (Frosch and Cribb 2012) | (Chen *et al.* 2013) |
|---|---|---|---|---|---|
| | | **Sect. *Subtropica*** | **Sect. *Subtropica*** | **Sect. *Subtropica*** | **Sect. *Subtropica*** |
| | | *C. subtropicum* | *C. subtropicum* | *C. subtropicum* | *C. subtropicum* |
| | | | | | *C. singchii* |
| | | | | | **Sect. *Wardiana*** |
| | | *C. wardii* | *C. wardii* | *C. wardii* | *C. wardii* |
| ***Foliosa* group** (lateral sepals free at apex) | **Series *Arcuinervia* Sect. *Eucypripedium* Subsect. *Obtusipetala*** | **Sect. *Irapeana*** | **Sect. *Irapeana*** | **Sect. *Irapeana*** | **Sect. *Irapeana*** |
| *C. irapeanum* | *C. irapeanum* | *C. irapeanum* | *C. irapeanum* | *C. irapeanum* | *C. irapeanum* |
| *C. molle* | | *C. molle* | *C. mole* (as *C. irapeanum* ssp. *molle*) | *C. molle* | *C. molle* |
| | | *C. dickinsonianum* | *C. dickinsonianum* | *C. dickinsonianum* | *C. dickinsonianum* |
| | | | | **Sect. *Californica*** | **Sect. *Californica*** |
| | *C. californicum* | *C. californicum* | *C. californicum* | *C. californicum* | *C. californicum* |
| | **Series *Arcuinervia* Sect. *Eucypripedium* Subsect. *Acutipetala*** | **Sect. *Cypripedium* Subsect. *Cypripedium*** | **Sect. *Cypripedium*** | **Sect. *Cypripedium* Subsect. *Cypripedium*** | **Sect. *Cypripedium* Subsect. *Cypripedium*** |
| *C. calceolus* | *C. calceolus* | *C. calceolus* | *C. calceolus* | *C. calceolus* | *C. calceolus* |
| | *C. henryi* | *C. henryi* | *C. henryi* | *C. henryi* | *C. henryi* |
| | | *C. shanxiense* | *C. shanxiense* | *C. shanxiense* | *C. shanxiense* |
| | | *C. segawai* | *C. segawai* | *C. segawai* | *C. segawae* |
| *C. cordigerum* | *C. cordigerum* | *C. cordigerum* | *C. cordigerum* | *C. cordigerum* | *C. cordigerum* |
| *C. parviflorum* (incl. *C. pubescens*) | *C. parviflorum* (incl. *C. pubescens*) | *C. parviflorum* (incl. *C. pubescens*) | *C. parviflorum* (incl. *C. pubescens*) | *C. parviflorum* (incl. *C. pubescens*) | *C. parviflorum* (incl. *C. pubescens*) |
| *C. candidum* | *C. candidum* | *C. candidum* | *C. candidum* | *C. candidum* | *C. candidum* |
| *C. montanum* | *C. montanum* | *C. montanum* | *C. montanum* | *C. montanum* | *C. montanum* |
| | | | *C. kentuckiense* | *C. kentuckiense* | *C. kentuckiense* |
| | *C. fasciolatum* | *C. fasciolatum* | *C. fasciolatum* | *C. fasciolatum* | |
| | | *C. farreri* | *C. farreri* (as *C. fasciolatum* ssp. *farreri*) | *C. farreri* | |

| | | | | | |
|---|---|---|---|---|---|
| | | **Sect. *Cypripedium*** **Subsect. *Macrantha*** | **Sect. *Macrantha*** | **Sect. *Cypripedium*** **Subsect. *Macrantha*** | **Sect. *Cypripedium*** **Subsect. *Macrantha*** |
| | | | | | *C. fasciolatum* |
| | | | | | *C. farreri* |
| | *C. himalaicum* | *C. himalaicum* | *C. himalaicum* | *C. himalaicum* | *C. himalaicum* |
| *C. macranthos* | *C. macranthos* (incl. *C. thunbergii*) | *C. macranthos* | *C. macranthum* | *C. macranthos* | *C. macranthos* |
| | | *C. tibeticum* | *C. tibeticum* | *C. tibeticum* | *C. tibeticum* |
| | *C. corrugatum* | *C. corrugatum* | | | |
| | | *C. calcicola* (as *C. smithii*) | *C. calcicola* (as *C. tibeticum* ssp. *calcicola*) | *C. calcicola* | *C. calcicola* |
| | *C. yunnanense* | *C. yunnanense* | *C. yunnanense* | *C. yunnanense* | *C. yunnanense* |
| | | *C. ludlowii* | *C. ludlowii* (as *C. tibeticum* ssp. *ludlowii*) | *C. ludlowii* | *C. ludlowii* |
| | | *C. franchetii* | *C. franchetii* | *C. franchetii* | *C. franchetii* |
| | | | *C. froschii* (as *C. tibeticum* var. *froschii*) | *C. froschii* | |
| | | | | | *C. taibaiense* |
| | **Series *Arcuinervia*** **Sect. *Enantiopedilum*** | **Sect. *Enantiopedilum*** | **Sect. *Enantiopedilum*** | **Sect. *Enantiopedilum*** | **Sect. *Enantiopedilum*** |
| | *C. fasciculatum* | *C. fasciculatum* | *C. fasciculatum* | *C. fasciculatum* | *C. fasciculatum* |
| | | | | *C. palangshanense* | |
| ***Foliosa* group** (lateral sepals connate to the apex) | **Series *Arcuinervia*** **Sect. *Eucypripedium*** **Subsect. *Obtusipetala*** | **Sect. *Obtusipetala*** | **Sect. *Obtusiflora*** | **Sect. *Obtusipetala*** | **Sect. *Obtusipetala*** |
| | *C. flavum* (as *C. luteum*) | *C. flavum* | *C. flavum* | *C. flavum* | *C. flavum* |
| *C. reginae* (as *C. spectabile*) | *C. reginae* | *C. reginae* | *C. reginae* | *C. reginae* | *C. reginae* |
| *C. passerinum* | *C. passerinum* | *C. passerinum* | *C. passerinum* | *C. passerinum* | *C. passerinum* |
| ***Acaulia* group** | **Series *Arcuinervia*** **Sect. *Eucypripedium*** **Subsect. *Acutipetala*** | **Sect. *Acaulia*** | **Sect. *Acaulia*** | **Sect. *Acaulia*** | **Sect. *Acaulia*** |
| *C. acaule* (as *C. humile*) | *C. acaule* | *C. acaule* | *C. acaule* | *C. acaule* | *C. acaule* |
| | **Sect. *Retinervia*** | **Sect. *Retinervia*** | **Sect. *Retinervia*** | **Sect. *Retinervia*** | **Sect. *Retinervia*** |
| | | *C. elegans* | *C. elegans* | *C. elegans* | *C. elegans* |
| | *C. debile* | *C. debile* | *C. debile* | *C. debile* | *C. debile* |
| | | | | | **Sect. *Palangshanensia*** |

| | | | | | |
|---|---|---|---|---|---|
| | | *C. palangshanense* | *C. palangshanense* | | *C. palangshanense* |
| ***Bifolia* group** | **Series *Arcuinervia*<br>Sect. *Eucypripedium*<br>Subsect. *Obtusipetalum*** | **Sect. *Bifolia*** | **Sect. *Bifolia*** | **Sect. *Bifolia*** | **Sect. *Bifolia*** |
| *C. guttatum* | *C. guttatum* | *C. guttatum* | *C. guttatum* | *C. guttatum* | *C. guttatum* |
| | | *C. yatabeanum* | *C. yatabeanum* (as *C. guttatum* ssp. *yatabeanum*) | *C. yatabeanum* | *C. yatabeanum* |
| | **Series *Flabellinervia*** | **Sect. *Flabellinervia*** | **Sect. *Flabellinervia*** | **Sect. *Flabellinervia*** | **Sect. *Flabellinervia*** |
| *C. japonicum* | *C. japonicum* | *C. japonicum* | *C. japonicum* | *C. japonicum* | *C. japonicum* |
| | | *C. formosanum* | *C. formosanum* | *C. formosanum* | *C. formosanum* |
| ***Arietinum* group** | **Series *Arcuinervia*<br>Sect. *Criosanthes*** | **Sect. *Arietinum*** | **Sect. *Arietinum*** | **Sect. *Arietinum*** | **Sect. *Arietina*** |
| *C. arietinum* | *C. arietinum* | *C. arietinum* | *C. arietinum* | *C. arietinum* | *C. arietinum* |
| | | *C. plectrochilum* | *C. plectrochilum* | *C. plectrochilum* | *C. plectrochilum* |
| | ***Series Arcuinervia*<br>Sect. *Trigonopedilum*** | **Sect. *Trigonopedia*** | **Sect. *Trigonopedia*** | **Sect. *Trigonopedia*** | **Sect. *Trigonopedium*** |
| | *C. margaritaceum* | *C. margaritaceum* | *C. margaritaceum* | *C. margaritaceum* | *C. margaritaceum* |
| | | *C. lichiangense* | *C. lichiangense* | *C. lichiangense* | *C. lichiangense* |
| | | *C. wumengense* | | *C. wumengense* | *C. wumengense* |
| | *C. fargesii* (as *C. ebracteatum*) | *C. fargesii* | *C. fargesii* (as *C. margaritaceum* ssp. *fargesii*) | *C. fargesii* | *C. fargesii* |
| | | | *C. lentiginosum* (as *C. lichiangense* ssp. *lentiginosum*) | *C. lentiginosum* | *C. lentiginosum* |
| | | | *C. sichuanense* (as *C. margaritaceum* ssp. *sichuanense*) | *C. sichuanense* | *C. sichuanense* |
| | | | | | *C. daweishanense* |
| | | | | | *C. malipoense* |
| | **Series *Arcuinervia*<br>Sect. *Enantiopedilum*** | | **Sect. *Sinopedilum*** | **Sect. *Sinopedilum*** | **Sect. *Sinopedilum*** |
| | *C. micranthum* | *C. micranthum* | *C. micranthum* | *C. micranthum* | *C. micranthum* |
| | | *C. bardolphianum* | *C. bardolphianum* | *C. bardolphianum* | *C. bardolphianum* |
| | | *C. forrestii* | *C. forrestii* (as *C. bardolphianum* ssp. *forrestii*) | *C. forrestii* | *C. forrestii* |

Table S2: *Cypripedium* specimens sampled from the Botanical Collection at Oberhof, associated with the BGM, and the herbarium M.

| Taxon | Accepted Name [a] | Lab No. [b] | Collection/ Herbarium | Collection Number | Collection Location | SRA Accession No. |
|---|---|---|---|---|---|---|
| *Cypripedium acaule* Aiton | - | 74* | Oberhof | - | - | - |
| *Cypripedium amesianum* Schltr. | *C. yunnanense* Franch. | 35* | Oberhof | - | - | - |
| *Cypripedium bardolphianum* W. W. Sm. & Farrer | - | 36 | Oberhof | 2023/1214-1 | - | - |
| *Cypripedium calceolus* L. | - | 1* | Oberhof | 2023/1209-1 | - | - |
| *Cypripedium calceolus* L. | - | 44* | Oberhof | 2023/1256-1 | - | - |
| *Cypripedium calcicola* Schltr. | - | 2* | Oberhof | 2023/1193-1 | - | - |
| *Cypripedium californicum* A. Gray | - | 3* | Oberhof | - | - | - |
| *Cypripedium candidum* Muehl. ex Willd. | - | 4* | Oberhof | - | - | - |
| *Cypripedium cordigerum* D. Don | - | 5* | Oberhof | 2023/1132-1 | - | - |
| *Cypripedium debile* Rchb.f. | - | 77* | Oberhof | 2023/1228-1 | - | - |
| *Cypripedium fargesii* Franch. | - | 47* | Oberhof | 2023/1218-1 | - | - |
| *Cypripedium farreri* W. W. Sm. | - | 6* | Oberhof | 2023/1213-1 | - | - |
| *Cypripedium fasciolatum* Franch. | - | 7* | Oberhof | 2023/1155-1 | - | - |
| *Cypripedium flavum* P. F. Hunt & Summerh. | - | 8* | Oberhof | 2023/1172-1 | - | - |
| *Cypripedium formosanum* Hayata | - | 9* | Oberhof | 2023/1202-1 | - | - |
| *Cypripedium franchetii* Wilson | - | 10* | Oberhof | 2023/1150-1 | - | - |
| *Cypripedium froschii* Perner | - | 11* | Oberhof | 2023/1126-1 | - | - |
| *Cypripedium guttatum* Swartz | - | 12* | Oberhof | 2023/1261-1 | - | - |
| *Cypripedium henryi* Rolfe | - | 13 | Oberhof | 2023/1179-1 | - | - |
| *Cypripedium himalaicum* Rolfe | - | 45* | Oberhof | - | - | - |
| *Cypripedium irapeanum* La Llave & Lex. | - | 14* | Oberhof | - | - | - |
| *Cypripedium japonicum* Thunb. | - | 15* | Oberhof | 2023/1147-1 | - | - |
| *Cypripedium kentuckiense* C. F. Reed | - | 16* | Oberhof | 2023/1166-1 | - | - |
| *Cypripedium lentiginosum* P. J. Cribb & S. C. Chen | - | 37 | Oberhof | - | - | - |
| *Cypripedium lichiangense* S. C. Chen & P. J. Cribb | - | 39 | Oberhof | 2023/ 1223-1 | - | - |
| *Cypripedium macranthos* Sw. var. *macranthos* | - | 17* | Oberhof | 2023/1242-1 | - | - |
| *Cypripedium macranthos* var. *alba* | Probably *C. macranthos* var. *albiflorum* Makino (now synonym of *C. macranthos* Sw. var. *macranthos*), or *C. macranthos* var. *album* Mandl | 18* | Oberhof |  | - | - |

| | | | | | | |
|---|---|---|---|---|---|---|
| *Cypripedium macranthos* ‘var. *hotei-atsmorianum* Sadovsky*’* | - | 19* | Oberhof | 2023/1176-1 | - | - |
| *Cypripedium macranthos* ‘var. *rebunense* (Kudo) Ohwi*’* | - | 20* | Oberhof | 2023/1151-1 | - | - |
| *Cypripedium macranthos* var. *speciosum* (Rolfe) Koidz. | - | 21* | Oberhof | 2023/1186-1 | - | - |
| *Cypripedium macranthos* ‘var. *taiwanianum* (Masam.) Maekwa’ | - | 46* | Oberhof | 2023/1219-1 | - | - |
| *Cypripedium micranthum* Franch. | - | 22* | Oberhof | 2023/1194-1 | - | - |
| *Cypripedium montanum* Douglas ex Lindl. | - | 23* | Oberhof | 2023/1212-1 | - | - |
| *Cypripedium parviflorum* Salisb var. *parviflorum* | - | 75* | Oberhof | 2023/1216-1 | - | - |
| *Cypripedium parviflorum* Salisb var. *makasin* (Farw.) C. J. Sheviak | *C. parviflorum* Salisb var. *parviflorum* | 24* | Oberhof | 2023/1192-1 | - | - |
| *Cypripedium parviflorum* Salisb. *var. pubescens* (Willd.) | - | 25* | Oberhof | 2023/1144-1 | - | - |
| *Cypripedium parviflorum* Salisb. var. *pubescens (*Willd.) O. W. Knigh*t forma planipetalum* (Fernald) P. J. Cribb | - | 26* | Oberhof | 2023/1210-1 | - | - |
| *Cypripedium passerinum* Richardson | - | 42* | Oberhof | 2023/1269-1 | - | - |
| *Cypripedium plectrochilum* Franch. | - | 27* | Oberhof | 2023/1217-1 | - | - |
| *Cypripedium reginae* Walter | - | 40* | Oberhof | 2023/1196-1 | - | - |
| *Cypripedium reginae* var. *alba* | *C. reginae* var. *album* (Aiton) Rolfe | 41* | Oberhof | 2023/1162-1 | - | - |
| *Cypripedium segawai* Masam. | - | 28* | Oberhof | 2023/1170-1 | - | - |
| *Cypripedium shanxiense* S. C. Chen | - | 29* | Oberhof | - | - | - |
| *Cypripedium subtropicum* S. C. Chen & K. Y. Lang | - | 76* | Oberhof | 2023/1233-1 | - | - |
| *Cypripedium tibeticum* King ex Rolfe | - | 30* | Oberhof | 2023/1127-1 | - | - |
| *Cypripedium wardii* Rolfe | - | 38* | Oberhof | - | - | - |
| *Cypripedium yatabeanum* Makino | - | 31* | Oberhof | 2023/1260-1 | - | - |
| *Cypripedium yunnanense* Franch. | - | 32* | Oberhof | - | - | - |
| *Cypripedium* × *alaskanum* P. M. Br. | - | 43* | Oberhof | 2023/1265-1 | - | - |
| *Cypripedium* × *columbianum* Sheviak | - | 33* | Oberhof | 2023/1130-1 | - | - |
| *Cypripedium* × *ventricosum* Sw. | - | 34* | Oberhof | 2023/1133-1 | - | - |
| *Cypripedium acaule* Aiton | - | 48* | M | Eric A. Bourdo, Jr. 28,450 (1974) | Michigan, U.S.A. | - |
| *Cypripedium californicum* A. Gray | - | 57* | M | Mary F. Spencer s.n. (1916) | California, U.S.A. | - |

| | | | | | | |
|---|---|---|---|---|---|---|
| *Cypripedium calceolus* L. var. *parviflorum* (Salsib) Fernald | *C. parviflorum* Salisb var. *parviflorum* | 53* | M | W. J. Dress 5965 (1958) | New York, U.S.A. | - |
| *Cypripedium calceolus* var. *pubescens* (Willd.) Correll | *Cypripedium parviflorum* Salisb. *var. pubescens* (Willd.) | 69* | M | Eric A. Bourdo 32295 (1976) | Michigan, U.S.A. | - |
| *Cypripedium debile* Rchb.f. | - | 62 | M | B. Dickoré 14259 (1996) | NW Yunnan, China | - |
| *Cypripedium himalaicum* Rolfe | - | 65* | M | F. Lobbichler 121 (1955) | Manangbhot, Nepal | - |
| *Cypripedium passerinum* Richardson var. *passerinum* | - | 52 | M | W. J. Cody 4044 (1950) | Mackenzie District, Northwest Territories, Canada | - |

[a] According to Frosch and Cribb (2012).

[b] Lab numbers followed by an asterisk ("*") indicate samples included in the cpDNA tree.

Table S3: *Cypripedium* DNA samples provided by the DNA collection of the Kew Royal Botanic Gardens.

| **Taxon** | **Lab No.** [a] | **Collector(s)** | **Collector Number** | **Collection Date** | **Country** | **Voucher** | **MWC** | **Dnald** | **SRA Accession No.** |
|---|---|---|---|---|---|---|---|---|---|
| *Cypripedium bardolphianum* W. W. Sm. & Farrer | 83* | Huang Long Si | s. n. | 1997-06 | China | Huang Long Si | 5722 | 5722 | - |
| *Cypripedium fasciculatum* Kellogg ex S. Watson | 88* | H. Peruv | (4/6/1996) | - | USA, Washington State, Killitas County, Mineral Springs. | H. Peruv (4/6/1996) | O-1269 | 5269 | - |
| *Cypripedium fasciolatum* Franch. | 84* | Huang Long Si | s. n. | 1997-06 | China | Huang Long Si | 5723 | 5723 | - |
| *Cypripedium lichiangense S. C. Chen & P. J. Cribb* | 87* | P. Cribb | (6/8/92) | - | Unknown | P. Cribb (6/8/92) | O-953 | 4953 | - |

[a] Lab numbers followed by an asterisk ("*") indicate samples included in the cpDNA tree.

Table S4: List publicly available orchid sequence data used in this study.

| Taxon | Accepted Name [a] | Source/Bioproject Accession Number | NCBI Accession Number [b] | Sequence type [c] |
|---|---|---|---|---|
| *Apostasia shenzhenica* Z. J. Liu & L. J. Chen | - | (Zhang *et al.* 2017) | - | genome* |
| *Apostasia shenzhenica* Z. J. Liu & L. J. Chen | - | PRJNA927338 | NC039812 | chloroplast genome |
| *Cypripedium acaule* Aiton | - | PRJNA412930 | SRX3240244 | transcriptome*† |
| *Cypripedium bardolphianum* W. W. Sm. & Farrer | - | PRJNA479379 | SRX4336453 | transcriptome*† |
| *Cypripedium bardolphianum* W. W. Sm. & Farrer | - | Hu *et al.,* 2022 | OL741711 | chloroplast genome |
| *Cypripedium calceolus* L. | - | PRJNA927338 | NC045400 | chloroplast genome |
| *Cypripedium debile* Rchb.f. | - | PRJNA838021 | SRX15379764 | genome skimming |
| *Cypripedium debile* Rchb.f. | - | PRJNA927338 | NC063681 | chloroplast genome |
| *Cypripedium fargesii* Franch. | - | PRJNA479379 | SRX4336442 | transcriptome*† |
| *Cypripedium fargesii* Franch. | - | PRJNA927338 | NC084418 | chloroplast genome |
| *Cypripedium farreri* W. W. Sm. | - | Hu *et al.,* 2022 | OM066273 | chloroplast genome |
| *Cypripedium fasciolatum* Franch. | - | Hu *et al.,* 2022 | OM066274 | chloroplast genome |
| *Cypripedium flavum* P. F. Hunt & Summerh. | - | PRJNA479379 | SRX4336448 | transcriptome*† |
| *Cypripedium flavum* P. F. Hunt & Summerh. | - | Hu *et al.,* 2022 | OM066275 | chloroplast genome |
| *Cypripedium formosanum* Hayata | - | PRJNA277578 | SRX911751 | transcriptome*† |
| *Cypripedium formosanum* Hayata | - | PRJNA927338 | NC026772 | chloroplast genome |
| *Cypripedium forrestii* P. J. Cribb | - | PRJNA1029356 | SRX22160934 | transcriptome† |
| *Cypripedium guttatum* Swartz | - | Hu *et al.,* 2022 | OM066278 | chloroplast genome |
| *Cypripedium henryi* Rolfe | - | Hu *et al.,* 2022 | OM066279 | chloroplast genome |
| *Cypripedium japonicum* Thunb. | - | Hu *et al.,* 2022 | OM066280 | chloroplast genome |
| *Cypripedium macranthos* 'var. *rebunense* (Kudo) Ohwi' | - | PRJDB15443 | DRX436158 | transcriptome |
| *Cypripedium margaritaceum* Franch. | - | PRJNA479379 | SRX4336454 | transcriptome*† |
| *Cypripedium micranthum* Franch. | - | PRJNA479379 | SRX4336451 | transcriptome*† |
| *Cypripedium lentiginosum* P. J. Cribb & S. C. Chen | - | PRJNA479379 | SRX4336441 | transcriptome*† |
| *Cypripedium lichiangense* S. C. Chen & P. J. Cribb | - | PRJNA1029356 | SRX22160950 | transcriptome† |
| *Cypripedium lichiangense* S. C. Chen & P. J. Cribb | - | PRJNA927338 | NC084419 | chloroplast genome |
| *Cypripedium palangshanense* T. Tang & F. T. Wang | - | PRJNA838021 | SRX15379763 | genome skimming |
| *Cypripedium palangshanense* T. Tang & F. T. Wang | - | PRJNA927338 | NC063680 | chloroplast genome |
| *Cypripedium plectrochilum Franch.* | - | Hu *et al.,* 2022 | OM066284 | chloroplast genome |
| *Cypripedium sichuanense* Perner | - | PRJNA479379 | SRX4336445 | transcriptome*† |
| *Cypripedium sichuanense* Perner | - | PRJNA927338 | NC084420 | chloroplast genome |

| | | | | |
|---|---|---|---|---|
| *Cypripedium subtropicum* S. C. Chen & K. Y. Lang | - | PRJNA927338 | NC053551 | chloroplast genome |
| *Cypripedium singchii* Z. J. Liu & L. J. Chen | *Cypripedium subtropicum* S. C. Chen & K. Y. Lang | PRJNA479379 | SRX4336446 | transcriptome*† |
| *Cypripedium* × *ventricosum Sw.* | - | Hu *et al.,* 2022 | OM066286 | chloroplast genome |
| *Dendrobium catenatum* Lindl. | *Dendrobium officinale* Kimura & Migo | (Zhang *et al.* 2016) | - | genome* |
| *Dendrobium catenatum* Lindl. | *Dendrobium officinale* Kimura & Migo | PRJNA927338 | NC037361 | chloroplast genome |
| *Mexipedium xerophyticum* (Soto Arenas, Salazar & Hágsater) V. A. Albert & M. W. Chase | - | PRJNA412930 | SRX3240239 | transcriptome* |
| *Mexipedium xerophyticum* (Soto Arenas, Salazar & Hágsater) V. A. Albert & M. W. Chase | | PRJNA927338 | NC069868 | chloroplast genome |
| *Paphiopedilum callosum* Pfitzer | - | PRJNA412930 | SRX3240238 | transcriptome* |
| *Paphiopedilum callosum* Pfitzer | - | PRJNA927338 | NC069960 | chloroplast genome |
| *Paphiopedilum concolor* Pfitzer | - | PRJNA252662 | SRX601820 | transcriptome* |
| *Paphiopedilum concolor* Pfitzer | - | PRJNA927338 | NC069964 | chloroplast genome |
| *Paphiopedilum hirsutissimum* Pfitzer | - | PRJNA252662 | SRX601821 | transcriptome* |
| *Paphiopedilum hirsutissimum* Pfitzer | - | PRJNA927338 | NC050871 | chloroplast genome |
| *Phalaenopsis equestris* (Schauer) Rchb.f. | - | (Cai *et al.* 2015) | - | genome* |
| *Phalaenopsis equestris* (Schauer) Rchb.f. | - | PRJNA927338 | NC017609 | chloroplast genome |
| *Phragmipedium lindleyanum* (R. H. Schomb. ex Lindl.) Rolfe | - | PRJNA412930 | SRX3240237 | transcriptome*† |
| *Selenipedium aequinoctiale* Garay | - | PRJNA412930 | SRX3240236 | transcriptome*† |
| *Vanilla planifolia* Andrews | - | (Piet *et al.* 2022) | - | genome* |
| *Vanilla planifolia* Andrews | - | PRJNA927338 | NC026778 | chloroplast genome |
| *Vanilla shenzhenica* Z. J. Liu & S. C. Chen | - | PRJNA310678 | SRX2938656 | transcriptome*† |

[a]According to Frosch and Cribb (2012) for *Cypripedium*, or POWO (2023) for the rest.

[b]Accession numbers starting with "SR" and "DR" were acquired from the SRA database of NCBI and accession numbers starting with "NC", "OM" and "OL" were acquired from NCBI GenBank.

[c] Transcriptomes and genomes with an asterisk ("*") were used to create the set of references to improve gene extraction from the target enrichment sequence data. Transcriptomes with a cross ("†") were included in the cpDNA tree.

Table S5: Modifications to the Macherey-Nagel NucleoSpin Plant II kit: Genomic DNA from plant (Macherey-Nagel – 07/2014, Rev.09) protocol.

| Step Nr. | Modified instructions |
|---|---|
| **1** | Approximately 25 mg (or less, in cases where <25 mg was available) of silica-dried or herbarium leaf material was fragmented with forceps and placed into a 2 mL microcentrifuge tube with two glass beads (6 mm in diameter). Next, the samples were homogenized using a Retsch Tissuelyser for 5 –10 mins at 30/s, until the tissue turned into very fine powder. |
| **2a** | The tubes containing the homogenized tissue were centrifuged shortly. 600 µL Buffer PL1 and 10 µL RNase A (from the MN kit) were added, and the samples were vortexed thoroughly. The suspensions were incubated on a thermomixer for 60 mins at 65 °C and 400 rpm, and they were shortly vortexed every 15 mins. |
| **6** | **1$^{st}$ wash**: 400 µL Buffer PW1 were added to the columns, and they were centrifuged for 1 min at 11,000 x g. The flow-through was discarded.<br>**2$^{nd}$ wash**: 600 µL Buffer PW2 were added to the columns, and they were centrifuged for 1 min at 11,000 x g. The flow-through was discarded.<br>**3$^{rd}$ wash**: 350 µL Buffer PW2 were added to the columns, and they were centrifuged for 1 min at 11,000 x g. The flow-through was discarded.<br>**4$^{th}$ wash**: 200 µL Buffer PW2 were added to the columns, and they were centrifuged for 2 min at 11,000 x g. The flow-through was discarded. |
| **7** | **Eluate A:** The column was placed in a new 1.5 mL microcentrifuge tube, and 50 µL Buffer PE (65 °C) was pipetted onto the membrane. The samples were incubated on a thermomixer for 5 mins at 65 °C and 300 rpm. Then, they were centrifuged for 1 min at 11,000 x g to elute the DNA.<br>**Eluate B:** The previous step was repeated by placing the column into another 1.5 mL microcentrifuge tube to produce a second eluate with another 50 µL Buffer PE (65 °C), to avoid decreasing the concentration of the first eluate.<br>The eluates were stored at -20 °C until further use. |

Table S6: *Cypripedium* specimens sampled from the Botanical Collection at Oberhof, associated with the BGM, for the production of transcriptomic data to be subsequently used in the Ks plots analysis.

| **Taxon** | **Lab No.** | **Collection Number** | **SRA Accession No.** |
|---|---|---|---|
| *Cypripedium californicum* A. Gray | 95 | 2023/1173-1 | - |
| *Cypripedium guttatum* Swartz | 12 | 2023/1261-1 | - |
| *Cypripedium henryi* Rolfe | 97 | 2023/1158-1 | - |
| *Cypripedium irapeanum* La Llave & Lex. | 94 | - | - |
| *Cypripedium kentuckiense* C. F. Reed | 98 | 2023/1237-1 | - |
| *Cypripedium parviflorum* Salisb var. *parviflorum* | 90 | 2023/1249-1 | - |
| *Cypripedium parviflorum* Salisb. *var. pubescens* (Willd.) | 91 | 2023/1138-1 | - |
| *Cypripedium plectrochilum* Franch. | 92 | 2023/1278-1 | - |
| *Cypripedium segawai* Masam. | 96 | 2023/1230-1 | - |
| *Cypripedium yatabeanum* Makino | 31 | 2023/1260-1 | - |

Table S7: Final occupancy statistics for 913 orthologous nuclear loci and characters (bp) per specimen following the concatenation step.

| **Taxon ID** [a] | **#of orthologs** | **# of characters** | **% of orthologs** | **% of characters** |
|---|---|---|---|---|
| C_palangshanense_gks | 18 | 12030 | 0.02 | 0.01 |
| C_debile_62 | 53 | 35231 | 0.06 | 0.04 |
| C_debile_gks | 60 | 20319 | 0.07 | 0.02 |
| C_passerinum_var_passerinum_52 | 181 | 128217 | 0.20 | 0.13 |
| C_himalaicum_65 | 236 | 184170 | 0.26 | 0.19 |
| C_calceolus_var_par_53 | 283 | 209943 | 0.31 | 0.21 |
| C_calceolus_var_pub_69 | 310 | 230140 | 0.34 | 0.24 |
| C_calceolus_44 | 441 | 433171 | 0.48 | 0.44 |
| C_acaule_48 | 442 | 354653 | 0.48 | 0.36 |
| C_lichiangense_39 | 449 | 391080 | 0.49 | 0.40 |
| C_henryi_13 | 455 | 405944 | 0.50 | 0.41 |
| C_lentiginosum_37 | 470 | 422874 | 0.51 | 0.43 |
| C_franchetii_10 | 493 | 457599 | 0.54 | 0.47 |
| C_froschii_11 | 508 | 469179 | 0.56 | 0.48 |
| C_shanxiense_29 | 535 | 543326 | 0.59 | 0.56 |
| C_californicum_57 | 545 | 432531 | 0.60 | 0.44 |
| C_bardolphianum_36 | 545 | 508112 | 0.60 | 0.52 |
| C_formosanum_09 | 561 | 532424 | 0.61 | 0.54 |
| C_guttatum_12 | 572 | 514172 | 0.63 | 0.53 |
| C_segawai_28 | 609 | 585758 | 0.67 | 0.60 |
| C_lichiangense_87 | 610 | 563905 | 0.67 | 0.58 |
| C_montanum_23 | 620 | 654321 | 0.68 | 0.67 |
| C_amesianum_35 | 626 | 596568 | 0.69 | 0.61 |
| C_acaule_74 | 627 | 568915 | 0.69 | 0.58 |
| C_par_var_pub_f_planipetalum_26 | 634 | 646728 | 0.69 | 0.66 |
| C_x_columbianum_33 | 643 | 643758 | 0.70 | 0.66 |
| C_cordigerum_05 | 646 | 670949 | 0.71 | 0.69 |
| C_kentuckiense_16 | 646 | 654250 | 0.71 | 0.67 |
| C_par_var_makasin_24 | 646 | 659109 | 0.71 | 0.67 |
| C_par_var_parviflorum_75 | 648 | 660305 | 0.71 | 0.67 |
| C_par_var_pub_25 | 649 | 654967 | 0.71 | 0.67 |
| C_passerinum_42 | 651 | 632800 | 0.71 | 0.65 |
| C_candidum_04 | 661 | 676019 | 0.72 | 0.69 |
| C_mac_var_hotei-atsumorianum_19 | 670 | 683432 | 0.73 | 0.70 |
| C_tibeticum_30 | 671 | 659175 | 0.73 | 0.67 |
| C_farreri_06 | 671 | 670480 | 0.73 | 0.69 |
| C_x_ventricosum_34 | 671 | 651993 | 0.73 | 0.67 |
| C_yunnanense_32 | 674 | 671975 | 0.74 | 0.69 |
| C_himalaicum_45 | 676 | 683549 | 0.74 | 0.70 |

| | | | | |
|---|---|---|---|---|
| C_fargesii_47 | 678 | 684155 | 0.74 | 0.70 |
| C_micranthum_22 | 679 | 675347 | 0.74 | 0.69 |
| C_fasciolatum_07 | 680 | 669205 | 0.74 | 0.68 |
| C_fasciolatum_84 | 681 | 688907 | 0.75 | 0.70 |
| C_calceolus_01 | 682 | 657206 | 0.75 | 0.67 |
| C_mac_var_speciosum_21 | 684 | 684491 | 0.75 | 0.70 |
| C_mac_var_alba_18 | 685 | 691253 | 0.75 | 0.71 |
| C_mac_var_taiwanianum_46 | 687 | 686315 | 0.75 | 0.70 |
| C_reginae_var_alba_41 | 689 | 676238 | 0.75 | 0.69 |
| C_mac_var_rebunense_20 | 693 | 703280 | 0.76 | 0.72 |
| C_subtropicum_76 | 693 | 688927 | 0.76 | 0.70 |
| C_wardii_38 | 695 | 684415 | 0.76 | 0.70 |
| C_calcicola_02 | 697 | 701028 | 0.76 | 0.72 |
| C_mac_var_mac_17 | 698 | 692451 | 0.76 | 0.71 |
| C_bardolphianum_83 | 700 | 696571 | 0.77 | 0.71 |
| C_reginae_40 | 705 | 693617 | 0.77 | 0.71 |
| C_debile_77 | 707 | 670775 | 0.77 | 0.69 |
| C_flavum_08 | 708 | 697712 | 0.78 | 0.71 |
| C_fasciculatum_88 | 709 | 701729 | 0.78 | 0.72 |
| C_yattabeanum_31 | 712 | 708863 | 0.78 | 0.72 |
| C_mac_var_rebunense_trp | 718 | 786843 | 0.79 | 0.80 |
| C_japonicum_15 | 724 | 724032 | 0.79 | 0.74 |
| C_x_alaskanum_43 | 733 | 719207 | 0.80 | 0.73 |
| C_californicum_03 | 739 | 751130 | 0.81 | 0.77 |
| C_singchii_trp | 743 | 755455 | 0.81 | 0.77 |
| C_lentiginosum_trp | 748 | 750604 | 0.82 | 0.77 |
| C_sichuanense_trp | 757 | 771902 | 0.83 | 0.79 |
| C_bardolphianum_trp | 760 | 819755 | 0.83 | 0.84 |
| C_plectrochilon_27 | 760 | 769505 | 0.83 | 0.79 |
| C_forrestii_trp | 764 | 819863 | 0.84 | 0.84 |
| C_micranthum_trp | 765 | 811088 | 0.84 | 0.83 |
| C_fargesii_trp | 770 | 830654 | 0.84 | 0.85 |
| C_irapeanum_14 | 770 | 803092 | 0.84 | 0.82 |
| C_margaritaceum_trp | 773 | 839348 | 0.85 | 0.86 |
| C_lichiangense_trp | 777 | 845618 | 0.85 | 0.86 |
| C_formosanum_trp | 782 | 836789 | 0.86 | 0.85 |
| C_flavum_trp | 784 | 845858 | 0.86 | 0.86 |
| C_acaule_trp | 795 | 837286 | 0.87 | 0.86 |
| Paphiopedilum_callosum_trp | 820 | 827313 | 0.90 | 0.85 |
| Apostasia_shenzhenica_gen | 838 | 819269 | 0.92 | 0.84 |
| Vanilla_planifolia_gen | 840 | 826307 | 0.92 | 0.84 |
| Selenipedium_aequinoctiale_trp | 851 | 883803 | 0.93 | 0.90 |
| Paphiopedilum_hirsutissimum_trp | 853 | 891870 | 0.93 | 0.91 |
| Paphiopedilum_concolor_trp | 862 | 901932 | 0.94 | 0.92 |

| | | | | |
|---|---|---|---|---|
| Mexipedium_xerophyticum_trp | 871 | 916935 | 0.95 | 0.94 |
| Phragmipedium_lindleyanum_trp | 875 | 925251 | 0.96 | 0.95 |
| Vanilla_shenzhenica_trp | 875 | 849137 | 0.96 | 0.87 |
| Phalaenopsis_equestris_gen | 898 | 928175 | 0.98 | 0.95 |
| Dendrobium_catenatum_gen | 900 | 922223 | 0.99 | 0.94 |

[a] Taxon ID is comprised of (a) the taxon name followed by the assigned Lab No. for samples collected from herbarium M, the Botanical Collection at Oberhof associated with the BGM or provided by the Kew Royal Botanical Gardens, or (b) the taxon name followed by the sequence type (trp = transcriptome, gen = genome, gks = genome skimming) for publicly available orchid sequences. Samples whose taxon ID starts with "C_" belong to the genus *Cypripedium*. See Supplementary Data Tables S2, S3, and S4 for more details.

Table S8: Final occupancy statistics of the 80 orthologous chloroplast loci and characters (bp) per specimen following the concatenation step.

| **Taxon ID** [a] | **# of orthologs** | **# of characters** | **% of orthologs** | **% of characters** |
|---|---|---|---|---|
| Cypripedium_himalaicum_65 | 39 | 32205 | 0.49 | 0.47 |
| Cypripedium_parviflorum_var_parviflorum_75 | 43 | 44406 | 0.54 | 0.65 |
| Cypripedium_froschii_11 | 45 | 46698 | 0.56 | 0.68 |
| Cypripedium_formosanum_09 | 47 | 46758 | 0.59 | 0.68 |
| Cypripedium_acaule_48 | 56 | 45900 | 0.70 | 0.67 |
| Cypripedium_franchetii_10 | 57 | 52008 | 0.71 | 0.76 |
| Cypripedium_macranthos_var_taiwanianum_46 | 58 | 52407 | 0.73 | 0.76 |
| Cypripedium_debile_77 | 58 | 51288 | 0.73 | 0.75 |
| Cypripedium_calceolus_var_pubescens_69 | 60 | 50196 | 0.75 | 0.73 |
| Cypripedium_wardii_38 | 62 | 51981 | 0.78 | 0.76 |
| Cypripedium_x_ventricosum_34 | 62 | 54819 | 0.78 | 0.80 |
| Cypripedium_acaule_74 | 62 | 55635 | 0.78 | 0.81 |
| Cypripedium_himalaicum_45 | 63 | 56079 | 0.79 | 0.81 |
| Cypripedium_calceolus_44 | 63 | 55872 | 0.79 | 0.81 |
| Vanilla_shenzhenica_trp | 63 | 33906 | 0.79 | 0.49 |
| Cypripedium_parviflorum_var_makasin_24 | 64 | 57894 | 0.80 | 0.84 |
| Cypripedium_x_columbianum_33 | 64 | 54978 | 0.80 | 0.80 |
| Cypripedium_japonicum_15 | 65 | 56043 | 0.81 | 0.81 |
| Cypripedium_guttatum_12 | 65 | 58968 | 0.81 | 0.86 |
| Cypripedium_singchii_trp | 66 | 44634 | 0.83 | 0.65 |
| Cypripedium_segawai_28 | 66 | 57753 | 0.83 | 0.84 |
| Cypripedium_macranthos_var_rebunense_20 | 67 | 59925 | 0.84 | 0.87 |
| Cypripedium_parviflorum_var_pubescens_25 | 67 | 59844 | 0.84 | 0.87 |
| Vanilla_planifolia_NC026778 | 67 | 55149 | 0.84 | 0.80 |
| Cypripedium_reginae_var_alba_41 | 68 | 58578 | 0.85 | 0.85 |

| | | | | |
|---|---|---|---|---|
| Cypripedium_macranthos_var_alba_18 | 68 | 60402 | 0.85 | 0.88 |
| Cypripedium_flavum_08 | 68 | 58746 | 0.85 | 0.85 |
| Cypripedium_irapeanum_14 | 68 | 57177 | 0.85 | 0.83 |
| Paphiopedilum_hirsutissimum_NC050871 | 70 | 60186 | 0.88 | 0.87 |
| Paphiopedilum_callosum_NC069960 | 70 | 59376 | 0.88 | 0.86 |
| Cypripedium_tibeticum_30 | 70 | 60051 | 0.88 | 0.87 |
| Mexipedium_xerophyticum_NC069868 | 70 | 59997 | 0.88 | 0.87 |
| Cypripedium_yunnanense_32 | 71 | 61008 | 0.89 | 0.89 |
| Cypripedium_parviflorum_var_pubescens_f_planipetalum_26 | 72 | 63021 | 0.90 | 0.92 |
| Cypripedium_cordigerum_05 | 72 | 63546 | 0.90 | 0.92 |
| Cypripedium_kentuckiense_16 | 72 | 62937 | 0.90 | 0.91 |
| Cypripedium_shanxiense_29 | 72 | 60678 | 0.90 | 0.88 |
| Phragmipedium_lindleyanum_trp | 73 | 59802 | 0.91 | 0.87 |
| Cypripedium_sichuanense_trp | 73 | 48675 | 0.91 | 0.71 |
| Phalaenopsis_equestris_NC017609 | 73 | 60312 | 0.91 | 0.88 |
| Cypripedium_macranthos_var_hotei-atsumorianum_19 | 73 | 63402 | 0.91 | 0.92 |
| Cypripedium_micranthum_22 | 74 | 66276 | 0.93 | 0.96 |
| Cypripedium_debile_NC063681 | 74 | 62445 | 0.93 | 0.91 |
| Cypripedium_plectrochilon_27 | 74 | 64959 | 0.93 | 0.94 |
| Cypripedium_fasciolatum_07 | 74 | 61299 | 0.93 | 0.89 |
| Cypripedium_fargesii_trp | 74 | 49977 | 0.93 | 0.73 |
| Cypripedium_macranthos_var_speciosum_21 | 74 | 63093 | 0.93 | 0.92 |
| Paphiopedilum_concolor_NC069964 | 74 | 61983 | 0.93 | 0.90 |
| Cypripedium_reginae_40 | 74 | 62031 | 0.93 | 0.90 |
| Cypripedium_palangshanense_NC063680 | 75 | 62778 | 0.94 | 0.91 |
| Cypripedium_amesianum_35 | 75 | 61542 | 0.94 | 0.89 |
| Cypripedium_calcicola_02 | 75 | 62193 | 0.94 | 0.90 |
| Cypripedium_farreri_06 | 75 | 65538 | 0.94 | 0.95 |
| Cypripedium_forrestii_trp | 75 | 47823 | 0.94 | 0.69 |
| Cypripedium_micranthum_trp | 76 | 52704 | 0.95 | 0.77 |
| Cypripedium_calceolus_var_parviflorum_53 | 76 | 60942 | 0.95 | 0.89 |
| Cypripedium_candidum_04 | 76 | 64689 | 0.95 | 0.94 |
| Cypripedium_lentiginosum_trp | 76 | 51645 | 0.95 | 0.75 |
| Apostasia_shenzhenica_NC039812 | 76 | 63060 | 0.95 | 0.92 |
| Cypripedium_margaritaceum_trp | 76 | 57273 | 0.95 | 0.83 |
| Dendrobium_catenatum_NC037361 | 77 | 62532 | 0.96 | 0.91 |
| Cypripedium_bardolphianum_trp | 77 | 56571 | 0.96 | 0.82 |
| Cypripedium_lichiangense_trp | 77 | 56112 | 0.96 | 0.82 |
| Cypripedium_flavum_trp | 77 | 54714 | 0.96 | 0.79 |
| Cypripedium_formosanum_trp | 77 | 51408 | 0.96 | 0.75 |
| Selenipedium_aequinoctiale_trp | 77 | 58287 | 0.96 | 0.85 |
| Cypripedium_montanum_23 | 77 | 66024 | 0.96 | 0.96 |
| Cypripedium_passerinum_42 | 78 | 64896 | 0.98 | 0.94 |
| Cypripedium_subtropicum_76 | 78 | 64956 | 0.98 | 0.94 |

| | | | | |
|---|---|---|---|---|
| Cypripedium_macranthos_var_macranthos_17 | 78 | 65361 | 0.98 | 0.95 |
| Cypripedium_californicum_57 | 79 | 66963 | 0.99 | 0.97 |
| Cypripedium_flavum_OM066275 | 79 | 65835 | 0.99 | 0.96 |
| Cypripedium_bardolphianum_83 | 79 | 67968 | 0.99 | 0.99 |
| Cypripedium_x_ventricosum_OM066286 | 79 | 67641 | 0.99 | 0.98 |
| Cypripedium_formosanum_NC026772 | 79 | 65724 | 0.99 | 0.95 |
| Cypripedium_plectrochilum_OM066284 | 80 | 67509 | 1 | 0.98 |
| Cypripedium_fasciolatum_OM066274 | 80 | 67914 | 1 | 0.99 |
| Cypripedium_fargesii_47 | 80 | 66669 | 1 | 0.97 |
| Cypripedium_lichiangense_NC084419 | 80 | 67374 | 1 | 0.98 |
| Cypripedium_sichuanense_NC084420 | 80 | 66606 | 1 | 0.97 |
| Cypripedium_subtropicum_NC053551 | 80 | 67545 | 1 | 0.98 |
| Cypripedium_lichiangense_87 | 80 | 68130 | 1 | 0.99 |
| Cypripedium_bardolphianum_OL741711 | 80 | 67710 | 1 | 0.98 |
| Cypripedium_x_alaskanum_43 | 80 | 66792 | 1 | 0.97 |
| Cypripedium_yattabeanum_31 | 80 | 67653 | 1 | 0.98 |
| Cypripedium_japonicum_OM066280 | 80 | 67401 | 1 | 0.98 |
| Cypripedium_farreri_OM066273 | 80 | 68025 | 1 | 0.99 |
| Cypripedium_calceolus_NC045400 | 80 | 67764 | 1 | 0.98 |
| Cypripedium_fasciolatum_84 | 80 | 68289 | 1 | 0.99 |
| Cypripedium_fasciculatum_88 | 80 | 67986 | 1 | 0.99 |
| Cypripedium_californicum_03 | 80 | 67698 | 1 | 0.98 |
| Cypripedium_acaule_trp | 80 | 64854 | 1 | 0.94 |
| Cypripedium_fargesii_NC084418 | 80 | 65424 | 1 | 0.95 |
| Cypripedium_henryi_OM066279 | 80 | 67755 | 1 | 0.98 |
| Cypripedium_calceolus_01 | 80 | 65427 | 1 | 0.95 |
| Cypripedium_guttatum_OM066278 | 80 | 67626 | 1 | 0.98 |

[a] Taxon ID is comprised of (a) the taxon name followed by the assigned Lab No. for samples collected from herbarium M, the Botanical Collection at Oberhof associated with the BGM or provided by the Kew Royal Botanical Gardens, (b) the taxon name followed by "trp" for publicly available orchid transcriptomes, or (c) the taxon name followed by the NCBI accession number for publicly available complete or partial chloroplast genome sequences for publicly available orchid sequences (see Supplementary Data Tables S2, S3, and S4 for more details).

Table S9: The $\alpha(x)$ calculations of the anomaly zone test with the corresponding branch numbers and their branch lengths.

| Branch $x$ | | Branch $y$ | | $\alpha(x)$ |
|---|---|---|---|---|
| Nr | length | Nr | length | |
| 17 | 4.598486 | 1 | 2.351982 | -0.402926145 |
| 1 | 2.351982 | 2 | 1.134936 | -0.3796615812450 |
| 2 | 1.134936 | 3 | 0.476353 | -0.3008610478330 |
| 3 | 0.476353 | 4 | 0.164628 | -0.1397532132990 |
| 4 | 0.164628 | 5 | 0.050474 | 0.1408893084410* |
| 5 | 0.050474 | 6 | 0.042781 | 0.6657161410210* |
| 6 | 0.042781 | 7 | 0.0094 | 0.7636745682490* |
| 7 | 0.0094 | 8 | 0.050667 | 1.9122368256400* |
| 8 | 0.050667 | 9 | 2.636816 | 0.6635291712650 |
| 9 | 2.636816 | 10 | 0.22772 | -0.3864359328600 |
| 9 | 2.636816 | 11 | 1.62219 | -0.3864359328600 |
| 12 | 0.184896 | 13 | 0.092595 | 0.1037583598950* |
| 13 | 0.092595 | 14 | 0.074992 | 0.3597101466500* |
| 14 | 0.074992 | 15 | 0.450694 | 0.4566492216810* |
| 6 | 0.042781 | 16 | 2.703795 | 0.76367456824900 |
| 18 | 0.014692 | 19 | 0.271572 | 1.5342160956800* |
| 20 | 0.178749 | 21 | 0.090561 | 0.1143601218400* |
| 21 | 0.090561 | 22 | 0.17481 | 0.3694667607310* |
| 21 | 0.090561 | 23 | 0.112374 | 0.3694667607310* |
| 22 | 0.17481 | 24 | 0.014068 | 0.1214406118860* |
| 24 | 0.014068 | 25 | 0.143974 | 1.5697848549700* |
| 24 | 0.014068 | 26 | 0.016977 | 1.5697848549700* |
| 27 | 0.066726 | 28 | 0.210933 | 0.5145939766250* |
| 29 | 0.101751 | 30 | 0.265228 | 0.3194486719450* |
| 31 | 0.150217 | 34 | 0.09857 | 0.1716905028390* |
| 32 | 0.139148 | 33 | 0.106984 | 0.1985105684870* |
| 34 | 0.09857 | 36 | 0.062525 | 0.3328029445320* |
| 34 | 0.09857 | 35 | 0.031944 | 0.3328029445320* |
| 35 | 0.031944 | 37 | 0.212367 | 0.9516747586560* |
| 37 | 0.212367 | 38 | 0.03756 | 0.0620216394774* |
| 38 | 0.03756 | 39 | 0.247431 | 0.8451225357780* |
| 39 | 0.247431 | 40 | 0.099602 | 0.0188788493410 |
| 40 | 0.099602 | 41 | 0.107745 | 0.3284008018810* |

"*" marks the $\alpha(x)$ of all internode pairs that were found to be in the anomaly zone [i.e., $y < \alpha(x)$]. For branch numbering, refer to Supplementary Data Figure S5.

Table S10: Calculation and comparison of 10 models produced by the PhyloNet analysis testing for up to one hybridization event in the network including *Cypripedium* × *alaskanum*.

| Topology[a] | lnL | Number of tips | Parameters[b] | Loci | Number of hybridizations | Information criteria | | | | | |
|---|---|---|---|---|---|---|---|---|---|---|---|
| | | | | | | AIC | AICc | BIC | deltaAIC | deltaAICc | deltaBIC |
| Network 1 | -556.90194 | 4 | 7 | 535 | 1 | 1127.80387 | 1128.0164 | 1157.77974 | 0 | 0 | 0 |
| Network 2 | -557.48334 | 4 | 7 | 535 | 1 | 1128.96668 | 1129.1792 | 1158.94255 | 1.16280778 | 1.16280778 | 1.16280778 |
| Network 3 | -557.6424 | 4 | 7 | 535 | 1 | 1129.2848 | 1129.49732 | 1159.26067 | 1.48092626 | 1.48092626 | 1.48092626 |
| Network 4 | -557.92218 | 4 | 7 | 535 | 1 | 1129.84437 | 1130.05689 | 1159.82024 | 2.04049583 | 2.04049583 | 2.04049583 |
| Network 5 | -578.93443 | 4 | 7 | 535 | 1 | 1171.86887 | 1172.08139 | 1201.84473 | 44.0649939 | 44.0649939 | 44.0649939 |
| Network 6 | -578.94751 | 4 | 7 | 535 | 1 | 1171.89503 | 1172.10755 | 1201.87089 | 44.0911535 | 44.0911535 | 44.0911535 |
| Network 7 | -578.96529 | 4 | 7 | 535 | 1 | 1171.93058 | 1172.1431 | 1201.90644 | 44.1267021 | 44.1267021 | 44.1267021 |
| Network 8 | -580.32144 | 4 | 7 | 535 | 1 | 1174.64288 | 1174.8554 | 1204.61875 | 46.8390073 | 46.8390073 | 46.8390073 |
| Network 9 | -580.32151 | 4 | 5 | 535 | 0 | 1170.64302 | 1170.75644 | 1192.05436 | 42.8391502 | 42.740048 | 34.2746167 |
| Network 10 | -580.32158 | 4 | 7 | 535 | 1 | 1174.64316 | 1174.85568 | 1204.61903 | 46.8392851 | 46.8392851 | 46.8392851 |

[a]Topology refers to each of the ten networks produced from the Phylonet analysis including *Cypripedium* × *alaskanum*, listed from highest (Network 1) to lowest (Network 10) total log probability (lnL).
[b]The number of parameters was estimated with p = (2n-3)+(2h), where p = number of parameters, n = number of tips, and h = number of hybridizations. The AIC, AICc, BIC scores were calculated according to (Yu et al. 2012).
Guide to deltaAIC and deltaAICc scores: less than 2 indicates there is substantial evidence to support the candidate model (i.e., the candidate model is almost as good as the best model), between 4 and 7 indicates that the candidate model has considerably less support, greater than 10 indicates that there is essentially no support for the candidate model (i.e., it is unlikely to be the best model). Guide for the deltaBIC scores: less than 2 is not worth more than a bare mention, between 2 and 6 indicates that the evidence against the candidate model is positive, between 6 and 10 indicates that the evidence against the candidate model is strong, greater than 10 indicates that the evidence is very strong.

Table S11: Results table of the BioGeoBEARS analysis using nine areas, showing the AIC and AICc scores and weights of each tested model for comparison.

| **Model** | **LnL** | **params** | **d** | **e** | **j** | **x** | **AIC** | **AIC_wt** | **AICc** | **AICc_wt** |
|---|---|---|---|---|---|---|---|---|---|---|
| DEC | -129.59 | 3 | 0.024 | 0.008 | 0.000 | -0.27 | 265.185 | 0.298 | 265.647 | 0.319 |
| DEC+J | -174.85 | 4 | 0.024 | 0.008 | 0.000 | 0.000 | 357.690 | 0.000 | 358.475 | 0.000 |
| DIVALIKE | -129.75 | 3 | 0.026 | 0.003 | 0.000 | -0.27 | 265.492 | 0.256 | 265.953 | 0.274 |
| DIVALIKE+J | -128.21 | 4 | 0.023 | 0.000 | 0.018 | -0.26 | 264.419 | 0.437 | 265.204 | 0.399 |
| BAYAREALIKE | -134.94 | 3 | 0.027 | 0.071 | 0.000 | -0.96 | 275.878 | 0.001 | 276.340 | 0.002 |
| BAYAREALIKE+J | -132.34 | 4 | 0.025 | 0.064 | 0.008 | -1.35 | 272.680 | 0.007 | 273.464 | 0.006 |

Table S12: Results table of the BioGeoBEARS analysis using two areas (New and Old World), showing the AIC and AICc scores and weights of each tested model for comparison.

| **Model** | **LnL** | **params** | **d** | **e** | **j** | **AIC** | **AIC_wt** | **AICc** | **AICc_wt** |
|---|---|---|---|---|---|---|---|---|---|
| DEC | -38.84 | 2 | 0.020 | 0.003 | 0.000 | 81.677 | 0.000 | 81.903 | 0.000 |
| DEC+J | -32.81 | 3 | 0.006 | 0.000 | 0.074 | 71.628 | 0.016 | 72.090 | 0.016 |
| DIVALIKE | -36.61 | 2 | 0.023 | 0.001 | 0.000 | 77.227 | 0.001 | 77.454 | 0.001 |
| DIVALIKE+J | -33.37 | 3 | 0.010 | 0.000 | 0.058 | 72.740 | 0.009 | 73.201 | 0.009 |
| BAYAREALIKE | -49.54 | 2 | 0.010 | 0.010 | 0.000 | 103.074 | 0.000 | 103.300 | 0.000 |
| BAYAREALIKE+J | -28.70 | 3 | 0.002 | 0.000 | 0.079 | 63.391 | 0.974 | 63.852 | 0.974 |

### Supplementary Figures

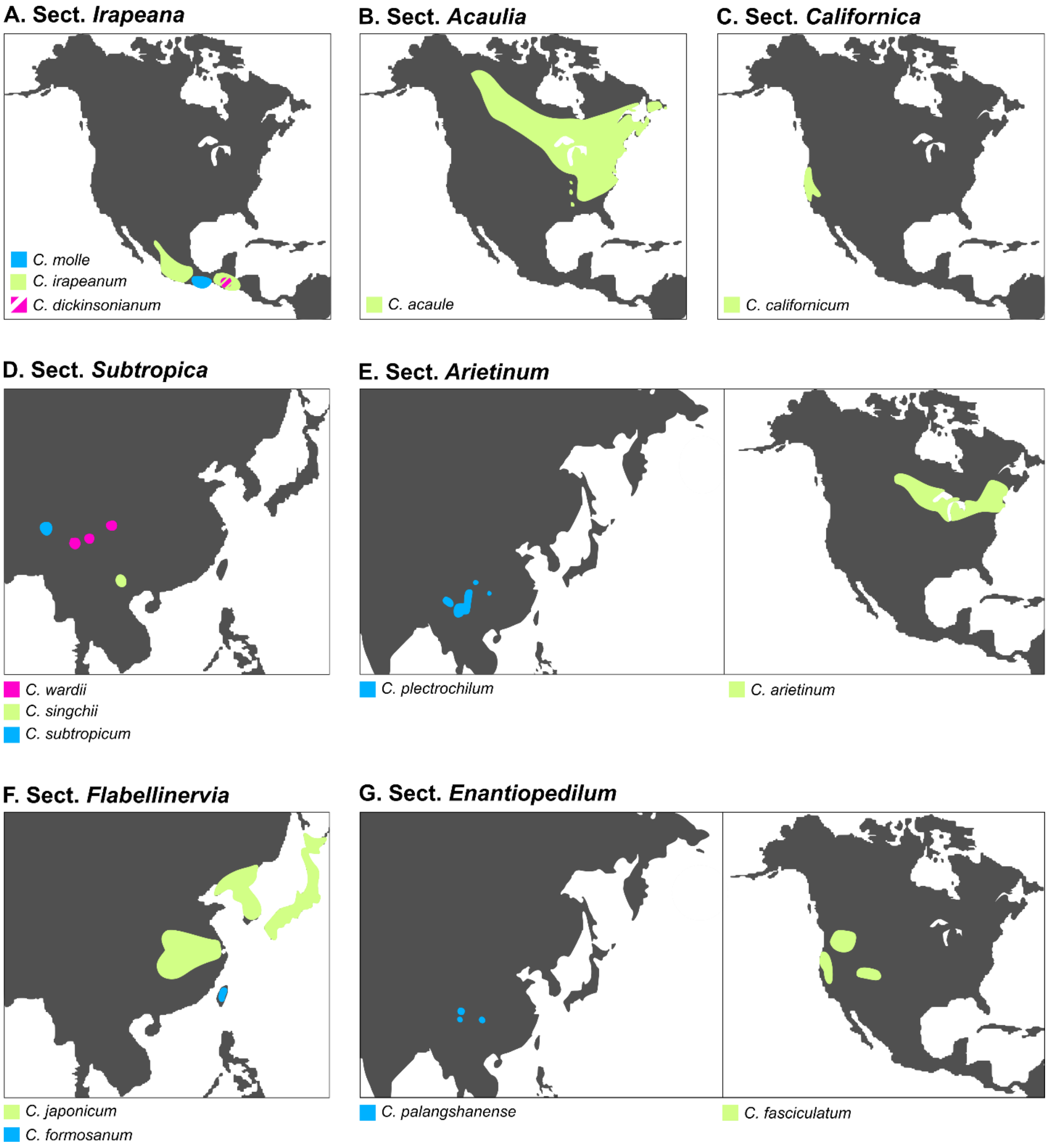

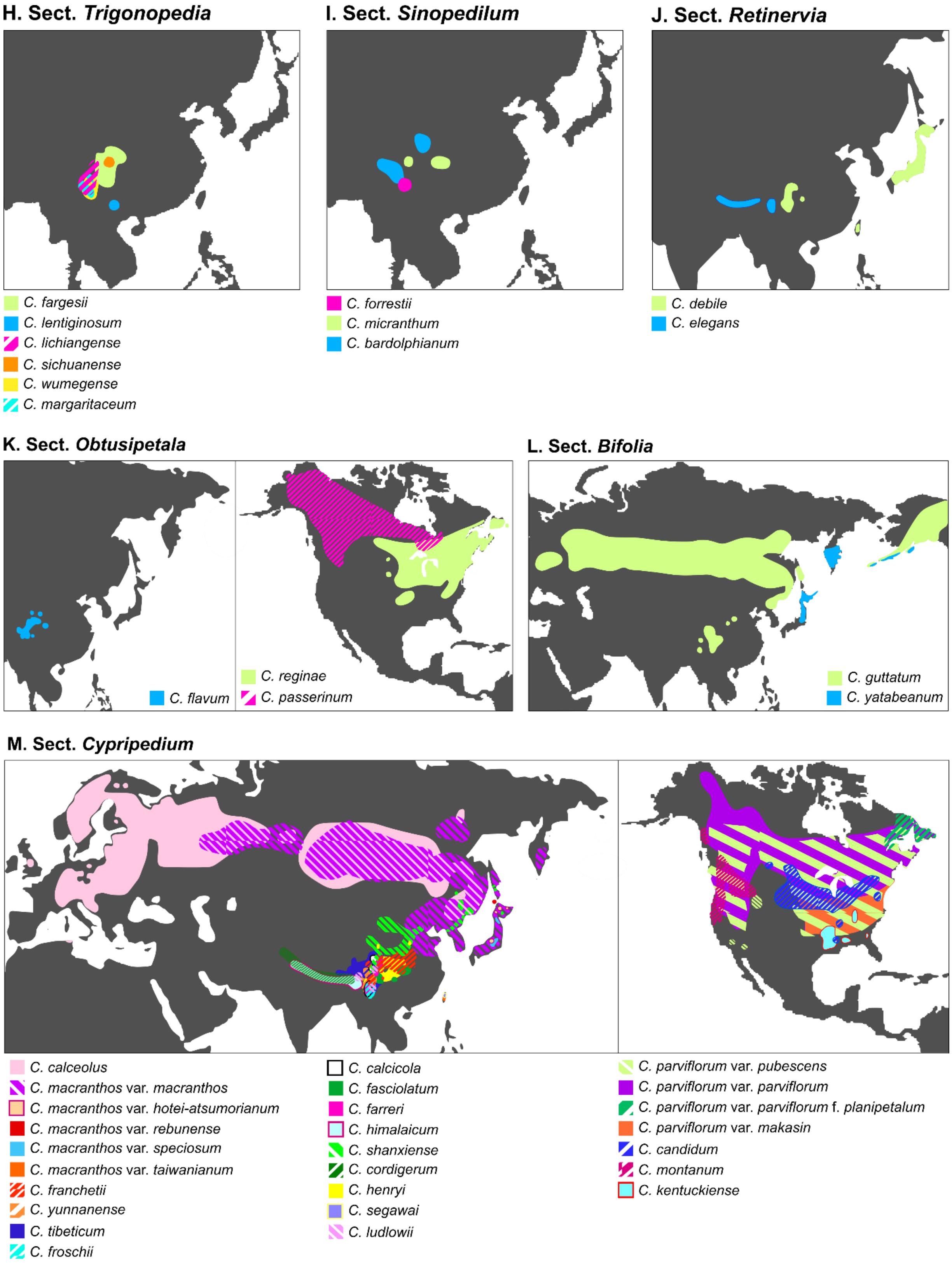

Figure S1: Distribution of *Cypripedium* species per section [classification following Frosch and Cribb (2012); distribution information based on Eccarius (2009), Frosch and Cribb (2012), Chen *et al.* (2013), and Walid *et al.* (2019)].

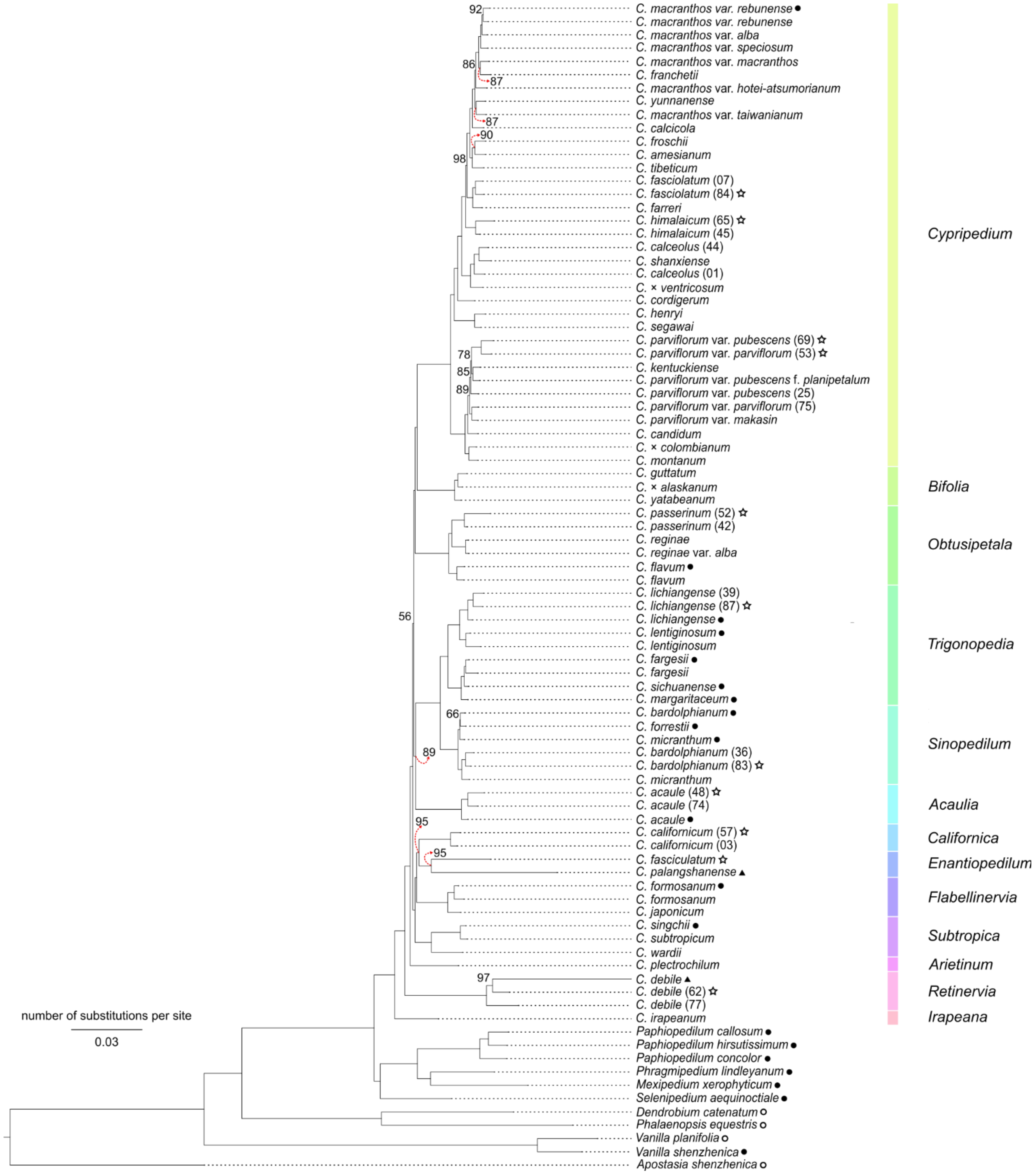

Figure S2: The concatenation-based phylogeny of *Cypripedium*, inferred with 913 nuclear loci using IQ-TREE. Bootstrap support values are shown above or below the branches when <100. Branches are annotated according to their section-level classification following Frosch and Cribb (2012). Tip symbols: filled circles "●" denote transcriptomes, unfilled circles "○" denote genomes, filled triangles "▲" denote genome skimming sequences, and unfilled stars "☆" denote herbarium or old silica-dried specimens. Tips without symbols come from living specimens of the Botanical Collection at Oberhof.

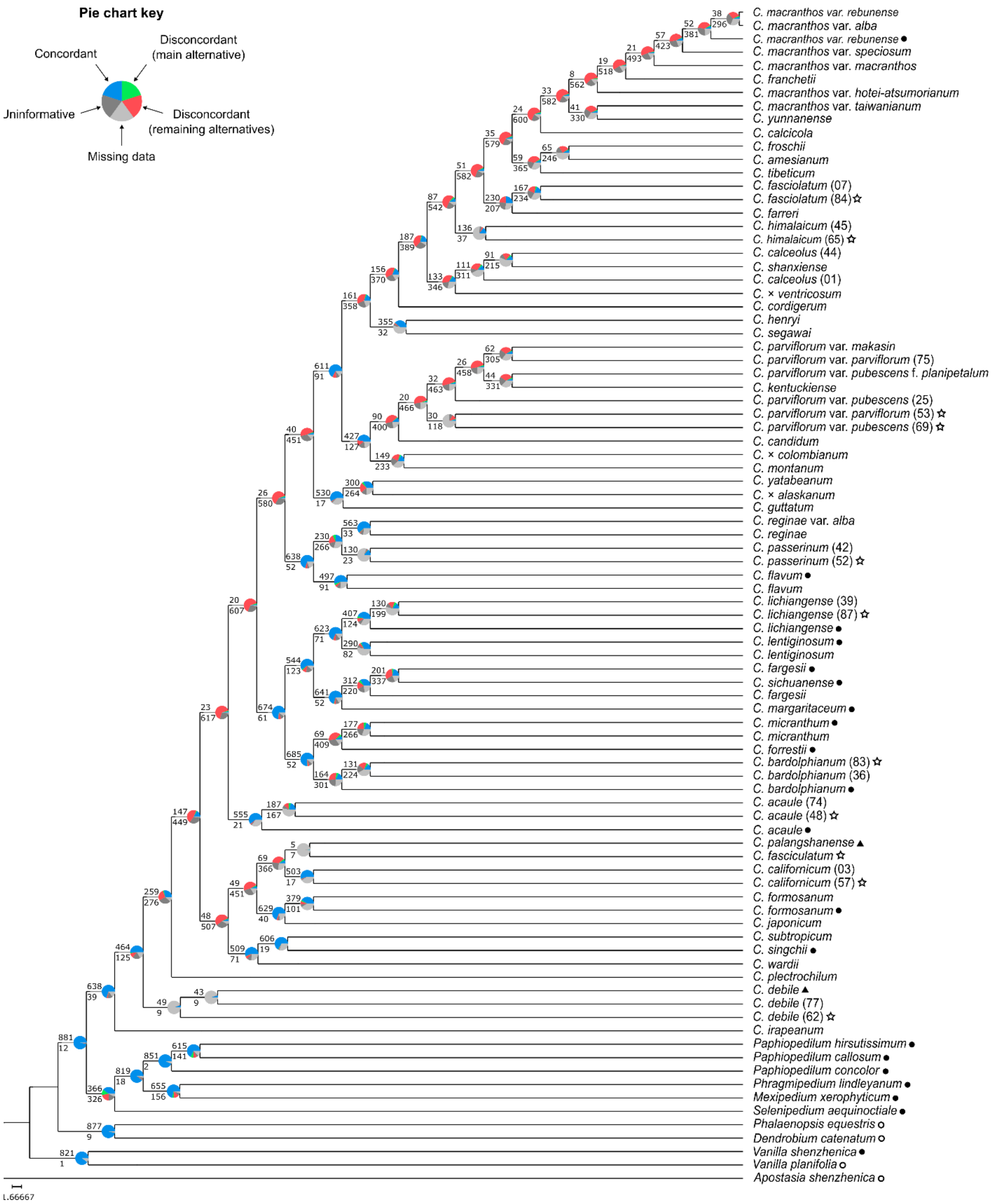

Figure S3: The output nuclear phylogeny of *Cypripedium* from the Phyparts analysis. The pie charts at the nodes indicate the proportions of informative concordant and discordant gene tree topologies (top and bottom numbers on each branch, respectively), along with the proportion of uninformative and missing loci (see pie chart key on the top left). Tip symbols: filled circles "●" denote transcriptomes, unfilled circles "○" denote genomes, filled triangles "▲" denote genome skimming sequences, and unfilled stars "☆" denote herbarium or old silica-dried specimens. Tips without symbols come from living specimens of the Botanical Collection at Oberhof.

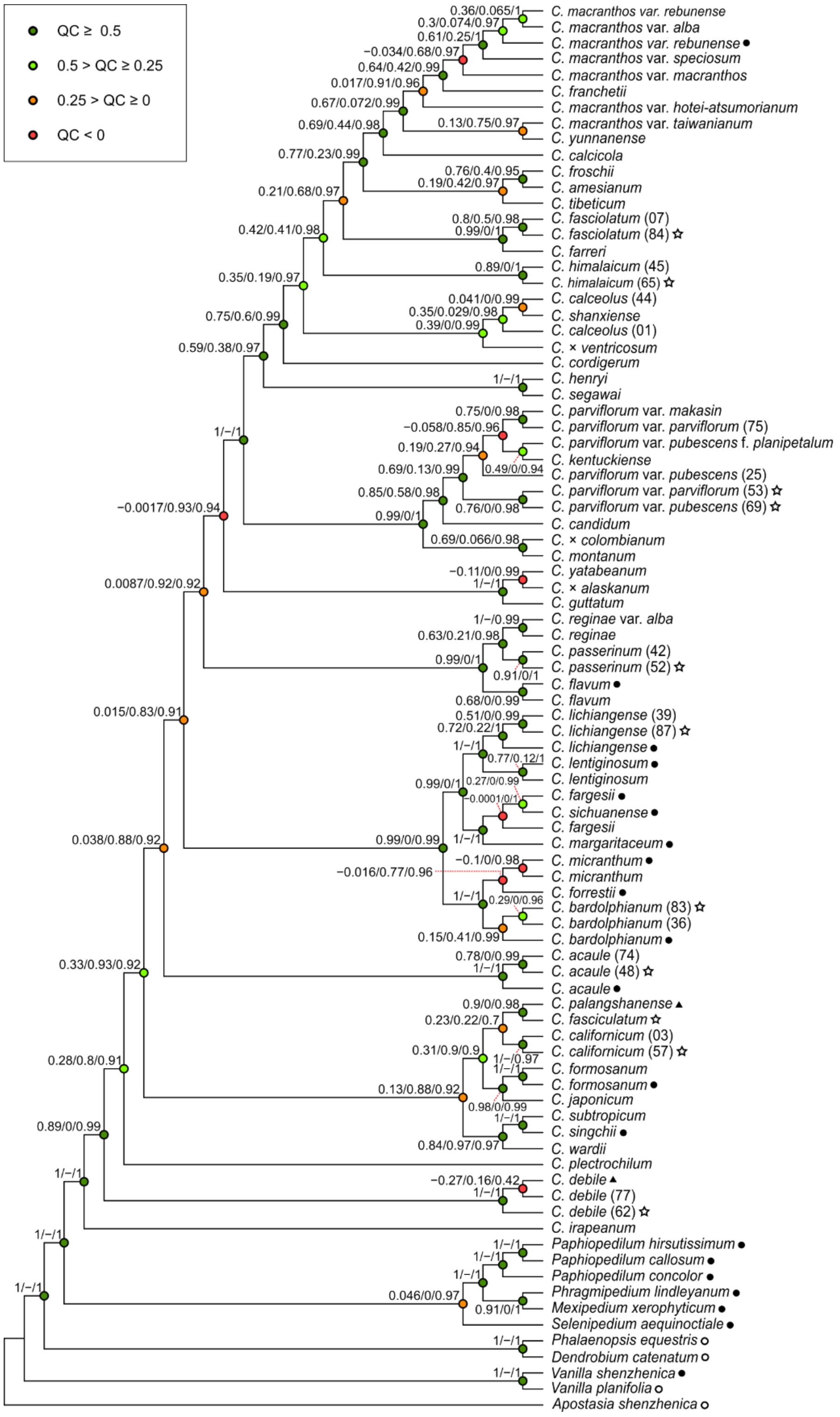

Figure S4: The output nuclear phylogeny of *Cypripedium* from the Quartet Sampling analysis. The nodes are annotated with the quartet support values Quartet Concordance/Quartet Differential/Quartet Informativeness in order (for interpretation, see Pease *et al.*, 2018) and colored based on the Quartet Concordance values (key to colors on the top left). Tip symbols: filled circles "●" denote transcriptomes, unfilled circles "○" denote genomes, filled triangles "▲" denote genome skimming sequences, and unfilled stars "☆" denote herbarium or old silica-dried specimens. Tips without symbols come from living specimens of the Botanical Collection at Oberhof.

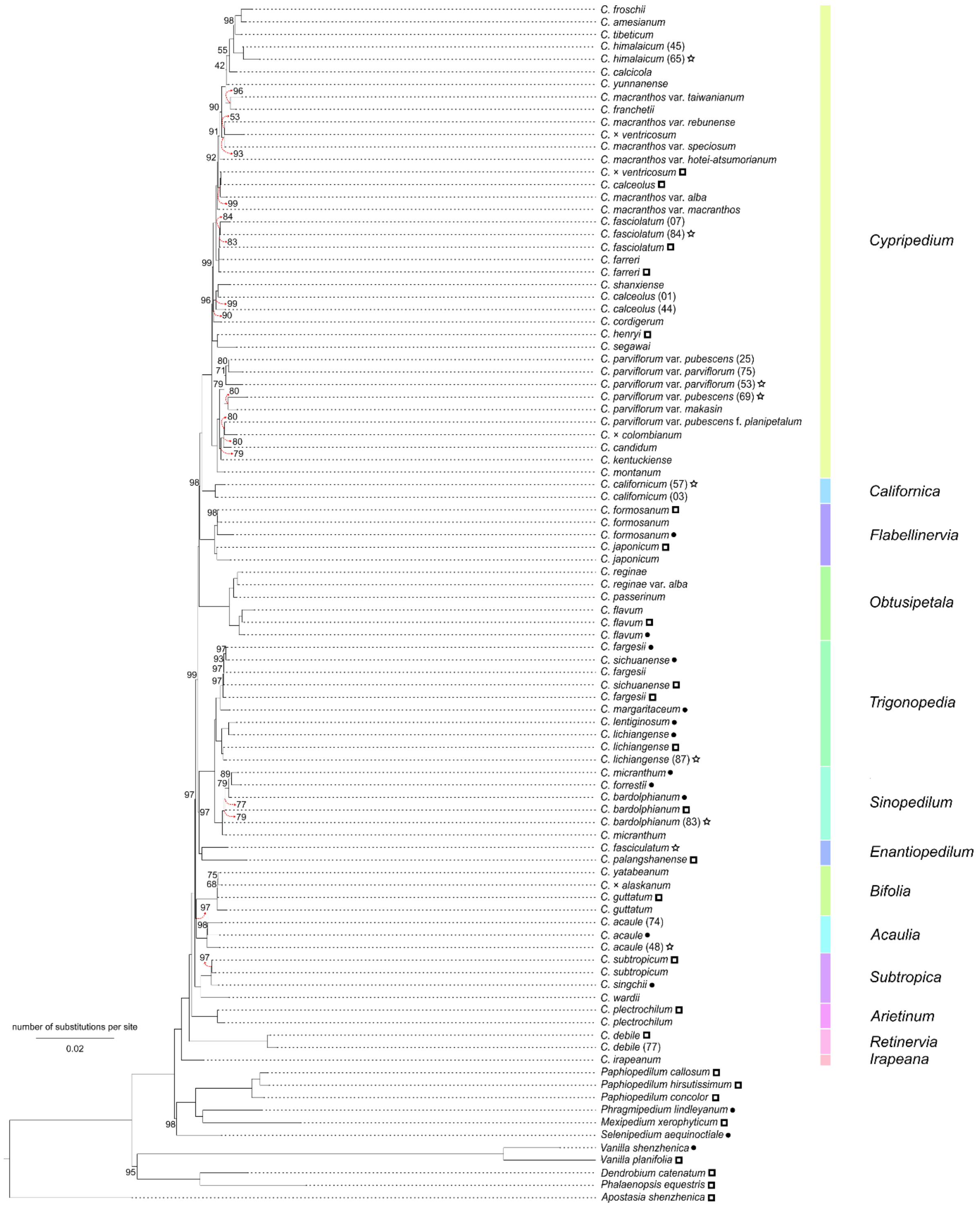

Figure S5: The concatenation-based phylogeny of *Cypripedium*, inferred with 80 chloroplast loci using IQ-TREE. Bootstrap support values are shown above or below the branches when <100. Branches are annotated according to their section-level classification following Frosch and Cribb (2012). Tip symbols: filled circles "●" denote transcriptomes, unfilled squares "□" denote chloroplast genomes, filled triangles "▲" denote genome skimming sequences, and unfilled stars "☆" denote herbarium or old silica-dried specimens. Tips without symbols come from living specimens of the Botanical Collection at Oberhof.

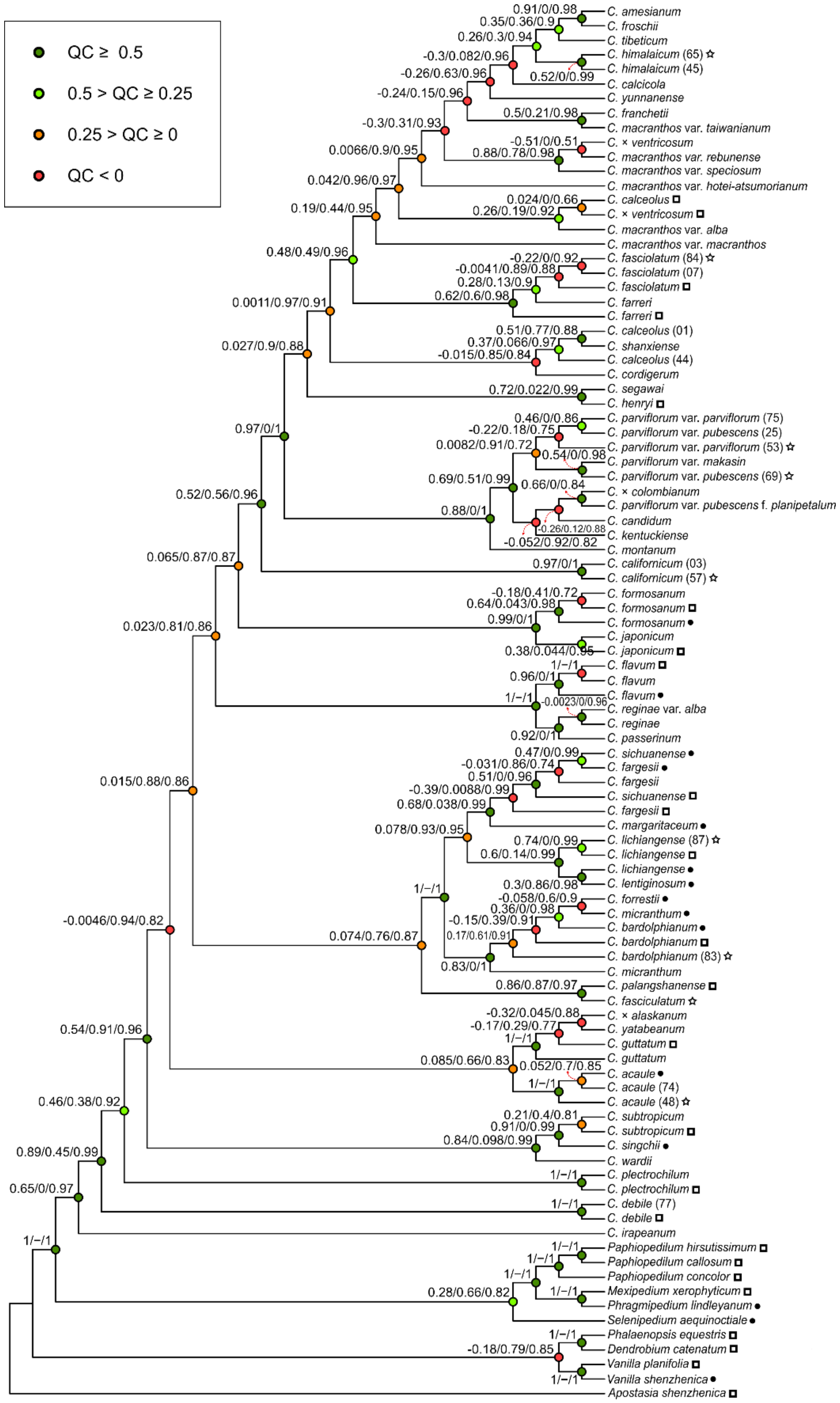

Figure S6: The output chloroplast *Cypripedium* phylogeny from the Quartet Sampling analysis. The nodes are annotated with the quartet support values Quartet Concordance/Quartet Differential/Quartet Informativeness in order (for interpretation, see Pease *et al.*, 2018) and colored based on the Quartet Concordance values (key to colors on the top left). Tip symbols: filled circles "●" denote transcriptomes, unfilled squares "□" denote chloroplast genomes, filled triangles "▲" denote genome skimming sequences, and unfilled stars "☆" denote herbarium or old silica-dried specimens. Tips without symbols come from living specimens of the Botanical Collection at Oberhof.

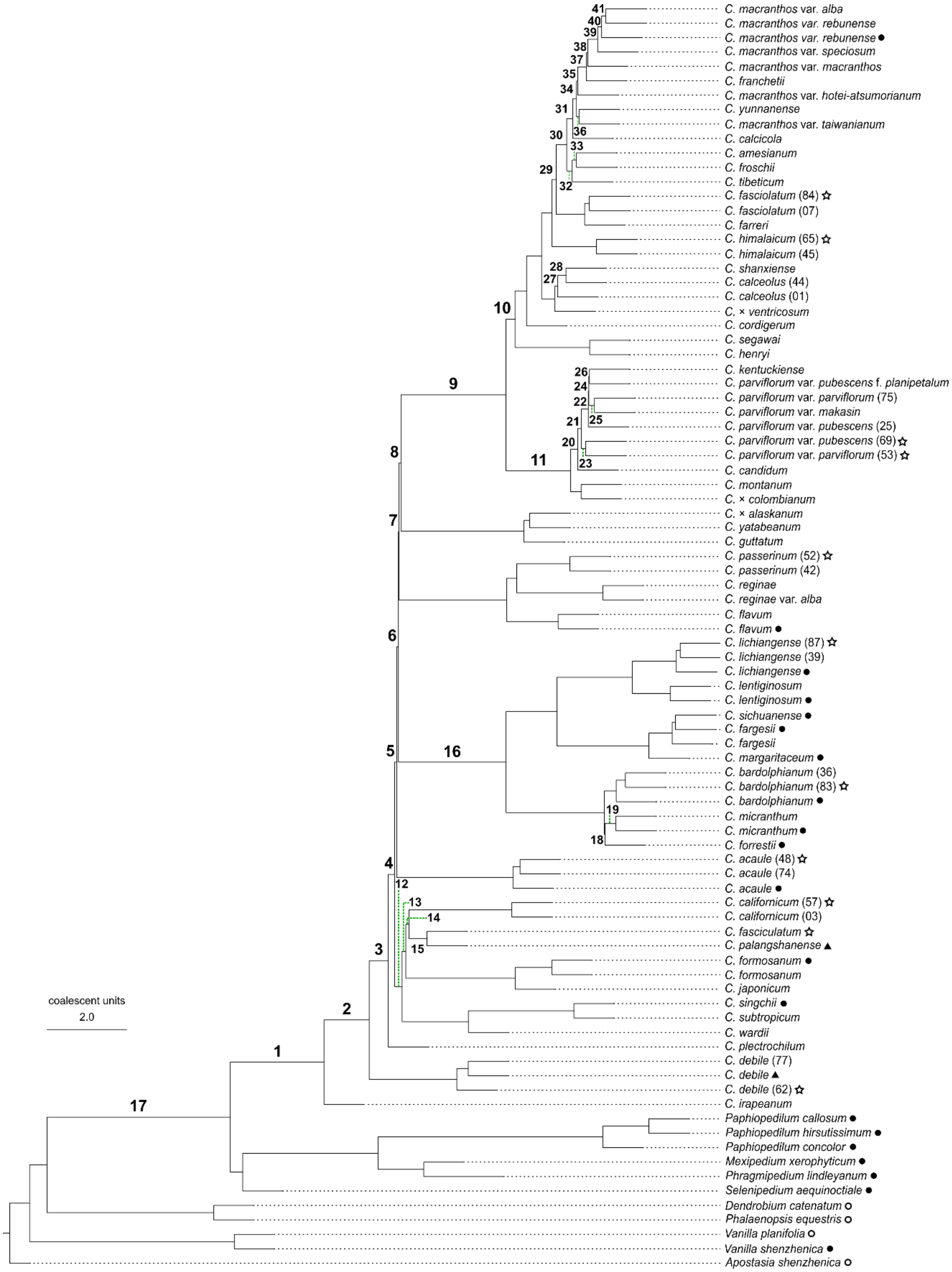

Figure S7: The nuclear ASTRAL phylogeny of *Cypripedium* annotated with the corresponding branch numbers referred to in Supplementary Table S9 for the results of the anomaly zone test calculations and in Figure 3 B. Tip symbols: filled circles "●" denote transcriptomes, unfilled circles "○" denote genomes, filled triangles "▲" denote genome skimming sequences, and unfilled stars "☆" denote herbarium or old silica-dried specimens. Tips without symbols come from living specimens of the Botanical Collection at Oberhof.

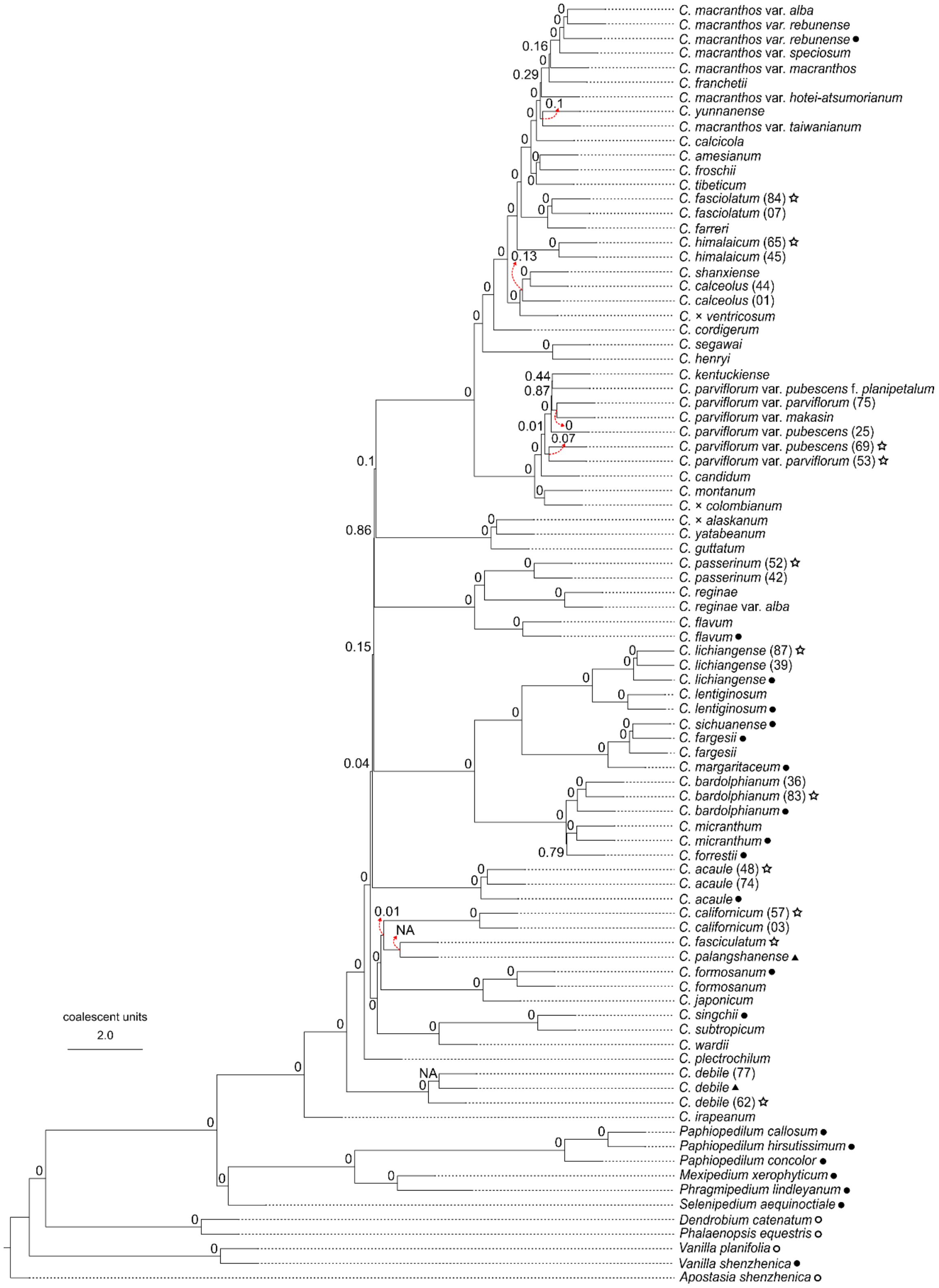

Figure S8: Results of the polytomy test on the nuclear ASTRAL phylogeny of *Cypripedium*. Tip symbols: filled circles "●" denote transcriptomes, unfilled circles "○" denote genomes, filled triangles "▲" denote genome skimming sequences, and unfilled stars "☆" denote herbarium or old silica-dried specimens. Tips without symbols come from living specimens of the Botanical Collection at Oberhof.

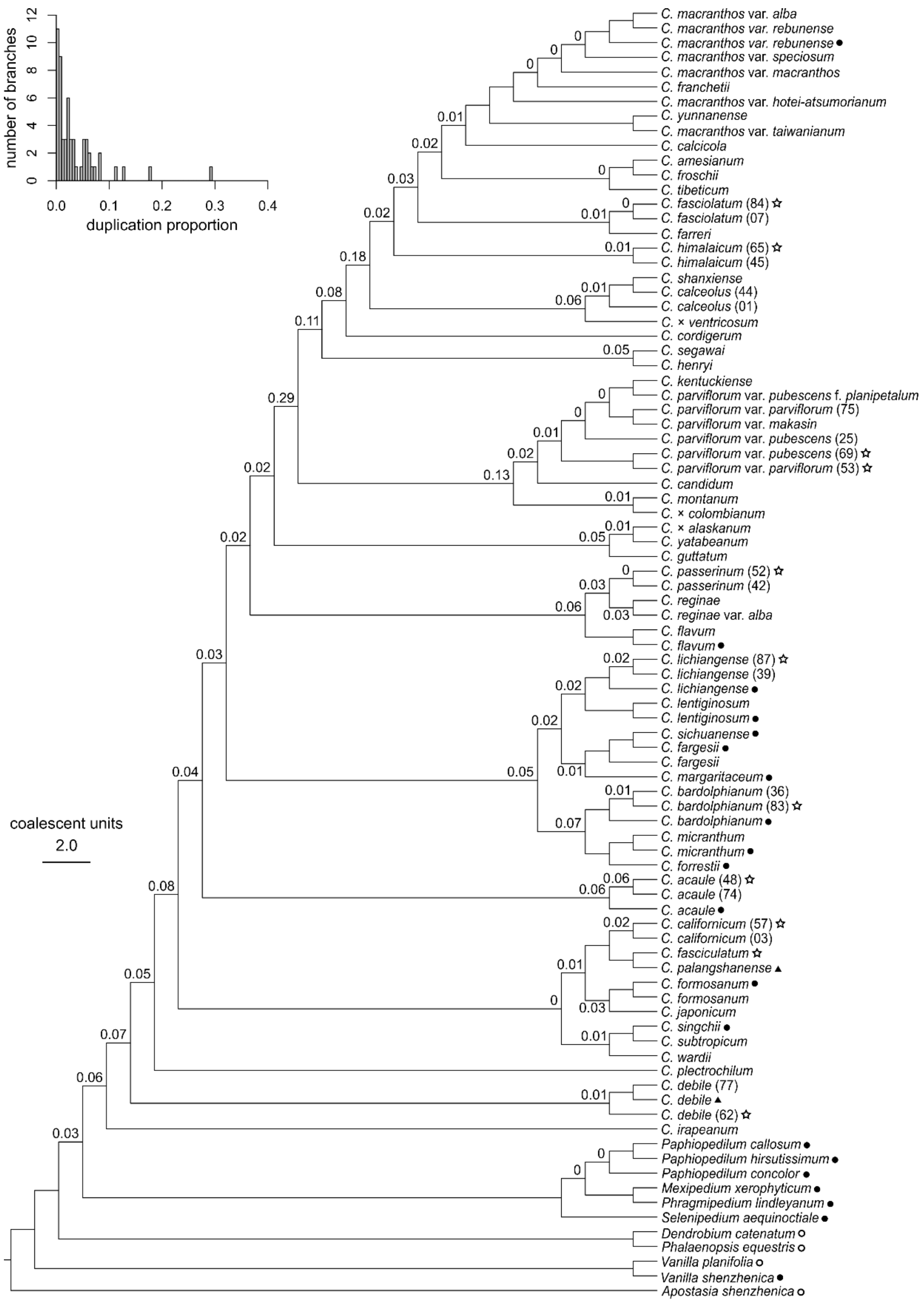

Figure S9: Results of the gene duplication test based on the subclade orthogroup tree topology method implemented with the nuclear ASTRAL phylogeny of *Cypripedium*, using the bootstrap filtering approach. The proportions of duplicated genes are labeled above or below the phylogeny's branches. The number of branches is plotted against the gene duplication proportions on the top left. Branches that did not receive a label did not meet the filtering requirements. Tip symbols: filled circles "●" denote transcriptomes, unfilled circles "○" denote genomes, filled triangles "▲" denote genome skimming sequences, and unfilled stars "☆" denote herbarium or old silica-dried specimens. Tips without symbols come from living specimens of the Botanical Collection at Oberhof.

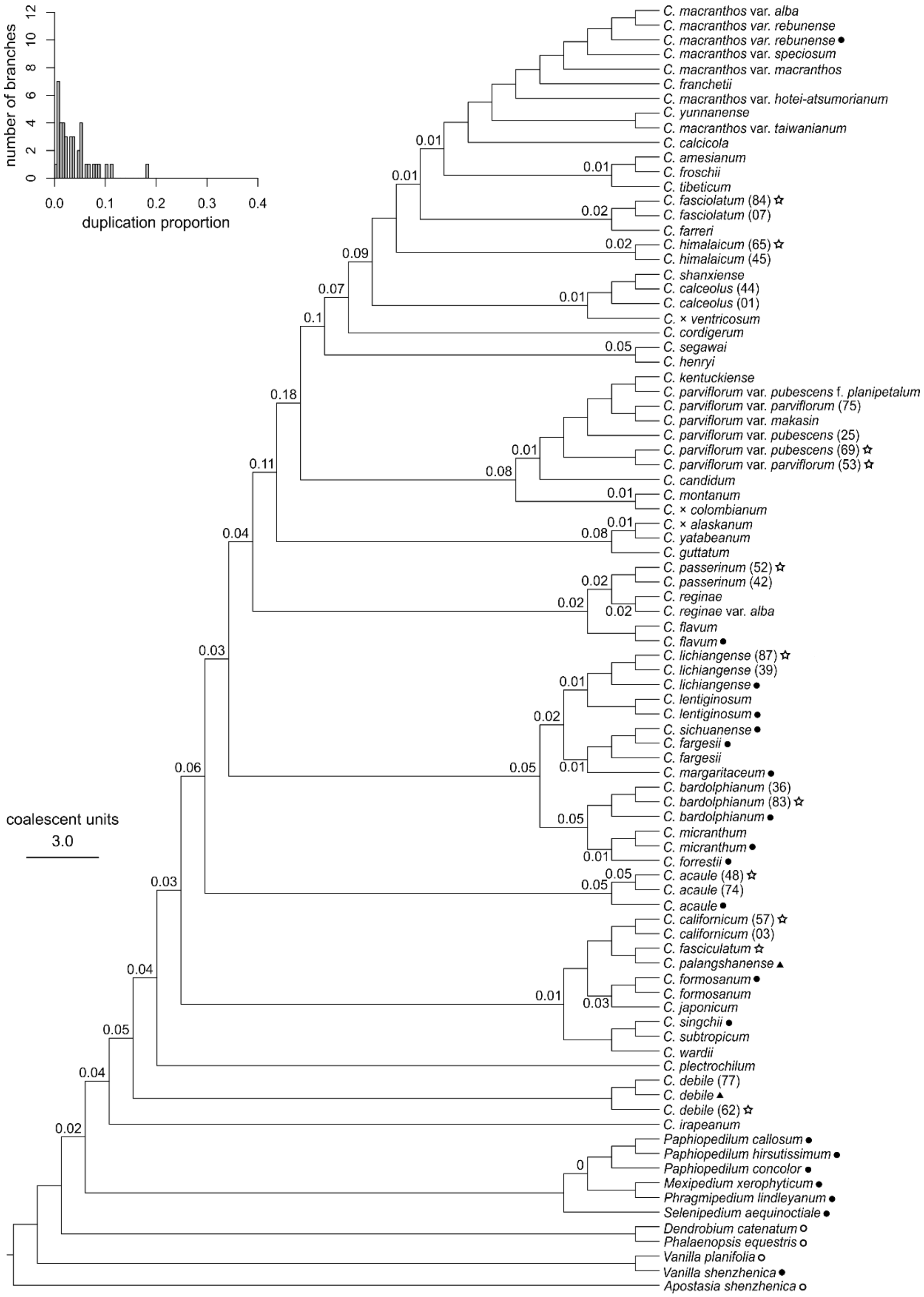

Figure S10: Results of the gene duplication test based on the subclade orthogroup tree topology method implemented with the nuclear ASTRAL phylogeny of *Cypripedium*, using the local topology filtering approach. The proportions of duplicated genes are labeled above or below the phylogeny’s branches. The number of branches is plotted against the gene duplication proportions on the top left. Branches that did not receive a label did not meet the filtering requirements. Tip symbols: filled circles “●” denote transcriptomes, unfilled circles “○” denote genomes, filled triangles “▲” denote genome skimming sequences, and unfilled stars “☆” denote herbarium or old silica-dried specimens. Tips without symbols come from living specimens of the Botanical Collection at Oberhof.

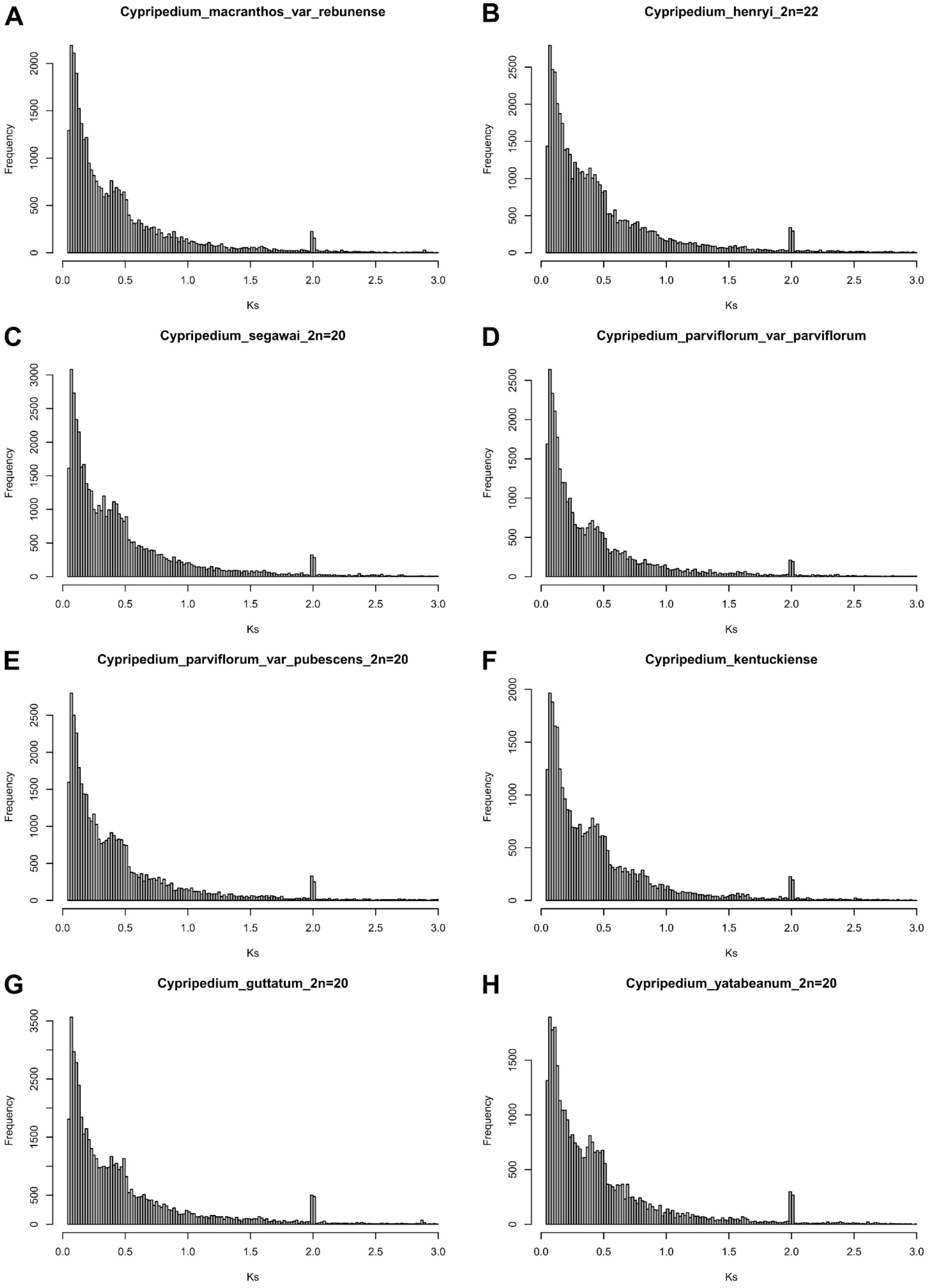
A
Cypripedium_macranthos_var_rebunense
B
Cypripedium_henryi_2n=22
C
Cypripedium_segawai_2n=20
D
Cypripedium_parviflorum_var_parviflorum
E
Cypripedium_parviflorum_var_pubescens_2n=20
F
Cypripedium_kentuckiense
G
Cypripedium_guttatum_2n=20
H
Cypripedium_yatabeanum_2n=20
Frequency
Ks

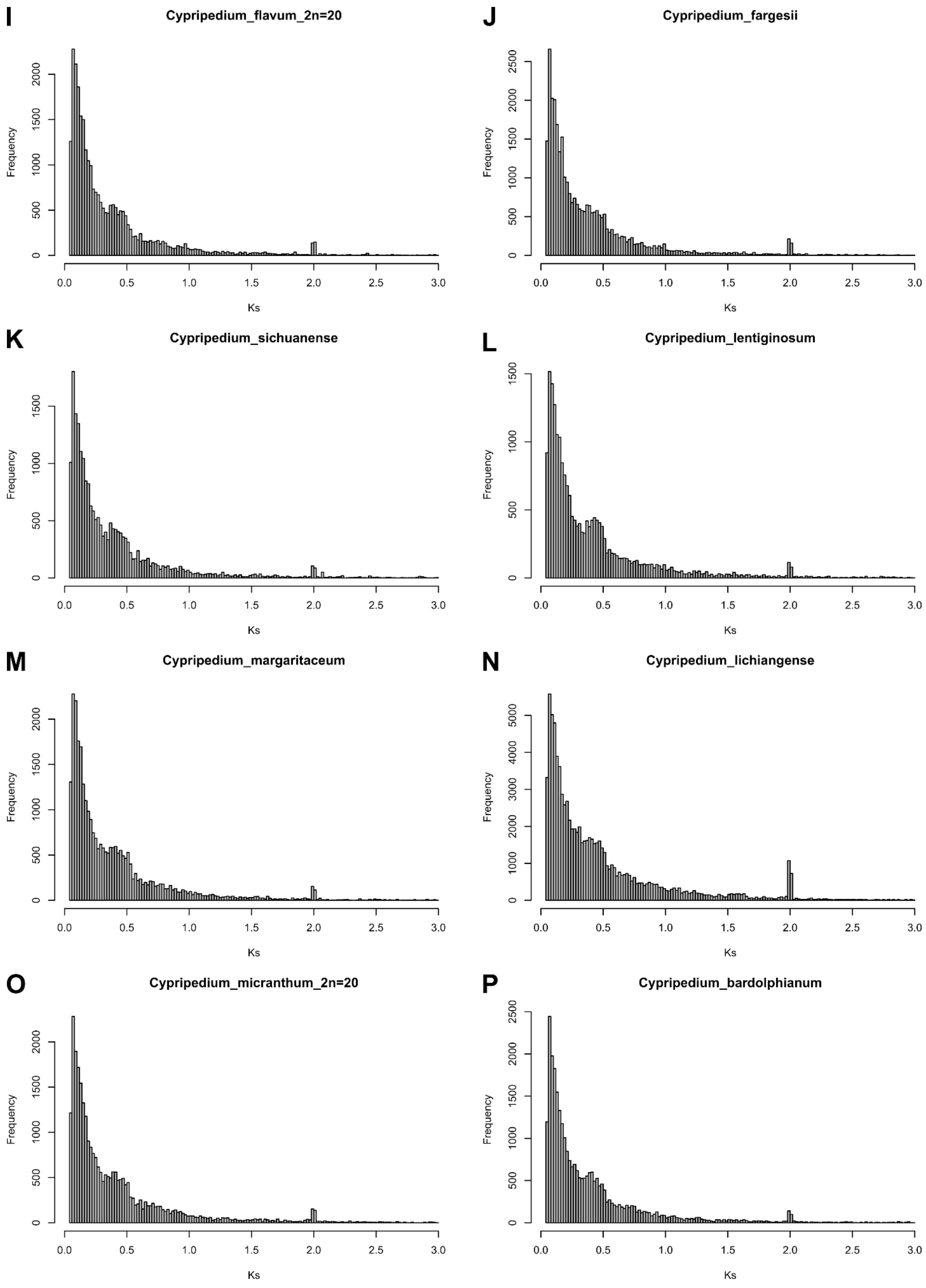
I
Cypripedium_flavum_2n=20
J
Cypripedium_fargesii
K
Cypripedium_sichuanense
L
Cypripedium_lentiginosum
M
Cypripedium_margaritaceum
N
Cypripedium_lichiangense
O
Cypripedium_micranthum_2n=20
P
Cypripedium_bardolphianum
Frequency
Ks
0.0 0.5 1.0 1.5 2.0 2.5 3.0

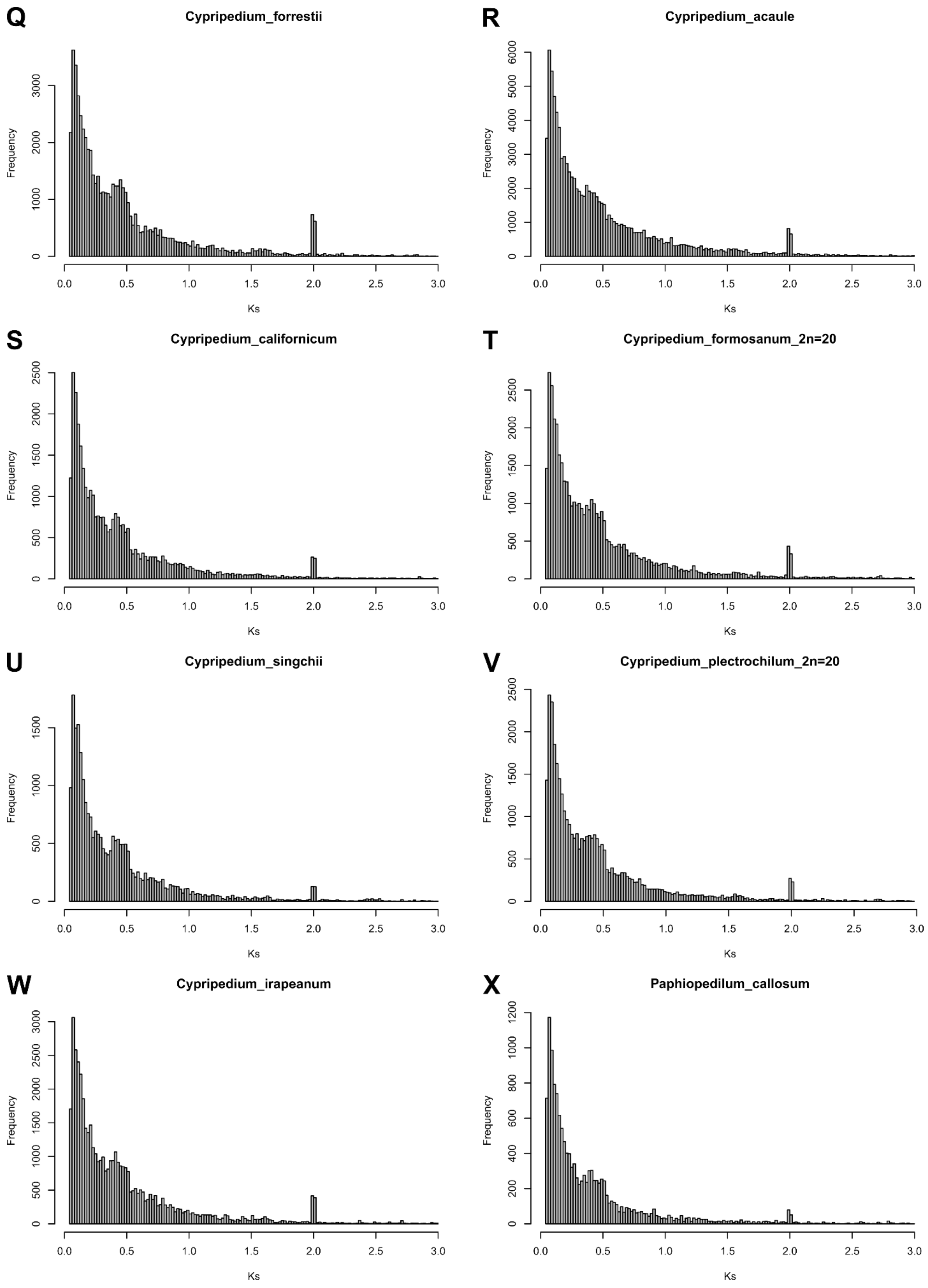
Q
Cypripedium_forrestii
Frequency
Ks
R
Cypripedium_acaule
Frequency
Ks
S
Cypripedium_californicum
Frequency
Ks
T
Cypripedium_formosanum_2n=20
Frequency
Ks
U
Cypripedium_singchii
Frequency
Ks
V
Cypripedium_plectrochilum_2n=20
Frequency
Ks
W
Cypripedium_irapeanum
Frequency
Ks
X
Paphiopedilum_callosum
Frequency
Ks

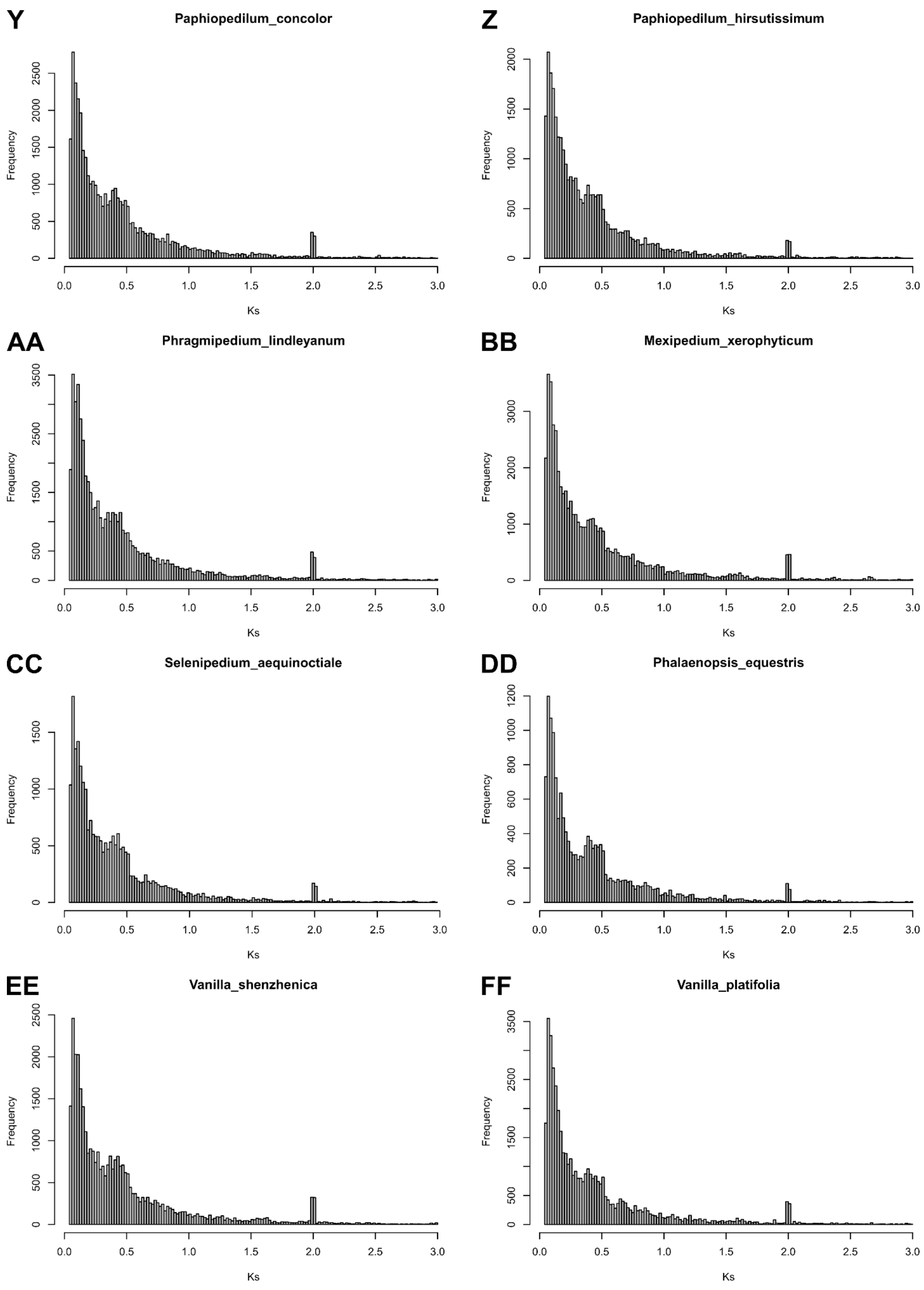
Y
Paphiopedilum_concolor
Z
Paphiopedilum_hirsutissimum
AA
Phragmipedium_lindleyanum
BB
Mexipedium_xerophyticum
CC
Selenipedium_aequinoctiale
DD
Phalaenopsis_equestris
EE
Vanilla_shenzhenica
FF
Vanilla_platifolia
Frequency
Ks

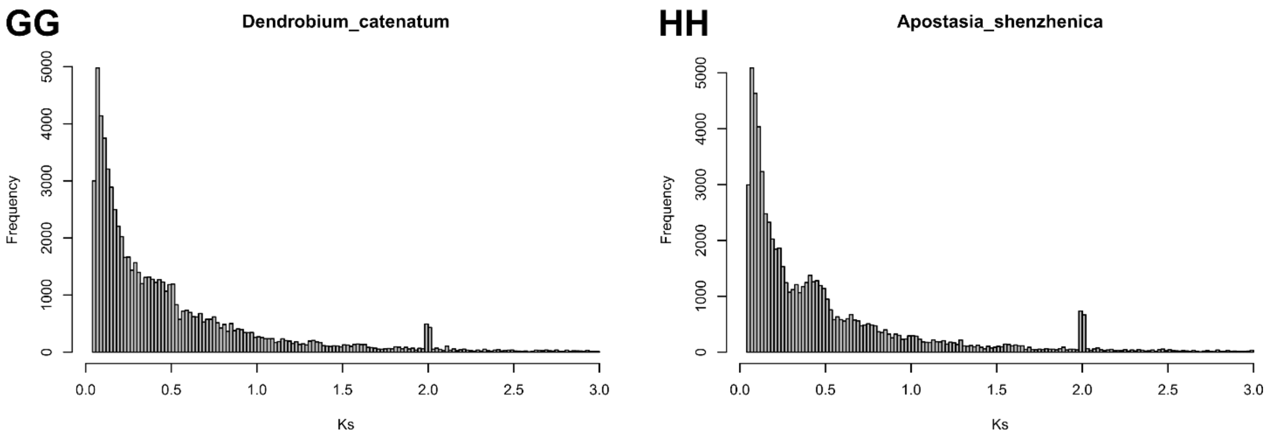

Figure S11: Ks plots for the *Cypripedium* transcriptomes generated in this study and additional orchid transcriptomes obtained from the NCBI (see Supplementary Tables S4 and S6). The usual count of diploid chromosomes is given for some *Cypripedium* species in the figure's title.

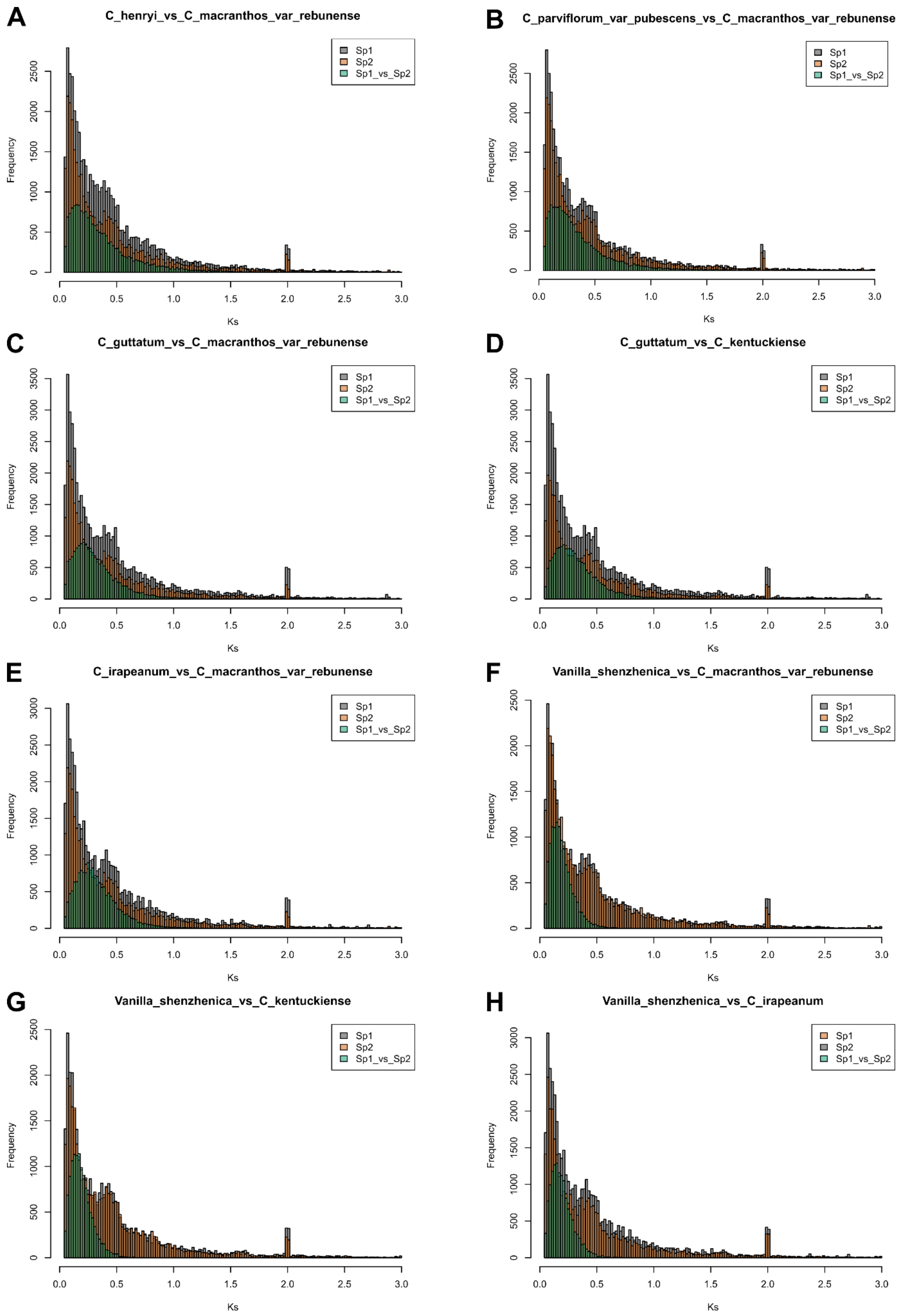
A
C_henryi_vs_C_macranthos_var_rebunense
B
C_parviflorum_var_pubescens_vs_C_macranthos_var_rebunense
C
C_guttatum_vs_C_macranthos_var_rebunense
D
C_guttatum_vs_C_kentuckiense
E
C_irapeanum_vs_C_macranthos_var_rebunense
F
Vanilla_shenzhenica_vs_C_macranthos_var_rebunense
G
Vanilla_shenzhenica_vs_C_kentuckiense
H
Vanilla_shenzhenica_vs_C_irapeanum
Sp1
Sp2
Sp1_vs_Sp2
Frequency
Ks

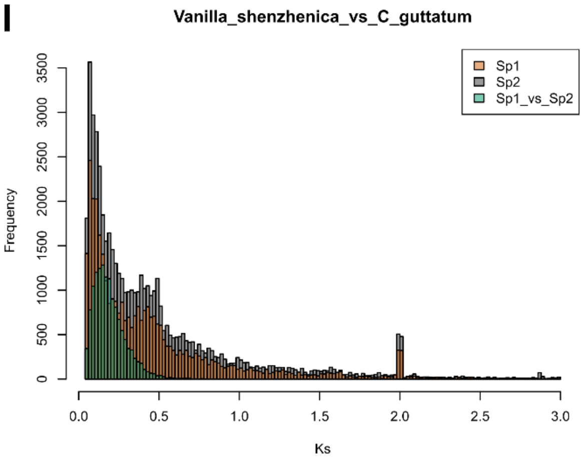

Figure S12: Ks plots comparison between species of the genus *Cypripedium* or between species of the genus *Cypripedium* and other orchids (see Supplementary Tables S4 and S6 for sequence details).

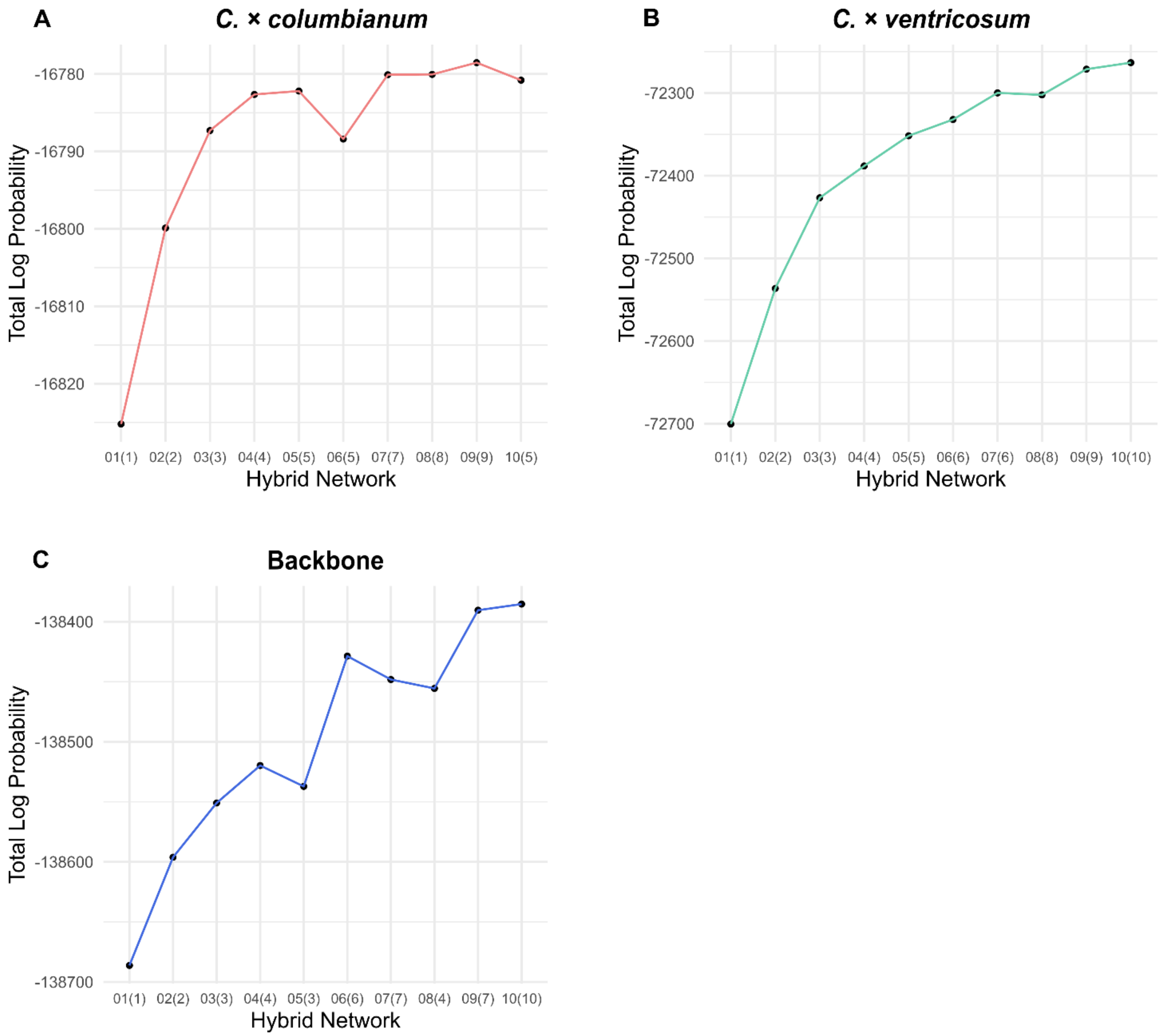

Figure S13: The total log probabilities (y-axis) of the most likely network from each run for the tests including (A) *C. × columbianum*, (B) *C. × ventricosum*, and (C) taxa from all sections to test for hybridizations at the backbone. Network IDs (x-axis): maximum number of hybridization events tested followed by the actual number of produced hybridization events in the parenthesis.

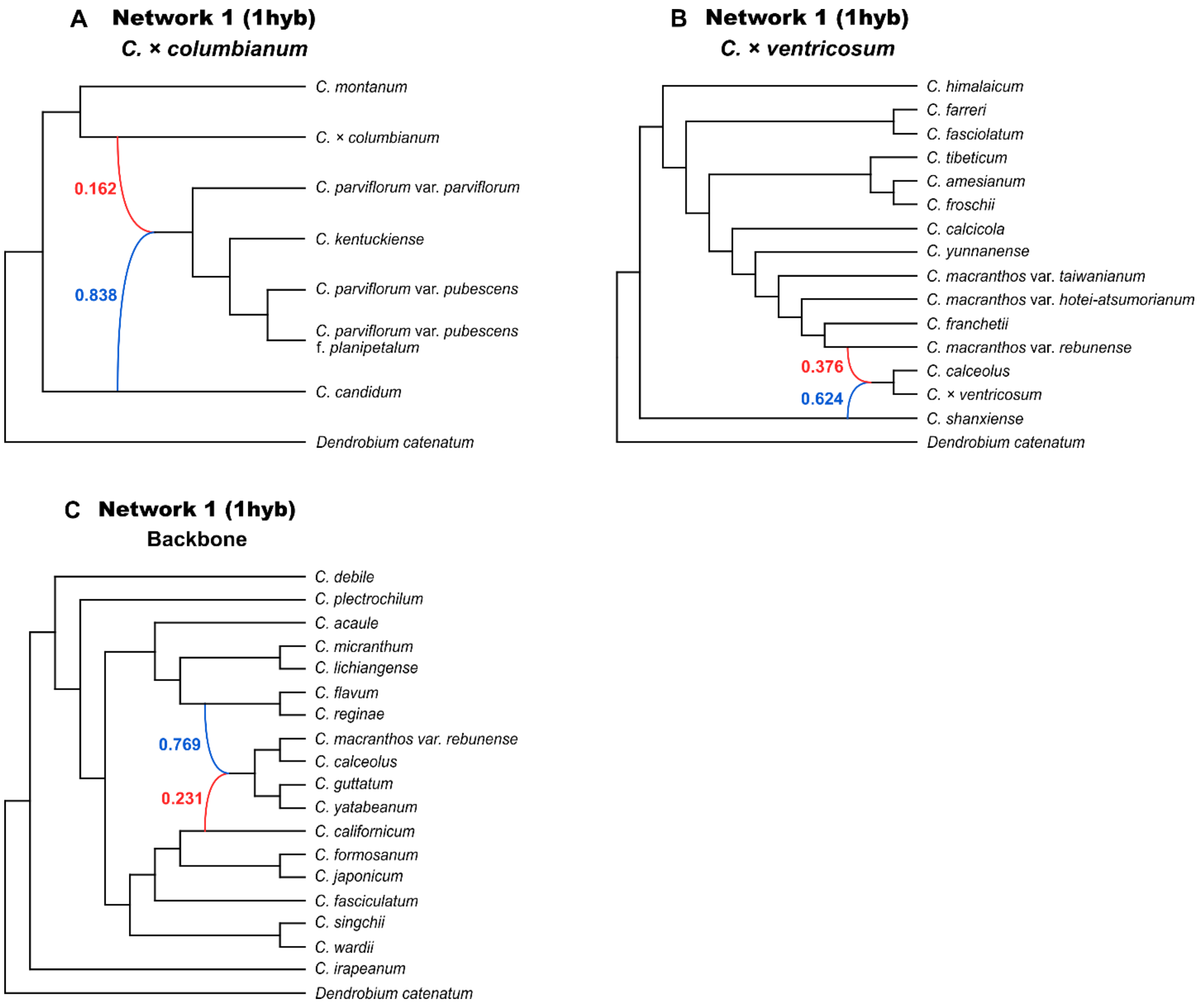

Figure S14: Phylogenetic networks with the highest total log probabilities resulting from the PhyloNet analysis testing for one hybridization event for the extracted subclades with the hybrids (A) *C.* × *columbianum* and (B) *C.* × *ventricosum* or (C) subclades from all sections to test for reticulation in the backbone. The inheritance probabilities are shown for each parent hybrid edge (blue = major hybrid edge; red = minor hybrid edge).

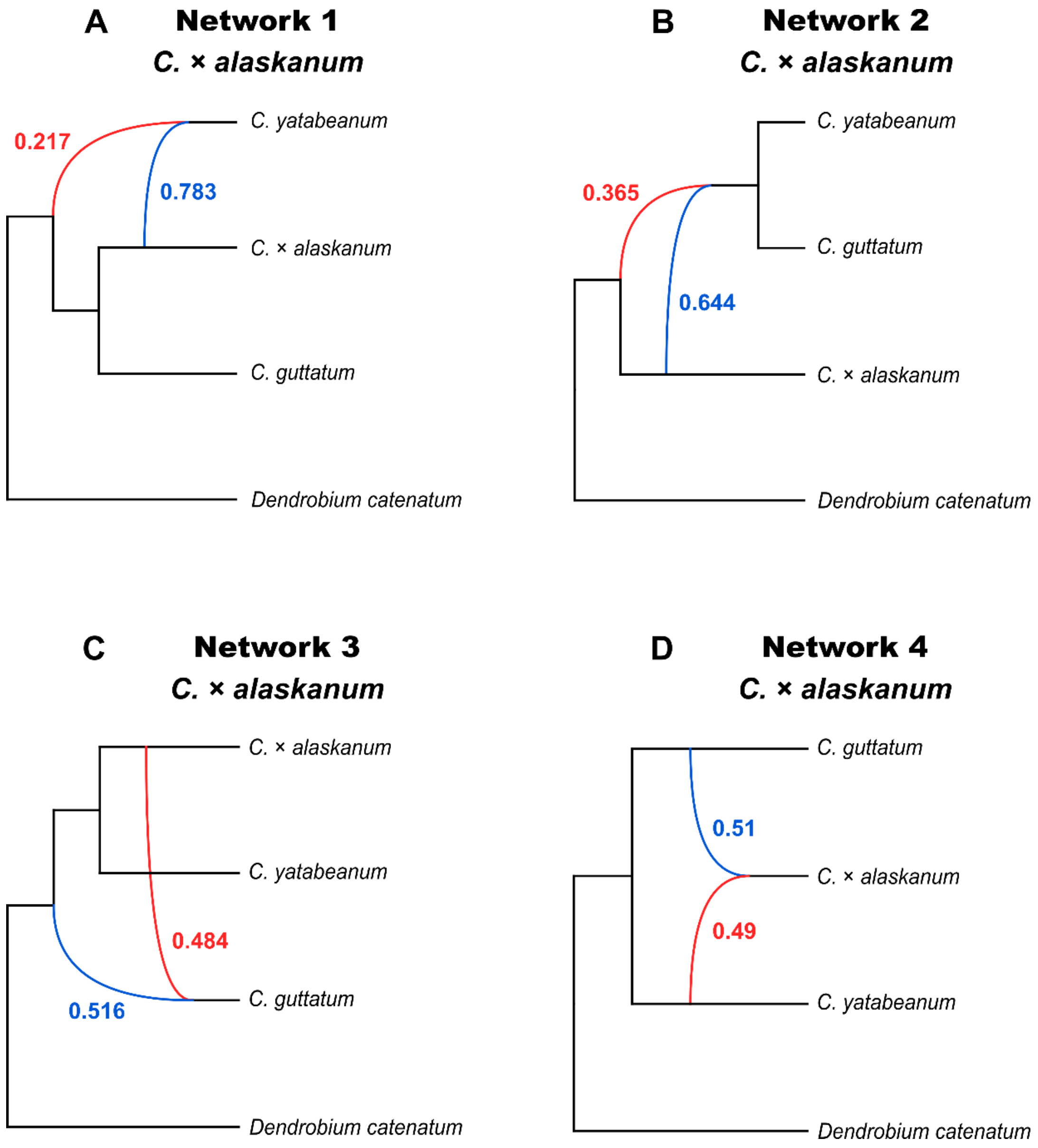

Figure S15: Phylogenetic networks including *C.* × *alaskanum* testing for one hybridization event (using the -po parameter) with the (A) first, (B) second, (C) third, and (D) fourth highest total log probability values. Based on the deltaAIC, deltaAICc, and deltaBIC scores listed in Supplementary Table S10, there is substantial evidence to support that the candidate models (B) and (C) are almost as good as the best model (A) (i.e., deltaAIC, deltaAICc, and deltaBIC < 2), while the support for candidate model (C) is comparatively lower (i.e., deltaAIC, deltaAICc, and deltaBIC = 2.04)

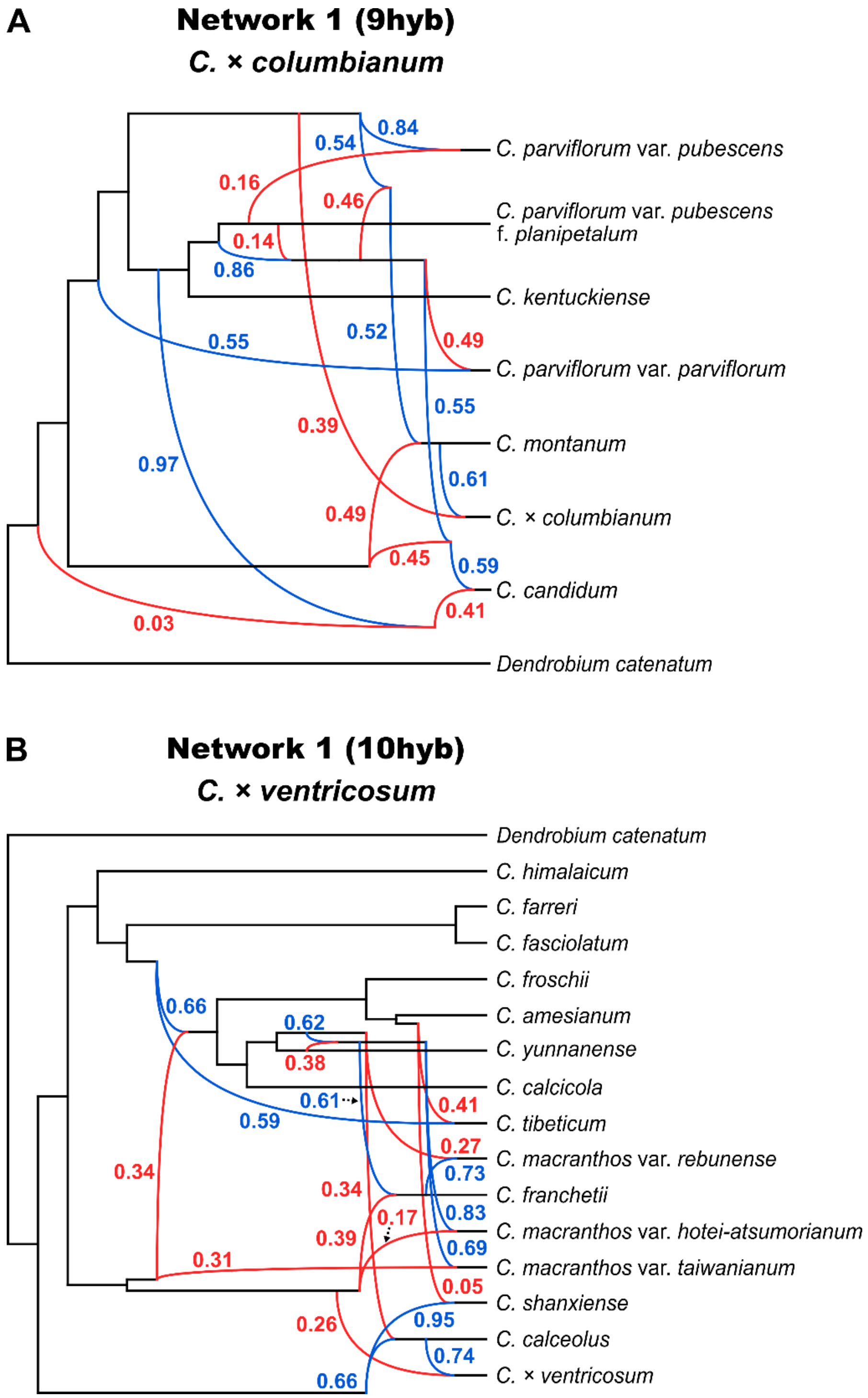

Figure S16: Phylogenetic networks with the overall highest total log probabilities testing for nine and ten hybridization events for the extracted subclades with the hybrids (A) *C.* × *columbianum* and (B) *C.* × *ventricosum*, respectively. The inheritance probabilities are shown for each parent hybrid edge (blue = major hybrid edge; red = minor hybrid edge).

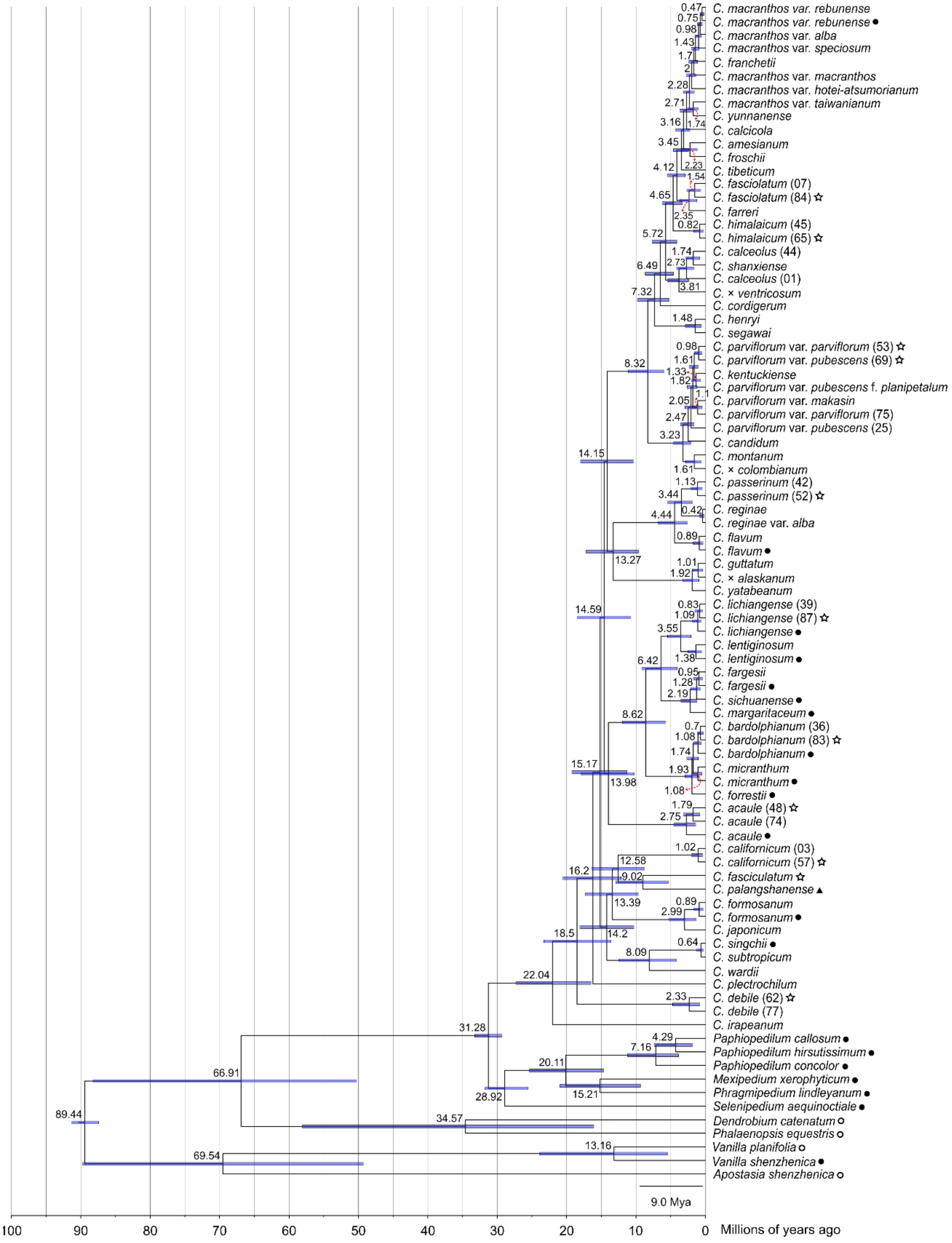

Figure S17: The maximum clade credibility tree of *Cypripedium* resulting from the molecular dating analysis with BEAST 2. The posterior median node height estimates (in Ma) are shown on the nodes, together with the 95% HPD bars. A time scale with the time before the present is shown at the bottom. Tip symbols: filled circles "●" denote transcriptomes, unfilled circles "○" denote genomes, filled triangles "▲" denote genome skimming sequences, and unfilled stars "☆" denote herbarium or old silica-dried specimens. Tips without symbols come from living specimens of the Botanical Collection at Oberhof.

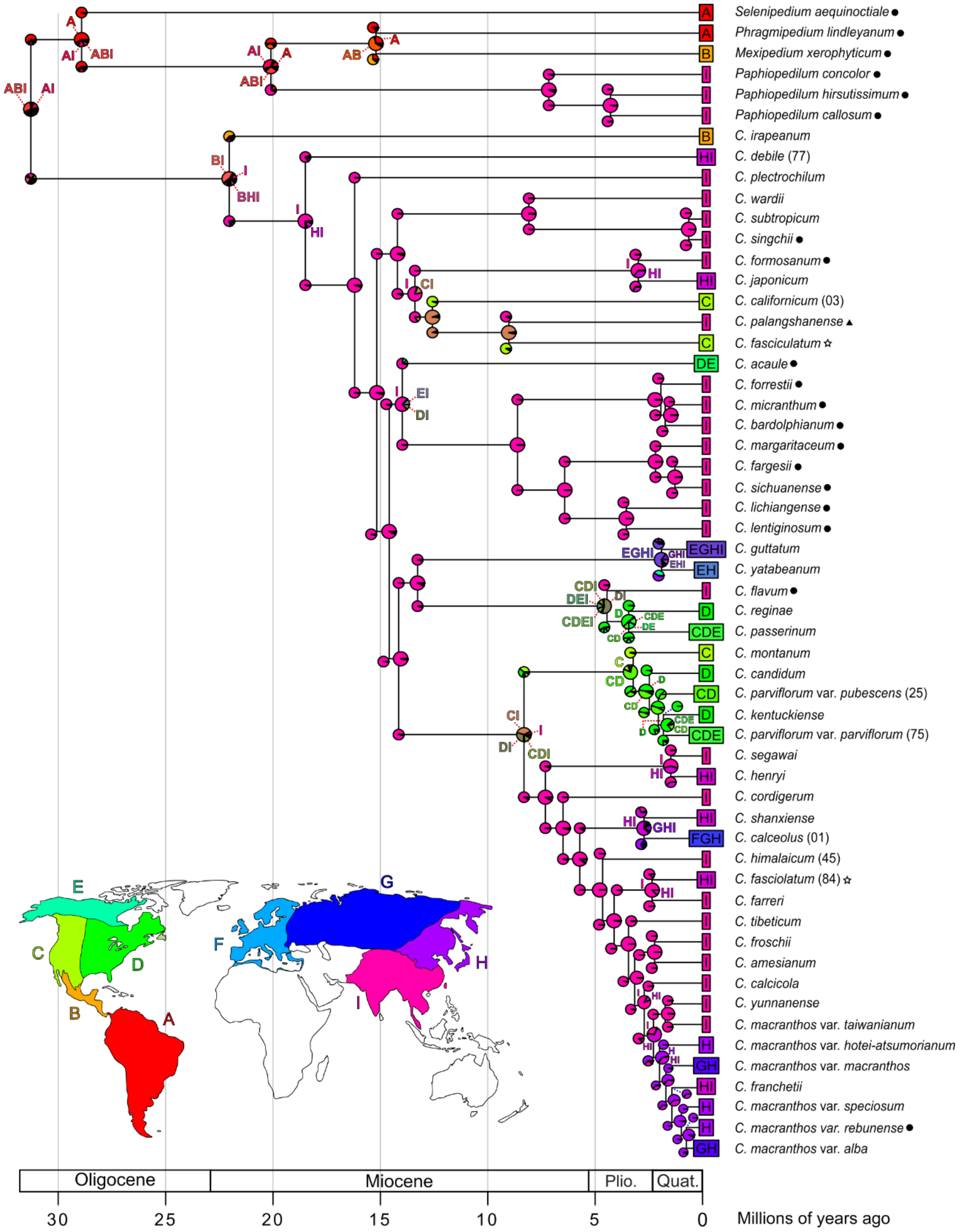

Figure S18: The relative proportions of the ancestral ranges estimated by BioGeoBEARS (nine area test, DEC model) and plotted as pie charts on the corresponding nodes of the dated maximum clade credibility tree of slipper orchids. Key to area codes: A = South America; B = Central America and Mexico; C = Western North America; D = Eastern North America; E = Northern North America; F = Eastern and Central Europe, the Mediterranean, and Scandinavia; G = Eastern Europe and Eurasia; H = Eastern Russia and Northeast Asia; I = Southeast Asia. Plio. = Pliocene; Quat. = Quaternary. Tip symbols: filled circles "●" denote transcriptomes, filled triangles "▲" denote genome skimming sequences, and unfilled stars "☆" denote herbarium or old silica-dried specimens. Tips without symbols come from living specimens of the Botanical Collection at Oberhof.

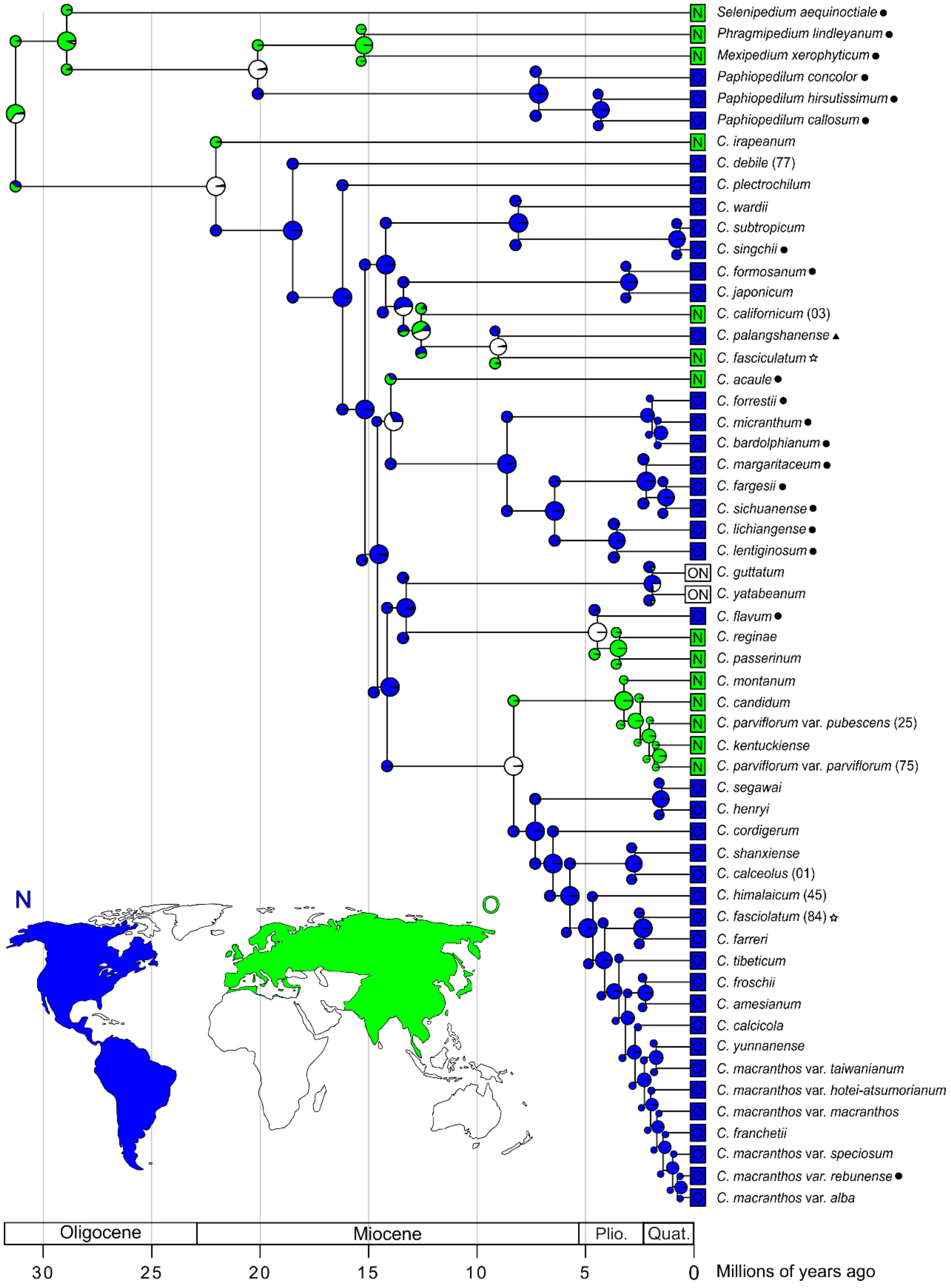

Figure S19: The relative proportions of the ancestral ranges estimated by BioGeoBEARS (two-area test, DIVALIKE model) and plotted as pie charts on the corresponding nodes of the dated maximum clade credibility tree of slipper orchids. Key to area codes: N = New World, O = Old World. Plio. = Pliocene; Quat. = Quaternary. Tip symbols: filled circles "●" denote transcriptomes, filled triangles "▲" denote genome skimming sequences, and unfilled stars "☆" denote herbarium or old silica-dried specimens. Tips without symbols come from living specimens of the Botanical Collection at Oberhof.

### Supplementary Text

*Library Preparation, Target Enrichment, and Sequencing*

**Methods S1**: Detailed library preparation and target enrichment protocol.

Total genomic DNA was isolated using the NucleoSpin Plant II kit: Genomic DNA from plants (Macherey-Nagel, Düren, Germany), following a modified version of the manufacturer's manual (Supplementary Data Table S5). Next, we quantified the concentration of all DNA samples using a Qubit 4 fluorometer with a Broad Range (BR) or High Sensitivity (HS) assay kit (Thermo Fisher Scientific, Waltham, Massachusetts, USA). After shearing the DNA to an average fragment size of 350 bp with a Covaris M220 Focused-ultrasonicator (Covaris, Woburn, Massachusetts, USA), we assessed the DNA fragment size distribution using a High Sensitivity DNA ScreenTape on a 4150 TapeStation System (Agilent Technologies, Santa Clara, California, USA).

For the preparation of the dual indexed libraries with the NEBNext Ultra II DNA Library Prep Kit for Illumina and the NEBNext Multiplex Oligos for Illumina (Dual Index Primers Set 1, New England Biolabs, Ipswich, Massachusetts, USA), we followed the recommended conditions of bead-based size selection according to distribution of DNA fragments per sample. We amplified the adaptor-ligated libraries in eight PCR cycles and measured DNA concentration with Qubit. The average fragment size of the libraries was assessed with the TapeStation. Prior to hybridization, the libraries were pooled in equal concentrations to include 250 ng of each library, with a maximum of 15 libraries per pooled library.

For the hybridization enrichment reaction, we combined the pooled libraries with the custom orchid-specific bait set Orchidaceae963 (Daicel Arbor Biosciences myBaits Target Capture Kit, Ann Arbor, MI, USA) and incubated at 60 °C (hybridization temperature, TH) for 16 hours overnight, following the Standard Protocol and the Blockers Mix setup designed for plants (myBaits Hybridization Capture for Targeted NGS, User Manual v. 5.02). The bead-based cleanup of the bait-target hybrids was performed at a wash temperature (TW) of 60 °C, and the hybridized libraries were subsequently amplified for 14 PCR cycles at 60 °C (annealing temperature, TA). Then, we purified the amplification reaction following the PCR clean-up protocol of the NucleoSpin Gel and PCR Clean-up kit (Macherey-Nagel, Düren, Germany). Finally, we checked the concentration and fragment size distribution of the libraries as before, using the Qubit and the TapeStation.

*Read Processing and Assembly*

**Methods S2**: Detailed read processing and assembly protocol for the target enrichment sequence data. To create the set of orchid genome and transcriptome references, original target exon sequences from the Orchidaceae963 bait set (https://github.com/laeserman/Orchidaceae963/blob/main/Orchidaceae963-targets.fa) were concatenated into 'genes' and used to identify the corresponding complete CDS from the *Phalaenopsis equestris* genome using BLAST. When no hits were produced, we used the genome of *Dendrobium catenatum* instead. In the end, 950 out of the original 963 genes were extracted. Then, we used the raw transcriptome assembly from 17 orchids (Supplementary Data Table S4), consisting of *Vanilla shenzhenica* and 16 species of slipper orchids to extend the genome references. RNAseq data processing and transcriptome assembly followed Morales-Briones *et al.* (2021). We used CAPTUS v.0.9.90 (Ortiz *et al.* 2023) to extract the corresponding loci from the 17 transcriptomes and the genomes of *Apostasia shenzhenica*, *D. catenatum*, *P. equestris*, and *Vanilla planifolia*. The extracted loci were used as the extended reference dataset for loci extraction in our own generated target enrichment data of *Cypripedium*.

We checked the quality of the raw reads using FastQC v.0.11.9 (Andrews, 2010) and MultiQC v.1.14 (Ewels *et al.,* 2016). PCR duplicates were removed with ParDRe v.2.1.5 (González-Domínguez and Schmidt, 2016). Then, using CAPTUS v.1.0.0 we trimmed the sequencing adaptors and low-quality bases, assembled the reads, and extracted the nuclear loci based on the reference dataset, setting the minimum contig depth to eight for the assembly step and the minimum percentages of identity and coverage to 75% and 50%, respectively, for the extraction step, to decrease the retention of contigs resulting from potential erroneous reads. As mentioned in the main text, the extracted loci were combined with loci extracted from the genomes and transcriptomes mentioned above, and the coding sequences of the combined dataset were extracted in FASTA files while keeping up to 25 paralog copies per sample.

*Orthology Inference of nuclear loci*

**Methods S3**: Detailed orthology inference protocol.

To infer orthologs for the phylogenetic reconstruction, we followed a modified version of the methods described in Morales-Briones *et al.* (2022; https://bitbucket.org/dfmoralesb/target_enrichment_orthology). First, we aligned each locus using the OMM_MACSE pipeline v.12.01 (Scornavacca *et al.*, 2019), which pre-filters non-homologous sequence fragments with HMMCleaner (Di Franco *et al.*, 2019), performs the multi-sequence alignment with MAFFT v.7.271, and carries out the translation accounting for frameshifts using MACSE v.2.08 (Ranwez *et al.,* 2018). OMM_MACSE was set to replace codons containing frameshifts with gaps. Next, we used Phyx ('pxclsq' Brown *et al.,* 2017) to remove aligned columns with more than 90% missing data. We inferred maximum likelihood (ML) homolog gene trees with IQ-TREE v.2.0.7 (Minh *et al.,* 2020) using extended model selection (Kalyaanamoorthy *et al.,* 2017) and no clade support. Then, we masked mono- and paraphyletic tips that belong to the same taxon, keeping the tips with the most unambiguous characters in the trimmed loci alignments for each taxon as described in (Yang and Smith, 2014). Spurious tips with unusually long branches were removed by reducing the tree diameter with TreeShrink v.1.3.9 (Mai and Mirarab, 2018). We ran TreeShrink with the ‘per-gene’ mode, a false positive error rate threshold (α) of 0.05, and excluding the outgroups. We wrote FASTA files from the output homolog trees and followed the same steps as for the output FASTA files from CAPTUS, aligning them with OMM_MACSE and removing aligned columns with >90% missing data using Phyx. To infer the final homolog gene trees, we used IQ-TREE with extended model selection and assessed the clade support with 1,000 ultrafast bootstrap (BS) replicates. Following orthology inference using the tree-based “monophyletic outgroup” (MO) approach, as described in the main text, we wrote FASTA files from the output ortholog trees, re-aligned the loci using OMM_MACSE, and cleaned the alignments with Phyx, as before.

*Testing for Hybridization Events*

**Methods S4**: Detailed methods regarding the phylogenetic network analyses for the test investigating intra-sectional hybridization within the subclades containing the three described hybrids that were included in this study, their putative parent taxa, and other taxa that share the same MRCA.

To test whether the hybrid status of the three described hybrids included in our taxon sampling is supported by our target enrichment data, we followed a similar approach to the one described for investigating potential hybridization at the backbone of the *Cypripedium* phylogeny in the main text. However, this time, we extracted the three subclades containing each hybrid, along with the putative parent taxa and other taxa sharing their MRCA. We reduced computational load by removing duplicated taxa, leaving a single representative for each monophyletic taxon. In the case of paraphyletic taxa, one representative taxon was left from each conspecific monophyletic subclade or from a group of consecutively diverging conspecific varieties. The number of maximum reticulation events and the number of optimal output networks were set to one and ten, respectively, for all three tests. The option “po” was specified to optimize the branch lengths and inheritance probabilities under full likelihood for the inferred *C.* × *alaskanum* networks, allowing for the direct comparison of all output networks with the Akaike's Information Criterion (AIC), the corrected Akaike's Information Criterion (AICc), and the Bayesian information criterion (BIC) scores calculated according to Yu *et al.* (2012). This optimization was only performed for the *C.* × *alaskanum* networks, which contain only four taxa, as it gets more time-consuming with an increasing number of taxa.

*Ancestral Range Estimation*

**Methods S5**: Division of nine areas for the biogeographical analyses in BioGeoBEARS.

The areas were divided based on the current distribution of the taxa included in the analysis, their proximity, and their distinct floristic and topoclimatic characteristics (e.g., climate, precipitation, elevation). South America (area A) was specified as a large distinct area since only a few outgroup slipper orchids are restricted to its Northern part [i.e., *Selenipedium aequinoctiale* and *Phragmipedium lindleyanum*; POWO, 2023]. We separated Central America and Mexico (area B, containing section Irapeana) from South America at the Isthmus of Panama and from North America at the deserts to its north (i.e., Baja, Mojave, Sonoran, and Chihuahuan deserts). North America was divided into three areas: the Northern area E (colder to polar, humid climate), the Eastern area D (lower altitudes, higher humidity and precipitation than the Western area), and the Western area C (higher altitudes, lower humidity and precipitation than Eastern area(Kottek *et al.* 2006; Xiao *et al.* 2020). All three areas match the distributions of different *Cypripedium* species, with only five species found in area E, while the Great Plains in the middle of North America seemingly create a distribution boundary for multiple *Cypripedium* species (Supplementary Data Fig. S1). The Mediterranean and Scandinavian regions were grouped with Western and Central Europe (area F) because only *C. calceolus* occurs in all three areas (Eccarius, 2009; Frosch and Cribb, 2012; Chen *et al.,* 2013; Walid *et al.,* 2019). We split area F from Eastern Europe and Russia (area G) at the boundaries of the Sarmatic and Pontic-South Siberian floristic provinces according to Schroeder (1998), as they have a more continental climate and match the limit of *C. guttatum's* distribution in the European continent (Pfadenhauer and Klötzli, 2020; Supplementary Data Fig. S1). Area G was separated from the two Asiatic areas (namely, the Southeast Asian area I and the Northeast Asian area H) due to the higher number of unique *Cypripedium* species occurring there, as well as their different climates and floristic provinces (Kottek *et al.*, 2006; Fridley, 2008). The Southeast and Northeast Asian areas are split around the Qinling Mountains–Huaihe River Line (aka Qinling–Huaihe line), a natural topographic boundary that separates North temperate from South tropical China (Hu *et al.*, 2020), which also seems to create a boundary for the distribution of several *Cypripedium* species. To reduce the state space and thus the computational time of the analysis, Taiwan was included in the same area as Southeast Asia (area I), and Japan in the same area as Northeast Asia and Eastern Russia (area H) due to their proximity.

**Methods S6**: Input tree and taxon distribution information for the biogeographical analyses in BioGeoBEARS.

To prepare the input tree for the biogeographic analyses with BioGeoBEARS, we removed hybrids and duplicated species from the time-calibrated maximum clade credibility tree obtained from the divergence time estimation analysis. When multiple varieties were present, a single specimen per each accepted variety (according to Frosch and Cribb, 2012) was kept due to distinct distributions. *Cypripedium amesianum* and the ambiguous *C. macranthos* var. *alba* were also kept since they were not monophyletic with their presumably synonymous taxa [i.e., *C. yunnanense* and *C. macranthos* var. *albiflorum* (now synonym of *C. macranthos* var. *macranthos*), respectively], and, therefore, considered distinct taxonomic units for this analysis. We also removed non-slipper orchid taxa because of scarce sampling.

The taxon distributions were based on Eccarius (2009), Frosch and Cribb (2012), Chen *et al.* (2013), and (Walid *et al.* 2019). The distributions of *C. amesianum* and *C. macranthos* var. *alba* were considered the same as their synonyms according to Frosch and Cribb (2012; Supplementary Data Figure S1).

*Inferred Nuclear Species Phylogeny and Discordance*

**Results S1:** Species-level results of the *Cypripedium* phylogeny inferred based on target enrichment data of 913 nuclear loci.

Regarding the monophyly at the species level, two species within sect. *Cypripedium* that included numerous infraspecific taxa (i.e., varieties and one form), *C. macranthos* and *C. parviflorum*, were consistently recovered as paraphyletic in both phylogenies, with *C. franchetii* and *C. yunnanense* nested in the former, and *C. kentuckiense* nested in the latter. Two pairs of synonyms (according to (Frosch and Cribb 2012) formed monophyletic clades: that is, (*C. parviflorum* var. *makasin*, *C. parviflorum* var. *parviflorum*), and (*C. subtropicum*, *C. singchii*). However, the former pair was only formed between specimens from silica-gel material, while *C. parviflorum* specimens from herbarium material created a separate clade. In contrast, *C. amesianum* was more closely related to *C. froschii* rather than its synonymous species, *C. yunnanense*. Additionally, the ambiguous taxon *C. macranthos* var. *alba*, which was presumed to be either *C. macranthos* var. *albiflorum* (now a synonym of *C. macranthos* var. *macranthos*) or *C. macranthos* var. *album*, was more closely related to the equally white-flowered *C. macranthos* var. *rebunense*. Regarding the rest of the species, ASTRAL recovered most as monophyletic, grouping all conspecific specimens retrieved from different sources (i.e., Botanical Collection at Oberhof, herbarium M, and SRA), except the paraphyletic *C. calceolus* and *C. fargesii*. Notably, the three included hybrids following Frosch and Cribb (2012) were placed in the same clades as one or both of their putative parent taxa.

*Hybridization Networks*

**Results S2:** Detailed phylogenetic network analysis results for the test investigating intra-sectional hybridization within the subclades containing the three described hybrids that were included in this study, their putative parent taxa, and other taxa that share the same MRCA.

To address our questions about whether the three known hybrid species included in this study are supported as products of hybridization between their putative parent taxa by our target enrichment data, we plotted the most likely phylogenetic networks that tested for one reticulation event. Regarding the test with *C.* × *alaskanum*, although the most likely network (Network 1, total log probability = ~ -556.90) indicated that *C. yatabeanum* is a hybrid between an unsampled taxon sharing an MRCA with sect. *Bifolia* (γ = 0.22) and *C.* × *alaskanum* (γ = 0.78), model selection comparing all networks resulting from this test suggested that the next three networks with the highest probability (total log probabilities for Networks 2: ~ -557.48, Network 3: ~ 557.64, and Network 4: ~ -557.92) were almost as good as the best model (deltaAIC, detlaAICc, and deltaBIC ≲ 2; Supplementary Data Fig. S15 and Table S10). Each of the latter networks provided support for different taxa being hybrids resulting from different crosses; therefore, it is not possible to infer accurate conclusions regarding the hybrid status of *C.* × *alaskanum* based on the PhyloNet analysis of our nuclear target enrichment data.

The most likely hybridization network of *C.* × *columbianum* (total log probability = ~ -16,825.20) supports that the clade containing the *C. parviflorum* varieties and *C. kentuckiense* was created following a hybridization between *C.* × *columbianum* and *C. candidum* (γ = 0.16 and 0.84, respectively; Supplementary Data Fig. S14 A). As for the network that includes *C.* × *ventricosum* (total log probability = ~ -72,700.35), the results indicate that both *C.* × *ventricosum* and *C. calceolus* are sister taxa that arose from a hybridization event between *C. macranthos* var. *rebunense* (γ = 0.38) and *C. shanxiense* (γ = 0.62; Supplementary Data Fig. S14 B). It is worth noting that when looking at the total log probabilities of all analyses testing for one to ten reticulation events in both subclades, the overall most likely models indicated a higher number of hybridizations; namely, 9 events in the *C.* × *columbianum* subclade (total log probability = ~ -72,263.36), and 10 events in the *C.* × *ventricosum* subclade (total log probability = ~ -16,778.53; Supplementary Data Fig. S13 and S16). However, since model selection was not performed for these analyses, further investigation is required to assess and validate these results.

*Rapid Radiation and Hybridization Promoted Diversification*

**Discussion S1:** Detailed discussion of the phylogenetic network analysis results for the test investigating intra-sectional hybridization within the subclades containing the three described hybrids that were included in this study, their putative parent taxa, and other taxa that share the same MRCA. Regarding reticulation at the intra-sectional level, our analyses indicated multiple potential reticulation events within sections *Bifolia*, *Cypripedium*, and *Subtropica*. Most nodes leading to the hybrids identified through our analyses had decreased concordance levels, including the MRCA node of the (*C. guttatum*, *C.* × *alaskanum*) clade. Although *C.* × *alaskanum* has been previously described as a natural hybrid between *C. yatabeanum* and *C. guttatum* by Brown (1995), it was difficult to assess its validity because no analysis accompanied the description (Cribb, 1997). Despite the hybrid status of *C.* × *alaskanum*, our target enrichment data suggested that *C. yatabeanum* is a hybrid between *C.* × *alaskanum* and an unsampled taxon closely related to the (*C. guttatum, C.* × *alaskanum*) clade. However, the support for this model was not significantly higher than that of other output networks, indicating different hybridization events. Therefore, additional analyses and molecular data are crucial to obtain robust molecular evidence for the hybrid status of *C.* × *alaskanum.*

As for the PhyloNet analyses within sect. *Cypripedium*, testing for a maximum of one hybridization event also yielded some unexpected results. Sheviak (1992) proposed that *C.* × *columbianum* is a hybrid between *C. montanum* and *C. parviflorum* var. *pubescens* based on an extensive survey and analysis of wild and herbarium specimens, with hybrids having intermediate morphological characteristics between the presumed parent taxa. The phylogenetic network containing *C.* × *columbianum* disagreed with Sheviak's taxon description, identifying the *C. parviflorum* complex clade as a product of a hybrid cross between *C.* × *columbianum* and *C. candidum*, although their current ranges do not overlap in North America.

The results of a similar test on the dataset containing *C.* × *ventricosum* supported that the taxon did not constitute a hybrid from a cross between *C. calceolus* and *C. macranthos* as it was previously suggested based on the resemblance of its flower with an artificial hybrid between the presumed parent taxa (Rolfe, 1904, 1910; Cribb, 1997). Instead, the results indicated that *C.* × *ventricosum* and *C. calceolus* are products of a hybridization event between *C. shanxiense* and *C. macranthos.* Another hybrid, *C.* × *catherinae*, which occurs in Far East Russia (Siberia) and possibly in Korea and Northeast China, where the parent taxa's distributions overlap, was already described from the hybridization between these two taxa (Frosch and Cribb, 2012; Chen *et al.*, 2013). *Cypripedium calceolus* and *C.* × *ventricosum* can also be found sympatrically with the parent taxa identified in our analysis, namely, in Siberia and Northeastern China, while *C. calceolus* is also found in Japan and on Sakhalin Island (Cribb, 1997; Frosch and Cribb, 2012; Chen *et al.*, 2013). Interestingly, *C.* × *ventricosum* and *C. calceolus* formed a clade with *C. shanxiense*—the putative parent taxon corresponding to the major edge ($\gamma = 0.62$)—in our target enrichment phylogenies, while in the chloroplast phylogeny, *C.* × *ventricosum* and one of the three *C. calceolus* specimens were more closely related to *C. macranthos* varieties. Moreover, a sister relationship is consistently recovered between *C. calceolus* and *C. shanxiense* (Fatihah *et al.*, 2011; Li *et al.*, 2011; Liu *et al.*, 2021; Szlachetko *et al.*, 2021) or between *C. calceolus* and *C. macranthos* in the absence of *C. shanxiense*

(Cox *et al.,* 1997). Additionally, *C.* × *ventricosum* and *C. macranthos* formed a clade closely related to the (*C. calceolus*, *C. shanxiense*) clade in the phylogeny of Liu *et al.* (2021).

In support of Rolfe's description, a statistical analysis of morphological and allozyme data from *C.* × *ventricosum* corroborated its status as a hybrid between *C. calceolus* and *C. macranthos* (Knyasev *et al.,* 2000). However, the fact that *C. calceolus* and *C.* × *ventricosum* may share the same parent taxa based on our results (i.e., *C. shanxiense* and *C. macranthos*) and that introgressive hybridization has been previously reported between *C. calceolus* and *C.* × *ventricosum* (Knyasev *et al.,* 2000) could elucidate the relationship between these species. Furthermore, it has been shown that the genetic structure of *C. shanxiense* was similar to *C. calceolus* from the eastern part of the range where they are sympatric, with the authors arguing that it may have been a result of introgressive hybridization, making the classification of these taxa even more challenging (Filippov and Andronova, 2011). At the same time, a hybrid has also been described between these two species (i.e., *C.* × *microsaccos*; Frosch and Cribb, 2012; Chen *et al.*, 2013).

When looking at the overall most probable phylogenetic networks for the subclades including *C.* × *ventricosum* and *C.* × *columbianum*, the results suggested that extensive hybridization events between sampled and unsampled taxa occurred within both subclades of sect. *Cypripedium*, which is supported by the evidence of pervasive hybridization within the genus, as mentioned before (Klier *et al.,* 1991; Hu *et al.,* 2011; Frosch and Cribb, 2012; Szlachetko *et al.,* 2017; Pupulin and Díaz-Morales, 2018). The mixed genetic and morphological signals created by these hybridizations could have significantly contributed to the difficulty in specific delimitation and taxonomic classification within the sect. *Cypripedium*, with the same taxa receiving a species, subspecies, variety, or hybrid rank by different taxonomists (e.g., *C. froschii*).

However, due to a high taxon sampling in most of our PhyloNet analyses except the one containing *C.* × *alaskanum*, we did not optimize the branch lengths and inheritance probabilities under full likelihood, as it becomes more computationally intensive and time-consuming with an increasing number of taxa. Consequently, we could not perform model selection to properly compare the network with the highest probability to other networks with lower probabilities inferred in these tests. Therefore, it is crucial to carry out additional studies and in-depth taxonomic revisions including further lines of evidence (e.g., a higher number and variety of molecular markers, allozyme markers, genetic studies at the population level, geometric morphometric studies of homologous morphological traits, etc.) to thoroughly assess the validity of the hybridization events that we identified in the genus *Cypripedium* through our investigation.

LITERATURE CITED

**Andrews S**. **2010**. FastQC: A Quality Control Tool for High Throughput Sequence Data [Online].

**Brown PM**. **1995**. New taxa and taxonomic notes. **1**: 195–200.

**Brown JW, Walker JF, Smith SA**. **2017**. Phyx: phylogenetic tools for unix. *Bioinformatics* **33**: 1886–1888.

**Cai J, Liu X, Vanneste K, *et al.* 2015**. The genome sequence of the orchid Phalaenopsis equestris. *Nature Genetics* **47**: 65–72.

**Chen SC, Liu ZJ, Chen LJ, Li LQ**. **2013**. *The Genus Cypripedium in China*. Peking: Science Press.

**Cox AV, Pridgeon AM, Albert VA, Chase MW**. **1997**. Phylogenetics of the slipper orchids (Cypripedioideae, Orchidaceae): Nuclear rDNA ITS sequences. *Plant Systematics and Evolution* **208**: 197–223.

**Cribb P**. **1997**. *The Genus Cypripedium*. Portland: Timber Press.

**Di Franco A, Poujol R, Baurain D, Philippe H**. **2019**. Evaluating the usefulness of alignment filtering methods to reduce the impact of errors on evolutionary inferences. *BMC Evolutionary Biology* **19**: 21.

**Eccarius W**. **2009**. *Orchideengattung Cypripedium*. EchinoMedia.

**Ewels P, Magnusson M, Lundin S, Käller M**. **2016**. MultiQC: summarize analysis results for multiple tools and samples in a single report. *Bioinformatics* **32**: 3047–3048.

**Fatihah HN, Fay M, Maxted N**. **2011**. Molecular Phylogenetics of Cypripedium L. (Cypripedioideae: Orchidaceae) Based on Plastid and Nuclear DNA Sequences. *Journal of Agrobiotechnology* **2**: 111–118.

**Filippov E, Andronova E**. **2011**. Genetic differentiation in plants of the genus Cypripedium from Russia inferred from allozyme data. *Genetika* **47**: 615–23.

**Fridley J**. **2008**. Of Asian Forests and European Fields: Eastern U.S. Plant Invasions in a Global Floristic Context. *PloS one* **3**: e3630.

**Frosch W, Cribb P**. **2012**. *Hardy Cypripedium: Species, hybrids and cultivation*. Kew Publishing Kew.

**González-Domínguez J, Schmidt B**. **2016**. ParDRe: faster parallel duplicated reads removal tool for sequencing studies. *Bioinformatics* **32**: 1562–1564.

**Hu S-J, Hu H, Yan N, Huang J-L, Li S-Y**. **2011**. Hybridization and asymmetric introgression between Cypripedium tibeticum and C. yunnanense in Shangrila County, Yunnan Province, China. *Nordic Journal of Botany* **29**: 625–631.

**Hu C, Jiao Z, Deng X, *et al.*** **2022**. The ecological adaptation of the unparalleled plastome character evolution in slipper orchids. *Frontiers in Plant Science* **13**: 1075098.

**Hu Y, Yao Y, Kou Z**. **2020**. Exploring on the climate regionalization of Qinling-Daba mountains based on Geodetector-SVM model. *PLoS ONE* **15**: e0241047.

**Kalyaanamoorthy S, Minh BQ, Wong TKF, von Haeseler A, Jermiin LS**. **2017**. ModelFinder: fast model selection for accurate phylogenetic estimates. *Nature Methods* **14**: 587–589.

**Klier K, Leoschke MJ, Wendel JF**. **1991**. Hybridization and Introgression in White and Yellow Ladyslipper Orchids (Cypripedium candidum and C. pubescens). *Journal of Heredity* **82**: 305–318.

**Knyasev MS, Kulikov PV, Knyaseva OI, Semerikov VL**. **2000**. Interspecific hybridization in northern Eurasian Cypripedium: morphometric and genetic evidence of the hybrid origin of C. ventricosum. *Lindleyana* **15**: 10–20.

**Kottek M, Grieser J, Beck C, Rudolf B, Rubel F**. **2006**. World Map of the Köppen-Geiger Climate Classification Updated. *Meteorologische Zeitschrift* **15**: 259–263.

**Li J, Liu Z, Salazar GA, *et al.*** **2011**. Molecular phylogeny of Cypripedium (Orchidaceae: Cypripedioideae) inferred from multiple nuclear and chloroplast regions. *Molecular Phylogenetics and Evolution* **61**: 308–320.

**Lindley J**. **1840**. *The genera and species of orchidaceous plants*. London: Ridgways.

**Liu H, Jacquemyn H, Chen W, *et al.* 2021**. Niche evolution and historical biogeography of lady slipper orchids in North America and Eurasia. *Journal of Biogeography* **48**: 2727–2741.

**Mai U, Mirarab S**. **2018**. TreeShrink: fast and accurate detection of outlier long branches in collections of phylogenetic trees. *BMC genomics* **19**: 23–40.

**Minh BQ, Schmidt HA, Chernomor O, *et al.* 2020**. IQ-TREE 2: New Models and Efficient Methods for Phylogenetic Inference in the Genomic Era. *Molecular Biology and Evolution* **37**: 1530–1534.

**Morales-Briones DF, Gehrke B, Huang C-H, *et al.* 2022**. Analysis of paralogs in target enrichment data pinpoints multiple ancient polyploidy events in Alchemilla sl (Rosaceae). *Systematic Biology* **71**: 190–207.

**Morales-Briones DF, Kadereit G, Tefarikis DT, *et al.* 2021**. Disentangling Sources of Gene Tree Discordance in Phylogenomic Data Sets: Testing Ancient Hybridizations in Amaranthaceae s.l. *Systematic Biology* **70**: 219–235.

**Ortiz EM, Höwener A, Shigita G, *et al.* 2023**. A novel phylogenomics pipeline reveals complex pattern of reticulate evolution in Cucurbitales. : 2023.10.27.564367.

**Pease JB, Brown JW, Walker JF, Hinchliff CE, Smith SA**. **2018**. Quartet Sampling distinguishes lack of support from conflicting support in the green plant tree of life. *American journal of botany* **105**: 385–403.

**Pfadenhauer JS, Klötzli FA**. **2020**. Fundamentals towards Understanding Global Vegetation In: Pfadenhauer JS, Klötzli FA, eds. *Global Vegetation: Fundamentals, Ecology and Distribution*. Cham: Springer International Publishing, 1–120.

**Pfitzer EHH**. **1903**. *Orchidaceae–Pleonandrae*. Leipzig: Engelmann.

**Piet Q, Droc G, Marande W, *et al.* 2022**. A chromosome-level, haplotype-phased Vanilla planifolia genome highlights the challenge of partial endoreplication for accurate whole-genome assembly. *Plant Communications* **3**: 100330.

**POWO**. **2023**. *Plants of the World Online. Facilitated by the Royal Botanic Gardens, Kew. Published on the Internet.* http://www.plantsoftheworldonline.org/. 12 Feb. 2023.

**Pupulin F, Díaz-Morales M**. **2018**. On the meaning of Cypripedium × grande (Orchidaceae) and its taxonomic history, with a new name for the nothospecies occurring in Costa Rica and Panama. *Phytotaxa* **382**: 167.

**Ranwez V, Douzery EJ, Cambon C, Chantret N, Delsuc F**. **2018**. MACSE v2: toolkit for the alignment of coding sequences accounting for frameshifts and stop codons. *Molecular biology and evolution* **35**: 2582–2584.

**Rolfe RA**. **1904**. Cypripedium calceolus × macranthos. **12**: 185.

**Rolfe RA**. **1910**. Cypripedium × ventricosum. **18**: 215.

**Schroeder FG**. **1998**. *Lehrbuch der Pflanzengeographie*. Quelle & Meyer.

**Scornavacca C, Belkhir K, Lopez J, *et al.*** **2019**. OrthoMaM v10: Scaling-Up Orthologous Coding Sequence and Exon Alignments with More than One Hundred Mammalian Genomes. *Molecular Biology and Evolution* **36**: 861–862.

**Sheviak C**. **1992**. Natural hybridisation between Cypripedium montanum and its yellow-lipped relatives. **61**: 558.

**Szlachetko DL, Górniak M, Kowalkowska AK, Kolanowska M, Jurczak-Kurek A, Archila Morales F**. **2021**. The natural history of the genus Cypripedium (Orchidaceae). *Plant Biosystems-An International Journal Dealing with all Aspects of Plant Biology* **155**: 772–796.

**Szlachetko DL, Kolanowska M, Muller F, Vannini J, Rojek J, Górniak M**. **2017**. First Guatemalan record of natural hybridisation between Neotropical species of the lady's slipper orchid (Orchidaceae, Cypripedioideae). *PeerJ* **5**: e4162.

**Walid N, Rebbas K, Krouchi F**. **2019**. Découverte de Cypripedium calceolus (Orchidaceae) au Djurdjura (Algérie), nouvelle pour l'Afrique du Nord. *Flora Mediterranea* **29**: 207–214.

**Xiao X, Liang S, He T, Wu D, Pei C, Gong J**. **2020**. *Estimating fractional snow cover from passive microwave brightness temperature data using MODIS snow cover product over North America*.

**Yang Y, Smith SA**. **2014**. Orthology inference in nonmodel organisms using transcriptomes and low-coverage genomes: improving accuracy and matrix occupancy for phylogenomics. *Molecular biology and evolution* **31**: 3081–3092.

**Yu Y, Degnan JH, Nakhleh L**. **2012**. The Probability of a Gene Tree Topology within a Phylogenetic Network with Applications to Hybridization Detection. *PLOS Genetics* **8**: e1002660.

**Zhang G-Q, Liu K-W, Li Z, *et al.*** **2017**. The Apostasia genome and the evolution of orchids. *Nature* **549**: 379–383.

**Zhang G-Q, Xu Q, Bian C, *et al.*** **2016**. The Dendrobium catenatum Lindl. genome sequence provides insights into polysaccharide synthase, floral development and adaptive evolution. *Scientific Reports* **6**: 19029.
